# Interspecies transfer of giant virulence-factor-like proteins in a bacterial symbiosis

**DOI:** 10.64898/2026.03.19.712371

**Authors:** Steven B. Kuzyk, Petra Henke, Anika Methner, Tina Rietschel, Franziska Burkart, Mathias Müsken, Meina Neumann-Schaal, Gerhard Wanner, Jörg Overmann

## Abstract

The transfer of virulence factors into eukaryotic cells is a hallmark of bacterial pathogenesis. We report the expression, interspecies transfer, subcellular localization, and potential functions of three unusually large virulence factor-like proteins that underlie a bipartite mutualistic bacterial symbiosis. These proteins are synthesized by green sulfur bacterial epibionts surrounding a central motile chemoheterotroph in the multicellular phototrophic consortium ‘*Chlorochromatium aggregatum*’. While symbiosis-proteins remain intracellular during axenic epibiont growth, they are transferred to the partner bacterium in the association. An RTX-like protein secreted towards the central bacterium is capable of degrading its alginate capsule, thereby promoting direct cell-to-cell contact. Two gigantic hemagglutinin-like proteins are predicted to fold when binding extracellular Ca^2+^ to form Type 6-like auto injection needles, explaining their observed transfer into the central bacterium. These functionalities extend far beyond the known pathogenic interactions of bacteria with eukaryotes and provide new perspectives on the evolution of bacterial virulence factors.

## Introduction

Prokaryotes have established multifarious symbioses with a broad diversity of plants, animals and protists. While several thousand bacterial pathogens of vertebrates have been documented [**1**], the number of non-redundant ribosomal sequence types attributed to mutualists is as high as that of pathogenic microorganisms (1799 versus 1778, respectively; [**2**]). Yet, the currently described species of bacterial pathogens still outnumber those of mutualists [**3**], with the molecular mechanisms underlying mutualistic associations and their evolutionary links to pathogenicity often remaining enigmatic [**4,5**]. This is particularly true for symbiotic interactions between different species of prokaryotes.

Prokaryotic consortia represent the most advanced symbiotic associations known to date, and involve two or more species maintaining direct cellular contact [**6**], a relationship coined ‘heterologous multicellularity’ [**7**]. They mediate key biogeochemical transformations in stratified aquatic lakes and sediments [**4**], intestinal microbiomes [**8**], and wastewater treatment plants [**9**]. The phototrophic consortium ‘*Chlorochromatium aggregatum*’ (Fig. 1a) so far represents the only bacterial consortium that can be cultivated in an intact state and grown to high densities in the laboratory [**10**]. It consists of up to 24 epibiotic green sulfur bacteria (*Chlorobium chlorochromatii*) surrounding a central, motile betaproteobacterium ‘*Candidatus* Symbiobacter mobilis’ (Fig. 1b,e). Both partners are highly adapted to this mutualistic interaction as indicated by several features: **(i)** The highly coordinated division of all cells which results in the formation of two intact daughter consortia (Fig. 1d; [**11**]), **(ii)** the absence of free-living epibionts or central bacteria in natural environments [**12,13**], **(iii)** an intracellular sorting of photosynthetic antenna in epibiont cells (Fig. 1f; [**14**]) together with a specialized subcellular architecture of cell contact sites (Fig. 1c,f,g; [**15**]), **(iv)** a reciprocal interdependence of metabolic activity between partners [**16**], **(v)** mutualistic chemotactic and scotophobic responses of the symbiotic central bacterium which carry the immotile epibionts to suitable habitats containing sulfide and light required for their anoxygenic photosynthesis [**10,16,17**], and, finally, **(vi)** the reduced genome content of central bacterium ‘*Ca.* S. mobilis’ that likely renders it obligately dependent on its partner [**18**]. ‘*C. aggregatum*’ thus constitutes a prime model system for studying the molecular basis of bacterial heterologous multicellularity [**7,19**].

**Fig. 1.**
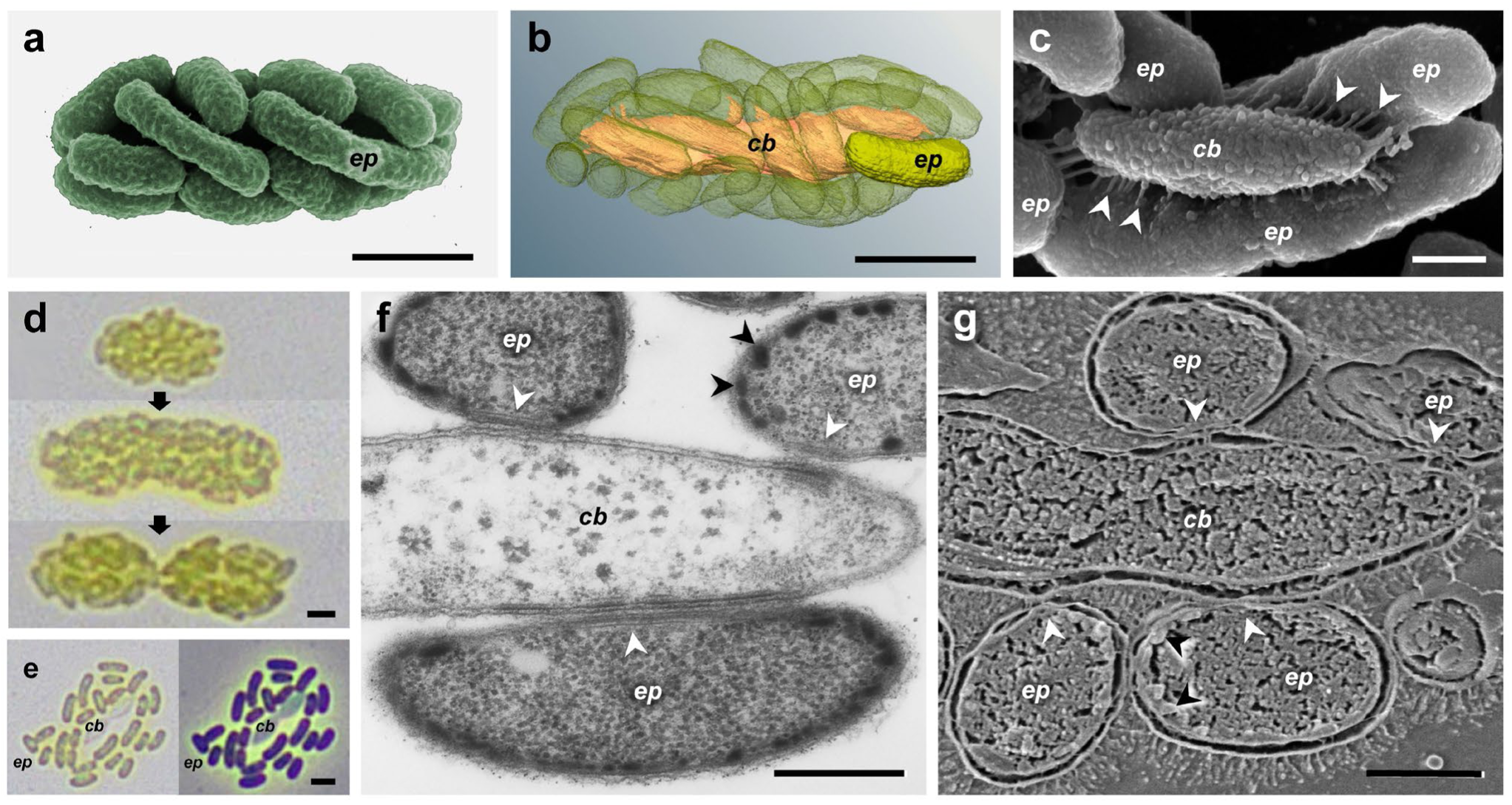
Morphology of ‘*Chlorochromatium aggregatum*’. **(a)** Colored scanning electron micrograph (SEM) of intact consortium depicting its shell of regularly ordered epibionts (*ep*). **(b)** 3D-reconstruction of a phototrophic consortium based on Focussed Ion Beam (FIB)/SEM tomography. Central rod colored yellow, epibionts transparent green. **(c)** SEM of partially disintegrated consortium, exposing periplasmic tubules (white arrowheads) which connect the central bacterium with the epibionts. **(d)** Bright field light-micrographs of subsequent stages of division, by which the two intact daughter consortia are formed. **(e)** Bright field (*left*), and phase contrast light-micrograph (*right*) of a single consortium disaggregated by pressure of the cover slip, thereby exposing a dividing central bacterium (*cb*) and 24 epibiont cells (*ep*). **(f)** TEM micrograph of an ultra-thin longitudinal section of a consortium showing adhesion sites between the central bacterium and epibionts, characterized by additional membranous structures (white arrowheads) and lack of chlorosomes (black arrowheads). **(g)** Cryo-SEM (hydrated high pressure cryofixation and cryofractured) of a consortium revealing tight, persisting contacts between the cell envelopes of epibionts and of the central bacterium. Bars, 2 µm in panels a,b,d,e, and 0.5 µm in panels c,f,g.

Here we report on the expression, interspecies transfer and subcellular localization of several symbiosis proteins and elucidate their possible functional role in maintaining the mutualistic interaction within ‘*C. aggregatum*’.

## Results and Discussion

### Unique symbiosis genes of the green sulfur bacterial epibiont

The epibiont genome (JBRGNI000000000; Fig. E1a) encodes 2,096 ORFs, of which 356 are missing in all known, non-symbiotic green sulfur bacteria (green arrow and dot, Fig. E1b; Table S1C). This number of species-specific genes was comparable to other *Chlorobiaceae* lineages (single, isolated black dots, Fig. E1b), indicating that symbiotic capacity does not depend on a particularly high gene gain. While 248 of the unique epibiont genes were hypothetical ORFs and 16 were transposons, the remainder were associated with interactions, photosystem arrangement, DNA recombination, motility/pili, capsule formation, secretion and signaling (Table S2). Three open reading frames (ORF) Cag_663, Cag_665, and Cag_2037 were of particular interest due to high homology with bacterial virulence factors (Table S3), and previous partial detection of their transcripts in consortia [**7,20**].

Cag_663 and Cag_665 represent two of the largest ORFs known to occur in prokaryotes at a length of 110,418 and 61,941 bp, respectively. If the respective 36,806 and 20,647 amino acids (aa) were fully transcribed and translated (Fig. E2a), the encoded proteins would only be rivaled in length by the exons of mammalian titin genes. Based on our improved sequencing, annotation, and domain characterization, both giant proteins were found to contain multiple novel regions, only showing homology to distant, unrelated, and typically uncultured bacteria (Fig. E2A, Table S3). They encompass N-terminal 24 aa LEPRxLL CS domains also found in other giant proteins [**21**], as well as C-terminal domains with highest similarity to a signaling magnetotaxis protein MtxA suggested to play a role in the aerotaxis of multicellular magnetotactic bacteria (MMB) [**22**]. Centrally located secreted-virulence-factor domains for hemolysin in Cag_663, and a crystalline βγ-Greek motif in Cag_665, were further detected by InterPro. Cag_663 also contained six HK97 folds spread at regular intervals over 4,000 aa near the N-terminus (Fig. E2a, Table S3); structures which typically occur as assembled coat subunits of phage capsids [**23**]. Cag_665 instead has multiple LktA-like tandem repeats known from secreted anaerobic leukotoxin proteins near its N-terminus (Fig. E2a; [**24**]). Yet, the most peculiar feature of both giant genes Cag_663 and Cag_665 were the numerous (50 and 49, respectively) filamentous hemagglutinin motifs (averaging 27 or 25 aa long, respectively), concentrated across the C-terminal half of either gene (Fig. E2a; grey triangles). These multiple β-sheets have highest, yet limited homology to VgrG secretion spike proteins of *Bacteroides fragilis* Type 6 secretion systems (Table S3; [**25**]), the β-helical shaft of filamentous hemagglutinin FhaB in *Bordetella* Type 5b toxin delivery systems [**26**], and to the hemagglutinin repeats of CdiA in *E. coli* Type 5b contact-dependent growth inhibition system targeting other *E. coli* strains [**27,28**]. Furthermore, Cag_663 and Cag_665 were separated by a TolC-like outer membrane protein (Cag_664), and preceded by Cag_662 annotated as *sapC* (Table S2; Fig. E2b), both found in many bacteria as Type 1 secretion system (T1SS) operon components [**29,30**].

The 4,581 bp-long Cag_2037 gene has been suggested to encode a C-terminal repeat-in-toxin (RTX) motif [**20**], as supported by multiple Ca^2+^ binding sites in the predicted model (Fig. 2). Upstream of the RTX motif, a mannuronan C5 epimerase/ lyase or extracellular alginate C5 epimerase domain was predicted by InterPro and HHpred, respectively (Fig. S2a, Table S3). Nine (bacterial)Immunoglobulin-like (BIg) folds extended and separated both termini, with an N-terminal Type II cohesin module identified. Known from anaerobic bacterial cellulosomes, enzymes with cohesin domains typically bind outer membrane scaffoldin dockerin modules with high affinity to degrade exogenous polysaccharide [**31**]. Searching the epibiont genome indeed revealed a corresponding dockerin domain in the neighboring Cag_2038, which also contained 18 central BIg folds (Fig. E3a). The C-terminal position of the Cag_2038 dockerin and the N-terminal localization of a cohesin domain in Cag_2037 align with the patterns observed from cellulosome proteins of anaerobic *Clostridia* and *Bacteroidia* [**32,33**], but were unknown for *Chlorobiota*. The size of the Cag_2037 gene product is well in the range of substrates of T1SS [**34**]. At the Cag_2037 C-terminus, a 40 aa transmembrane domain constituting a secretion signal characteristic of T1SS substrates in Gram-negative pathogens was also detected [**34,35**].

**Fig. 2.**
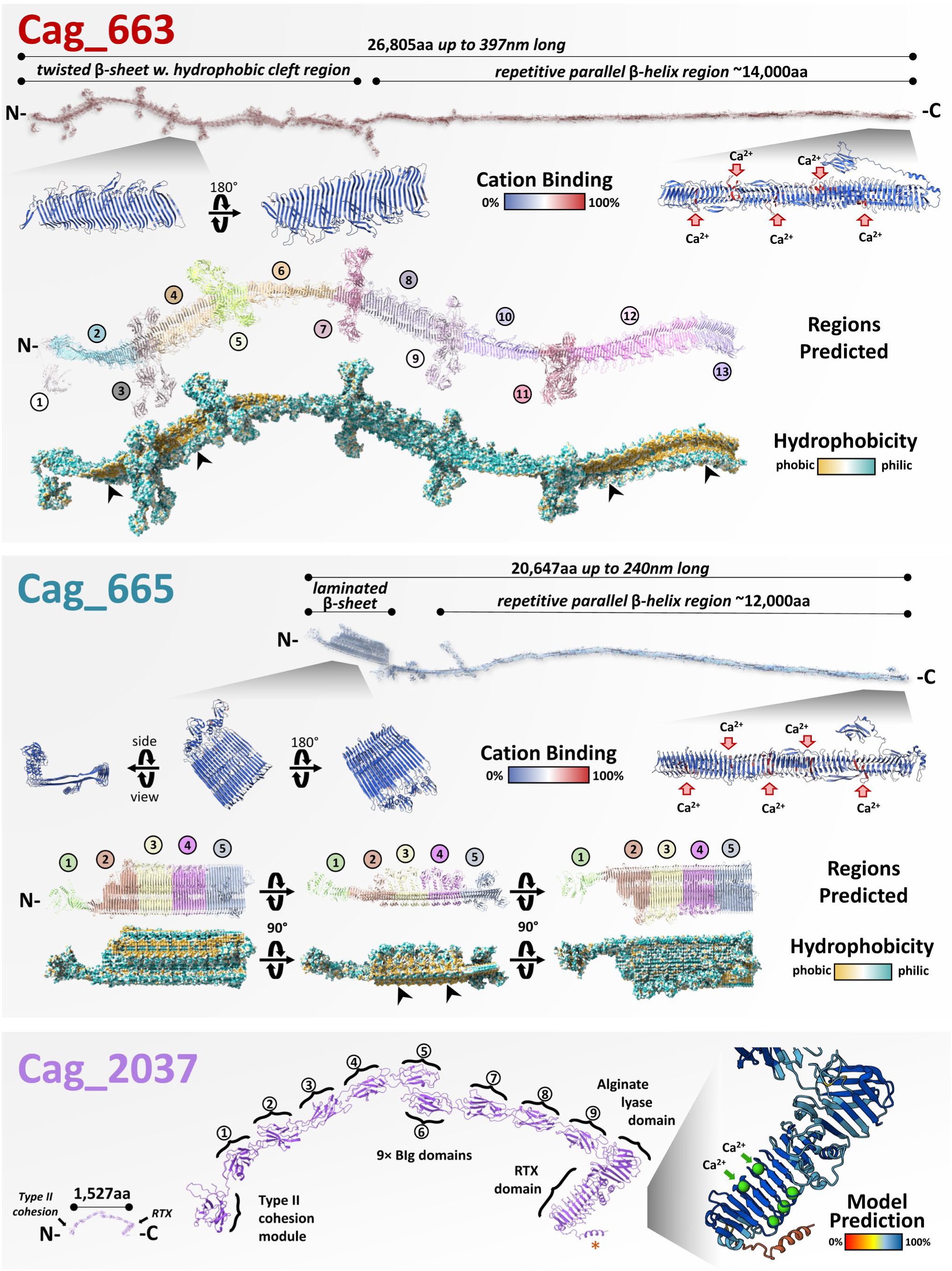
Structural modelling of *Chl. chlorochromatii* symbiosis proteins encoded by Cag_663, Cag_665 and Cag_2037. The giant 26,805 aa Cag_663 required an assembly of 32 overlapping ∼1,500 aa predicted domains (*AlphaFold3*; Fig S1a) to generate a structure measuring nearly 400 nm long. It comprised a C- terminal elongated repetitive parallel β-helix of 14,000 aa that had multiple predicted calcium cation bindings sites (*Pesto* and *InterPro*), and a N- terminal twisted β-sheet with hydrophobic cleft region (arrows) that had regular intervals of outstretched HK97 folds, shown in expanded view (detailed in Fig. E5c). The second large Cag_665 of 20,647 aa required 17 ∼1,500 aa predicted domains (*AlphaFold3*; Fig S1b) to generate a similar extended structure measuring nearly 240nm. Its 12,000 aa C- terminal was identical to Cag_663 as an elongated repetitive parallel β-helix with calcium cation bindings sites (*PeSTo* and *InterPro*). However, a unique N- laminated β-sheet with hydrophobic domain replaced the N-terminus of this protein (detailed in Fig. E5d). The 1,527 aa Cag_2037 schematic model reveals a N- terminus type II cohesion module followed by 9 bacterial immunoglobulin-like (BIg) domains, outstretches an alginate lyase-linked RTX domain that has a small C- terminus transmembrane helix (orange asterisks). As expected, the RTX domain prediction is supported by additional bound Calcium cations (details in Fig. S2a).

While secreted hemolysin-related proteins of the RTX toxin family were long known for pathogens and more recently also detected in bacterial predators [**36**], they were so far not reported to be involved in mutualistic symbioses. Matching the secretion signals of Cag_2037, *Chl. chlorochromatii* encodes an unusually large number of T1SS components for protein export (16 *hylB*, 6 *hylD*, 9 *tolC* homologs; Table S4), compared to all other *Chlorobiota* (Fig. E4a). 23 of the 31 homologs occur in six distinct operons encoding all three necessary components of a T1SS (Fig. E2c, Table S4). Again, this is higher than in other bacteria containing duplicate T1SSs [**37**], where up to five have been found in pathogenic bacteria, secreting different substrates [**38**]. As Cag_2037 shares highest homology to genes of non-related bacterial pathogens (Table S3), it was likely acquired in parallel to some of the T1SS through horizontal transfer (HGT). Alternatively, the high number of T1SS components might have resulted from gene duplication as seen in betaproteobacterial pathogens [**37**]. Our detailed phylogenetic analysis revealed that nine genes, of which four are in an operon (ORFs Cag_939-942 of operon Cag_936-942), likely were recently obtained from *Betaproteobacteria* or *Desulfuromonadales* (Fig. E3b, Table S4). Notably, Cag_662 *sapC*, Cag_664 *tolC* and Cag_1507 *hlyD* were also acquired by HGT from such sulfur-reducing bacteria, where Cag_1507 is homologous to a gene in MMB (Table S4). Beyond T1SS, a complete *lolCDE* T9SS [**39,40**] could be identified as vertically inherited in addition to *sec* and *sec*-independent systems, with only one Cag_1676 *secA* suggested to be recently acquired (Table S4). Finally, the repetitive filamentous hemagglutinin motifs of Cag_663 and Cag_665 are likely also relevant for protein excretion, as similar hydrophobic interactions between β-strands have been predicted to form large needle- or shaft-like modules capable of autotransport in the structurally related VgrG, CdiA and FhaB proteins [**25,26**].

### Predicted structures of symbiosis proteins

To further assess the annotated domains of these putatively excreted symbiosis factors, schematic reconstructions of each theoretical protein enabled hydrophobicity, electrostaticity, cation binding, polymerization, and size of the resulting structures to be assessed (Figs. 2, E3b, E5a,b). Because of the extremely large sizes of Cag_663 and Cag_665, modeling had to be performed separately for 1500 aa-fragments in AlphaFold3. By choosing longer sections than in previous approaches [**41**] and through modeling of overlapping fragments, predictive power was significantly improved so that 3D-structures of the entire proteins could be obtained via multiple alignments in PyMOL (see Methods; Fig. S1).

Complete models revealed rigid (low B-factor), elongated, filamentous structures for Cag_663 and Cag_665, extending 240-400 nm for Cag_665 and Cag_663 (Fig. 2a,b). In both symbiosis proteins, the repetitive β-helix structures spanned 12,000 to 14,000 aa from the C-terminus with only one or two other recognizable domain types, and resembled the overall structure of the Vgr spike protein that our domain searches had identified as closest homologs (see above). Proteins with similar repetitive β-helix domains occur in diverse bacteria including not-yet-cultured Gram-negative and Gram-positive bacteria, as well as MMB (Table S5A,B; Fig. E5a,b), which suggests that these filamentous domains are distributed more widely among prokaryotes, particularly among rare multicellular bacteria. The folded β-helix regions were shown to feature hydrophilic surfaces containing multiple cation binding sites (every 249±57 aa; Fig. 2a,b) predicted with high probability (97.8% likelihood; Table S5C). Since parallel β-helices, Ig-folds, BIg-folds and RTX-toxin β-roll domains have been shown to specifically bind Ca^2+^ [**42**], we inferred that the cation binding sites identified in the symbiosis proteins are also occupied by this ion (Fig. 2). The binding of 59 or 40 Ca^2+^ ions along the filamentous C-terminal region of Cag_663 and Cag_665, respectively, would contribute to their rigid elongated structure [**43**], whereas the 21 Ca^2+^ ions in Cag_2037 are predicted to stabilize the elongated structure formed by 9 BIg-/Ig-folds as well as the β-roll of the RTX domain (Fig. 2; [**42**]). For the RTX motif, inclusion of five Ca^2+^ also improved the modelling (Fig.2; Fig. S2). Ca^2+^-dependent folding is expected to occur only in the extracellular freshwater environment where Ca^2+^ concentrations reach the required range (0.1-2.5 mM) [**43**]. The predicted binding of numerous Ca^2+^ ions is in line with an extracellular localization of the repetitive β-helix structures of Cag_663 and Cag_665, and of the Cag_2037 protein. In fact, independent, experimental evidence for the role of Ca^2+^ in the cell-cell-interaction within ‘*C. aggregatum*’ exists: The highly ordered consortia of ‘*C. aggregatum*’ can only be maintained with 1.7 mM Ca^2+^, whereas consortia rapidly disintegrate when laboratory cultures are treated with EGTA, a chelating agent specific for Ca^2+^ [**20**].

In contrast to their similar filamentous β-helix structures, modelling of the N-termini of Cag_663 and Cag_665 giant proteins yielded different and distinct structures which did not match any structurally characterized proteins to date. The first 13,000 aa encoded by Cag_663 form an extended β-sheet that develops a continuous hydrophobic cleft running along one surface (Figs. 2, E5c). Five capsid-like/HK97-fold domains (Table S3; [**23**]) are found protruding at regular intervals without disrupting the sheet. Considering possible polymerization, two Cag_663 proteins potentially dimerize by sheltering their hydrophobic clefts together, leaving co-located capsid-like/HK97-fold motifs inversely exposed to the outsides (Fig. S3). In comparison, the β-helix repeats of Cag_665 at its N-terminal 5100 aa were predicted to form a laminated plate-like structure, consisting of two highly regular layered β-sheets (Figs. 2, E5d). Again, this region showed no homology to any characterized protein or structure, but the presence of hydrophobic surfaces may suggest interactions with other proteins or membranes. However, similar to C-termini, fragments of both N-termini did show highest homology to predicted proteins from MMB draft genomes (Fig. E5).

The model of Cag_2037 revealed domains similar to those predicted from aa homology searches (Fig. 2). Preceding the β-roll of the Cag_2037 RTX-domain, a 200 aa-long region of Cag_2037 not only showed sequence similarity to a known alginate epimerase of *Azotobacter vinelandii* (Table S3) but also matched its structure (Table S5A). The gene product of upstream Cag_2038 was further shown to generate a filamentous structure formed by the repetitive BIg domains, where its C-terminal dockerin domain was exposed through a protein bend, and found to bind the N- terminal cohesin domain of Cag_2037 as suggested in literature (Fig. E3b; [**32,33**]).

### Expression of symbiosis genes and co-expression of other genes

While partial transcripts of the three epibiont symbiosis genes had been detected previously [**7**], key aspects remained unclear: **(i)** whether the unique, giant genes were actually transcribed over their full length, **(ii)** if gene expression was affected by the symbiotic state or growth phase of the epibiont, and **(iii)** which other epibiont or central bacterium genes, particularly those related to protein secretion, are co-expressed and vital for heterologous multicellularity.

For global transcriptomic profiling, cultivation conditions of pure epibiont and consortia were standardized and growth was tracked employing different measures for biomass (Materials and Methods; Fig. E6). This ensured that cells from the axenic epibiont and consortia cultures were sampled in similar growth states (exponentially growing or in stationary phase; Fig. 3a), enabling a direct comparison of gene expression patterns (Fig. S4). Aligning transcripts along the *Chl. chlorochromatii* genome (Fig. E7a) showed all three symbiosis genes to be transcribed over their entire length. Thus, Cag_663 and Cag_665 could potentially give rise to giant proteins, while both were separated by the small ORF Cag_664 for which specific transcripts were detected. In contrast, the Cag_2037 transcript was part of a bicistronic mRNA that extended through the upstream ORF Cag_2038. The relative abundances of all symbiosis gene transcripts varied with the growth phase and symbiotic state of the epibiont. The two filamentous hemagglutinin-like Cag_663 and Cag_665 were among a small group of 19 genes which maintained highest relative cellular transcript counts in symbiotic *Chlorobium*, particularly during the H_2_S-limited stationary phase of growth (Fig. 3b). Both genes were further up-regulated in exponentially growing consortia as compared to axenic *Chlorobium* epibiont cells. While the RTX-alginate lyase Cag_2037 (and associated Cag_2038) was also highly transcribed in stationary phases vs. exponentially grown cells, it did not significantly change in transcription depending on symbiotic state, similar to 111 other epibiont genes (Fig. 3c). These patterns suggest that the three symbiosis proteins are most relevant during sulfide limiting conditions, with some particularly expressed during symbiotic interactions of consortia.

**Fig. 3.**
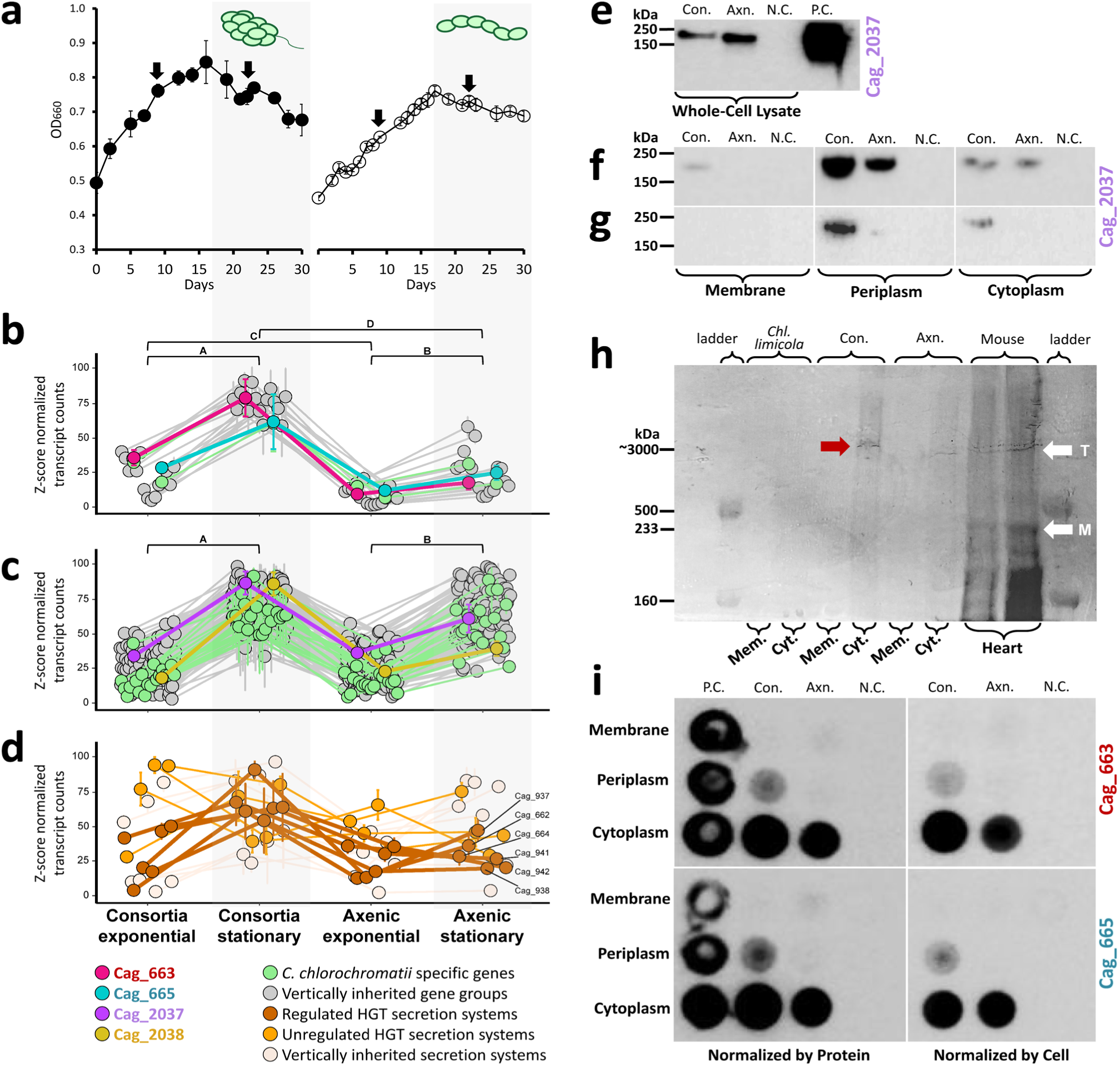
Expression of giant symbiosis genes. **(a)** The axenic epibiont *Chl. chlorochromatii* (white dots) and intact ‘*C. aggregatum*’ consortia (black dots) were grown in triplicate cultures, supplied with sulfide during exponential growth, and then starved for sulfide to induce the stationary phase. Cells were harvested for transcriptomic analysis in exponential or stationary phases of growth as indicated by black arrows. **(b)** Expression pattern of 19 *Chl. chlorochromatii* genes, including Cag_663 and Cag_665, with highest expression in stationary consortia (significant differences A,B,C,D; Table S7). **(c)** 112 genes, including Cag_2037 and Cag_2038, which are significantly up regulated in exponential phases regardless of symbiotic state. **(d)** Expression patterns of vertically inherited or horizontally transferred protein secretion systems. **(e)** Western blots of whole-cell lysates showing Cag_2037 protein to be present in consortia and axenic *Chl. chlorochromatii*. **(f,g)** Western blots of membrane, periplasm and cytoplasm fractions each **(f)** normalized by protein or **(g)** normalized by number of *Chlorobium* cells. **(h)** Extracted proteins run on 2% agarose stabilized polyacrylamide gel, reveal large peptide running as a similar size to titin protein (white arrow, T) and myosin (white arrow, M) from mouse cardiac tissue confirmed via MS (Table S11A). **(i)** Dot blots of Cag_663 or Cag_665 proteins in membrane, periplasmic and cytoplasmic fractions of consortia and axenic *Chl. chlorochromatii*, normalized to protein content (left) and numbers of epibiotic cells (right). Con., consortia; Axn., axenic *Chl. chlorochromatii;* N.C., *E. coli* XL-1 Blue cells serving as negative control; P.C., positive controls consisting either of purified recombinant protein of Cag_2037 or purified recombinant protein fragments of Cag_663 or Cag_665.

In order to identify additional genes with potential specific relevance to the symbiotic interaction, a network analysis of epibiont transcriptomes across the different growth conditions was first conducted, showing that the expression followed four major patterns (corresponding to four network clusters in Fig. E7b, Table S6). Of the overall 2,152 epibiont ORFs, cluster 1 comprised the largest number of 579 genes encompassing Cag_663, Cag_665 and Cag_2037 but also the neighboring SapC (Cag_662), and TolC (Cag_664) homologs, as well as the dockerin-domain containing Cag_2038. Notably, several genes of the horizontally acquired T1SS (Cag_937,_938,_941,_942) followed a similar transcription pattern (Fig. E7c, Table S6B). Cluster 2 mostly comprised genes up-regulated in exponential phases of growth, and encoded metabolic enzymes, photosynthesis, tRNAs, and genes of additional T1SS’s (Cag_781,_782,_940, _1065,_1066,_1507; Table S6C), while cluster 3 encompassed genes for ribosomal proteins, numerous hypotheticals, a number of specific metabolic enzymes, and of a fourth T1SS operon (Cag_807,_808; Table S6D), whereas cluster 4 had the three rRNAs with various ribosomal proteins, tRNAs, several photosynthesis genes, nitrogenase genes, and a fifth and sixth T1SS (Cag_42,_43,_363,_364_365; Table S6E). As several secretion systems fell into different clusters, we analyzed their transcriptional patterns in more detailed comparisons (Figs. 3d; Tables S7A-N). Supporting results of network analysis above, symbiotic proteins (Fig. S4c; Fig. S5; Tables S7A-N), and most ORFs encoding the horizontally transferred T1SS ORF (Cag_937,_938,_941_942) largely followed the transcriptional patterns observed for the three symbiosis genes (Fig. 3d). Of the 2,152 *Chl. chlorochromatii* ORFs (including 274 unique epibiont and 154 *Chlorobiaceae*-specific genes), 693 ORFs were found to be constitutively expressed (Fig. S4a,b; Table S7N).

The considerable number of coexpressed genes are a first indication that additional common or unique genes are of functional relevance for the mutualistic interaction. The coordinated expression of Cag_2037 and Cag_2038 (Fig. 3c) corresponds with the interaction of both gene products through complementary cohesin and dockerin modules inferred by AlphaFold 3 modeling (Fig. E3a,b). Based on their coexpression, Cag_937-942 together constitute the most probable T1SS candidate for exporting the Cag_2037 RTX-like protein and were likely acquired by HGT from a *Betaproteobacterium* or *Desulfuromonadales* that are known to co-occur with phototrophic consortia in the chemocline of freshwater lakes. This is in contrast to the cooption of a preexisting T1SS after the acquisition of a β-hemolytic RTX toxin that for instance has occurred in the evolution of pathogenic *Betaproteobacterium* genus *Kingella* [**37**]. However, interactions also typically involve additional cellular functions beyond virulence factors, e.g. in metabolism [**44**] which often evolve from preexisting genes [**45**]. Indeed, the considerable number of non-epibiont-specific ORFs showing similar transcriptional patterns as symbiosis genes suggests that, besides HGT events, additional preexisting genes were recruited during evolution of mutualism in the epibiont and hence constitute potential pre-adaptations.

Transcriptomics also yielded new insights into the response of ‘*Ca.* Symbiobacter mobilis’ under different growth conditions of phototrophic consortia. While its small genome shares 777 genes with all other *Comamonadaceae* species [**18**], we found 254 genes missing as a result of genome reduction (Fig. E8a; Table S8A-C). Despite the depleted gene content, 657 ORFs are still unique to ‘*Ca.* Symbiobacter mobilis’ (Table S8D), including genes related to flagellation, pili and quorum sensing that are relevant for symbiosis (Table S9A). Complete Type IV pili and flagellated motility are common among the *Comamonadaceae* (Table S9B), indicating pre-adaptation. A total of 301 genes of the central bacterium changed expression in response to the consortia growth state (Fig. E8; Table S10A). Clustering of regulatory groups (Fig. S11b; Table S11B) showed maximum levels of flagellum- and pili-related gene expression coinciding with the limitation by sulfide (Figs. E6, E8c), suggesting induction of adhesion and motility. Flagella and pili related genes were among the most highly expressed during sulfide starvation, underlining their conditional relevance to the consortia. Several unique chemotaxis genes were differentially expressed either in exponential or stationary phases, with chemoreceptor Smb_1671 switching with Smb_774 in the respective growth states, and therefore may be linked to sulfide limitation. Bacteriophytochrome Smb_130 is also replaced with a highly expressed Smb_2467 in stationary phase, a 700-800 nm red-light sensor previously shown to be utilized by ‘*Ca.* S. mobilis’ [**18**], implying that its scotophobic response [**10**] is also growth phase dependent.

To date, none of the symbiosis proteins had been detected by mass spectrometry analysis of the cytoplasmic and membrane proteomes [**46**], or by exploiting their Ca-binding properties via ^45^Ca^2+^ autoradiography [**20**]. Therefore, immunoblotting of different cell fractions was conducted to increase the sensitivity of detection and to gain initial information on the cellular localizations of the three proteins (Fig. 3e-i). Cag_2037 protein was observed as a single band in Western blots of both consortia and axenic epibiont whole-cell extracts (Fig. 3e). It constituted a large fraction of the periplasmic proteins and a lower amount in the cytoplasm of both consortia and axenic epibiont, whereas only traces of Cag_2037 protein were detected in the membranes of ‘*C. aggregatum*’ (Fig. 3f). On a per cell basis, however, Cag_2037 was predominant in the periplasm and less abundant in the cytoplasm of the consortium, while it was barely detectable in the periplasm of axenic epibiont cells (Fig. 3g). In agreement with their inferred large molecular size, the Cag_663 and Cag_665 proteins did not enter gel matrices of standard acrylamide gel concentrations, and barely entered agarose stabilized 2% acrylamide gels like similarly gigantic titin (Fig. 3h). Here, existing bands of titin and myosin were confirmed via trypsin-digests, while the large peptides in consortia remained enigmatic (Table 11A). *In silico* digest revealed both giant proteins to have limited trypsin target sites, suggesting them to be resistant to excreted proteases, which might represent an evolutionary adaptation as an external protein. Multiple runs of whole-cell proteomics incorporating multi-enzyme digests and higher SDS content revealed both to have multiple peptide hits (Table 11B; Fig. S6), where Cag_663 had better identification rates than Cag_665 (323 vs. 40 identified peptides, 14.72% vs. 2.56% coverage, respectively). While suggesting these proteins are difficult to degrade, the whole cell digest coverages mirrored *in silico* digests, confirming both to be present within the consortia biomass as produced proteins (Fig. S6). Yet, dot immunoblot analyses (Fig. 3i) revealed that both proteins were mostly present in the cytoplasmic fractions of consortia and axenic epibionts. Notably, a smaller, but clearly detectable, signal was observed for the periplasmic fractions of ‘*C. aggregatum*’ but not for axenic epibionts. Based on our results, all three symbiosis proteins are in fact synthesized in axenic as well as symbiotic epibionts. Proteins matched the predicted sizes indicating no significant processing of transcripts. Most importantly, the presence of all three symbiosis proteins in the periplasmic fraction provides evidence for protein export, which supports the results of our protein domain analysis and structure predictions. The different amounts detected in consortia *vs.* axenic epibiont periplasm further suggests that not only gene expression but also the protein distribution patterns across cellular compartments change between the free-living and symbiotic growth states.

### Cellular and subcellular localization of symbiosis proteins

Two types of super-resolution immunofluorescence (IMF) microscopy were used to directly localize the three symbiosis proteins in cells of intact consortia or axenic *Chl. chlorochromatii* epibionts (Fig. 4, Fig. E9). Specific antibodies labeled with Alexa 488 nm were employed in total internal reflection fluorescence (TIRF) microscopy, whereas Alexa 647 nm was used for the higher resolution direct stochastic optical reconstruction microscopy (dSTORM). The natural autofluorescence of bacteriochlorophyll (BChl) *c* in chlorosomes served to distinguish the phototrophic epibiont cells from the central bacterium (see Materials and Methods), while the cytoplasmic DNA of all cells was stained with 4’,6-diamidino-2-phenylindole (DAPI).

**Fig. 4.**
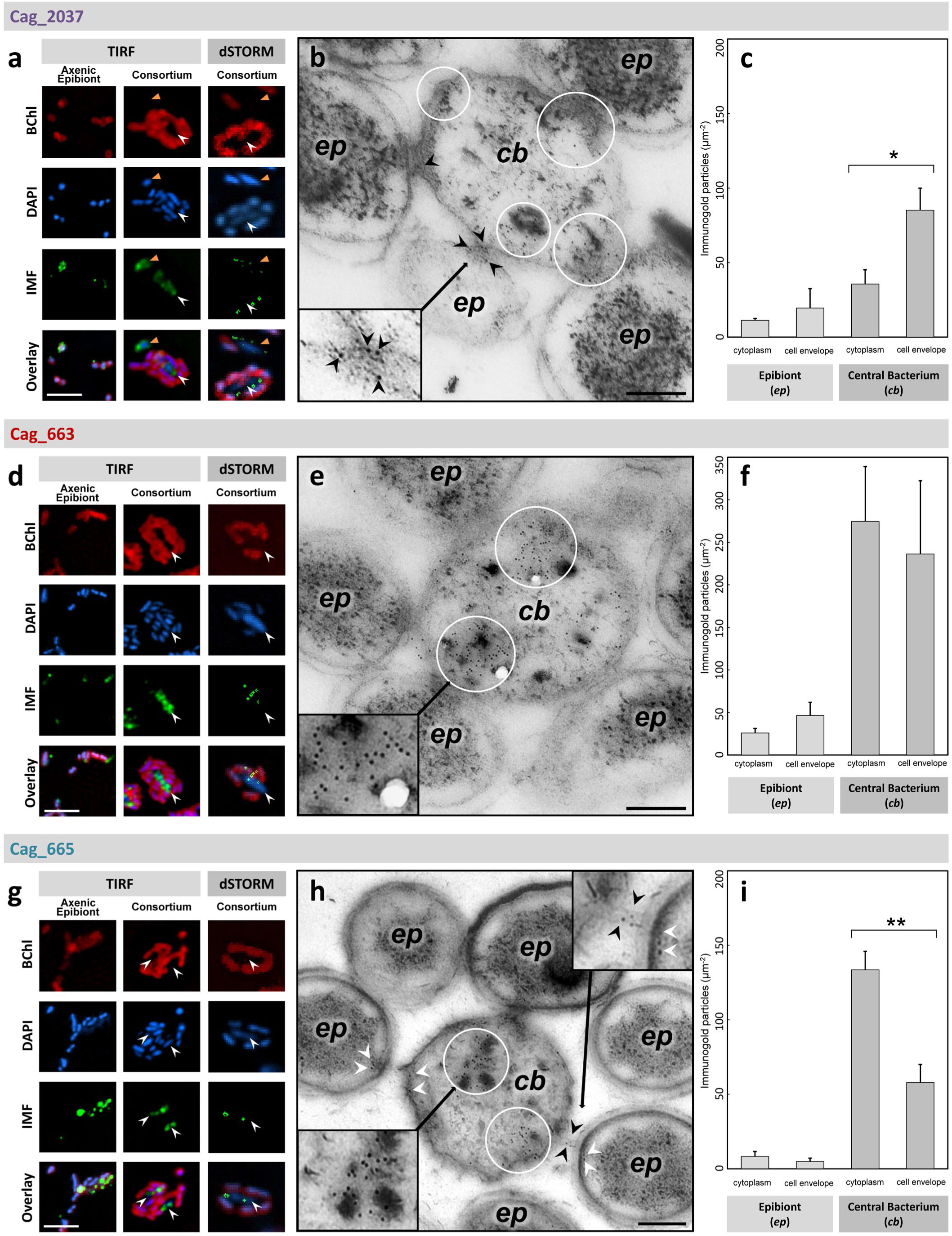
Cellular and subcellular localization of the three symbiosis proteins. TIRF and dSTORM microscopic analysis of intact phototrophic consortium ‘*C. aggregatum*’ and its epibiont *Chl. chlorochromatii* CaD in axenic culture employing antisera against the proteins **(a)** Cag_2037, **(d)** Cag_663 and **(g)** Cag_665 among bacterial cells. Fluorescence images for bacteriochlorophyll *c* (BChl), 4’,6-diamidino-2-phenylindole (DAPI), and Alexa Fluor 488nm immunofluorescence (IMF) of the individual targeted symbiotic proteins are depicted separately as well as overlay. White arrows indicate the position of the central bacterium ‘*Ca.* S. mobilis’ (*cb*) and orange arrows show a single cell separated from its epibionts (*ep*) during preparation of the wet mounts. Central panels depict transmission electron micrographs of cryosectioned ‘*C. aggregatum’* treated with immunogold-labeled antibodies targeting proteins **(b)** Cag_2037, **(e)** Cag_663, and **(h)** Cag_665. Accumulations of gold particles are noted by white circles. Arrows mark proteins detected in or in the vicinity of the periplasmic tubules at the contact sites between epibionts and central bacterium. Right panels give quantification of **(c)** Cag_2037, **(f)** Cag_663, or **(j)** Cag_665 protein-specific immunogold-particles per µm^2^ in different cell compartments of ‘*C. aggregatum*’ in across the TEM micrographs. Vertical bars show standard error. Significant differences were determined by pairwise t-tests as indicated (*, p<0.05; **, p<0.01). Fluorescence and TEM scale bars, 5 µm and 200 nm, respectively.

Despite being encoded by the epibiont genome, Cag_2037 proteins exclusively co-localized with ‘*Ca.* S. mobilis’ in consortia IMF preparations (Fig. 4a; white arrows), including central bacterium cells that had been separated from their epibionts during preparation of wet mounts (Fig. 4a; orange arrows). In strong contrast to the phototrophic consortia, a significant number of axenic *Chl. chlorochromatii* epibiont cells expressed and retained the protein Cag_2037 (Fig. 4a), averaging 44.1% (±7.2%, n=442) of the cells. At higher resolution, dSTORM detected Cag_2037 proteins at up to six well-separated positions per consortium (Fig. 4a). Most signals of TIRF and dSTORM were positioned at the cell periphery of ‘*Ca.* S. mobilis’, outside of the DNA-containing central regions, and among cells that were completely dislodged from their epibionts. Subsequent immunogold transmission electron microscopy (TEM) of cryosectioned consortia substantiated the subcellular localization of Cag_2037 which was predominantly found within the central rod ‘*Ca.* S. mobilis’ cellular envelope, particularly among contact sites with epibiotic cells (Fig. 4b). A quantitative analysis of immunogold particles per µm^2^ among cellular cross-sections revealed that 79.9 % of the signals detected were associated with the central bacterium as a whole and that the largest fraction (56.3 %) of the signals was found within the cell envelope of ‘*Ca.* S. mobilis’ (Fig. 4c).

IMF detected Cag_663 and Cag_665 gene products predominantly in consortia where they also co-localized with the central bacterium (Fig. 4d,g). In contrast to Cag_2037, a tenfold lower fraction of only 4.4% (n=618) and 3.8 % (n=726) of the axenic epibiont cells yielded IMF signals for either Cag_663 or Cag_665, respectively. The higher resolution dSTORM corroborated this co-localization of the two symbiosis proteins with the central rod. In some instances, series of signals reaching from epibiont cells into central bacteria could be discerned (as shown for Cag_663 in Fig. 4d). In line with these results, immunogold-labeled Cag_663 and Cag_665 antibodies bound to proteins in the ‘*Ca.* S. mobilis’ cytoplasm rather than to epibiont cells (Fig. 4e,f, and 4h,j, respectively). While Cag_663 proteins were found to be equally distributed between the cell envelope and cytoplasm of the central bacterium (Fig. 4f), Cag_665 was predominantly detected in its cytoplasm (Fig. 4j). Notably, immunogold particles were frequently observed at the location of periplasmic tubules that protrude from epibiont cells and connect them with the central bacterium (Fig. 4b, h; black arrows, inserts) [**15**], resembling the delivery of RTX-proteins by outer-membrane vesicles to the target cell in several pathogenic bacteria [**34,35**].

Together, our findings show that the synthesis and localization of all three symbiosis proteins largely follows the patterns of gene expression, whereby the Cag_2037 protein is detected equally among axenic or symbiotic states, whereas Cag_663 and Cag_665 proteins are found less frequently in axenic cultures and upregulated in consortia, and all three migrate to the central bacterium from the epibiont cell envelope in consortia. While the canonical production and secretion of RTX proteins (such as HlyA of *E. coli* or CyaA of *Bordetella bronchiseptica*) occurs by a one-step mechanism through their cognate T1SSs, few large RTX-type adhesins (LapA of *Pseudomonas fluorescens* and the ice-binding adhesin of *Marinomonas primoryensis*) remain stalled as a secretion intermediate at the cell surface tethered to TolC [**47**]. Here, we discovered a previously unknown pattern in the phototrophic ‘*C. aggregatum*’, where RTX-type and other symbiosis proteins are synthesized early, but remain in the epibiont periplasm if the partner bacterium is absent. Only during cell-contact, will all three symbiosis proteins actually be excreted and transferred to the cell envelope or cytoplasm of the heterotrophic partner. We conclude that the export, rather than expression, of symbiosis proteins is impaired in the axenic state of the epibiont and occurs only during contact with the central bacterial symbiont. For the Cag_2037 RTX-protein, the Cag_937-942 T1SS represents the most likely candidate system for entire transmembrane translocation since **(i)** its components largely followed the transcriptional patterns of the RTX-protein and **(ii)** were similarly horizontally transferred to the epibiont genome (as indicated by the presence of transposons and sequence similarities; Fig. E2b; [**20**]). The early expression of *hlyB*-homolog Cag_940 likely causes the unfolded Cag_2037 to be transported and reside in the periplasm, prior to an excretion trigger producing and utilizing the remainder of the T1SS for full external export. This again is in contrast to other bacteria, which rely on the accumulation of RTX protein to induce the assembly of complete T1SSs [**34**].

While Cag_663 and Cag_665 proteins lack known signals for transport systems, the similarly expressed neighboring genes Cag_662 and Cag_664 are homologs of *sapC* and *sapF*/ *tolC*, respectively. Known to transport virulence factors without conventional N-terminal signal sequences, the *sap*-system does so for proteins C- terminus first [**29**], which matches the location of filamentous hemagglutinin repeats homologous to T6SS spikes in both Cag_663 and Cag_665. Moreover, as Cag_1507 (*hylD*) follows Cag_663, Cag_665 by sharing highest similarity to MMB, it is the most probable associated membrane fusion protein (mfp) candidate. We therefore hypothesize that Cag_662, Cag_664, and Cag_1507 are involved in the transport of the two giant epibiont proteins.

### Functionality of a new type of RTX-domain alginate lyase

Of the three symbiosis proteins, specific functionalities could only be inferred for several domains of the Cag_2037 protein. RTX toxins are widespread among bacteria as essential components for microbial interaction [**48,49**], being involved in mammalian pathogenesis [**37**], nodulation of plant root cells [**35**], endosymbiosis of sulfur-oxidizers and pathogenesis of mussels, tube worms, or polychaetes [**48,50,51**], as well as for adherence to ice and diatoms [**52**]. Most of the >1000 RTX protein family representatives that occur in >250 bacterial species are characterized from antagonistic interactions with eukaryotes [**35,52**], whereas none have been demonstrated for interspecific bacteria-bacteria interactions, let alone from mutualistic systems. While many are pore-forming cytotoxins, bacteriocins, adhesins or S-layer proteins, several also contain zinc metalloprotease and lipase domains that aid in releasing cytoplasmic constituents [**34,53**]. The lack of central bacterium lysis or cytoplasm degradation suggests Cag_2037 is not a harmful toxin in consortia, whereas the putative alginate epimerase/lyase domain indicated it could transform or break polymers. Although Cag_2037 contains an additional cellulosome anchoring region, the combination of RTX and bifunctional mannuronan C5 epimerases/alginate lyase domains is most similar to AlgE1-E7 of *Azotobacter vinelandii*, which likewise utilize a T1SS to bind and act on the external cell envelope [**54,55**]. Consequently, a better understanding of this potential enzymatic function would provide key insights into how bacteria interact within phototrophic consortia.

Alginate is a predominant constituent of brown algal cell walls [**56**], but can be occasionally found in capsules of *Pseudomonadaceae* [**57,58**], including cysts of *Azotobacter* that render these bacteria resistant to desiccation and oxygen diffusion [**59**]. Indeed, thick capsules have been previously reported for natural populations of phototrophic consortia [**14**], but their subcellular structure and chemical composition had not been analyzed. A visualization of ‘*C. aggregatum*’ by TEM ruthenium red-treated thin sections revealed capsular material surrounds the central bacterium in the intercellular space (Fig. 5a,b; black arrows), but is absent at the cell-cell contact sites with epibionts (Fig. 5a). The alginate monomers mannuronic acid and guluronic acid (Fig. 5c) were confirmed by mass spectrometry from consortia culture biomass, while being absent in pure axenic cultures of the epibiont (Fig. 5d). Upon addition of exogenous alginate lyase, a macroscopically visible disappearance of consortia biofilms on the walls of culture flasks was observed (Fig. 5e) and a quantitative disintegration of the phototrophic consortia (Fig. 5f). Evidently, alginate represents a major constituent of the capsule that enables continued association of partner cells in intact ‘*C. aggregatum*’.

**Fig. 5.**
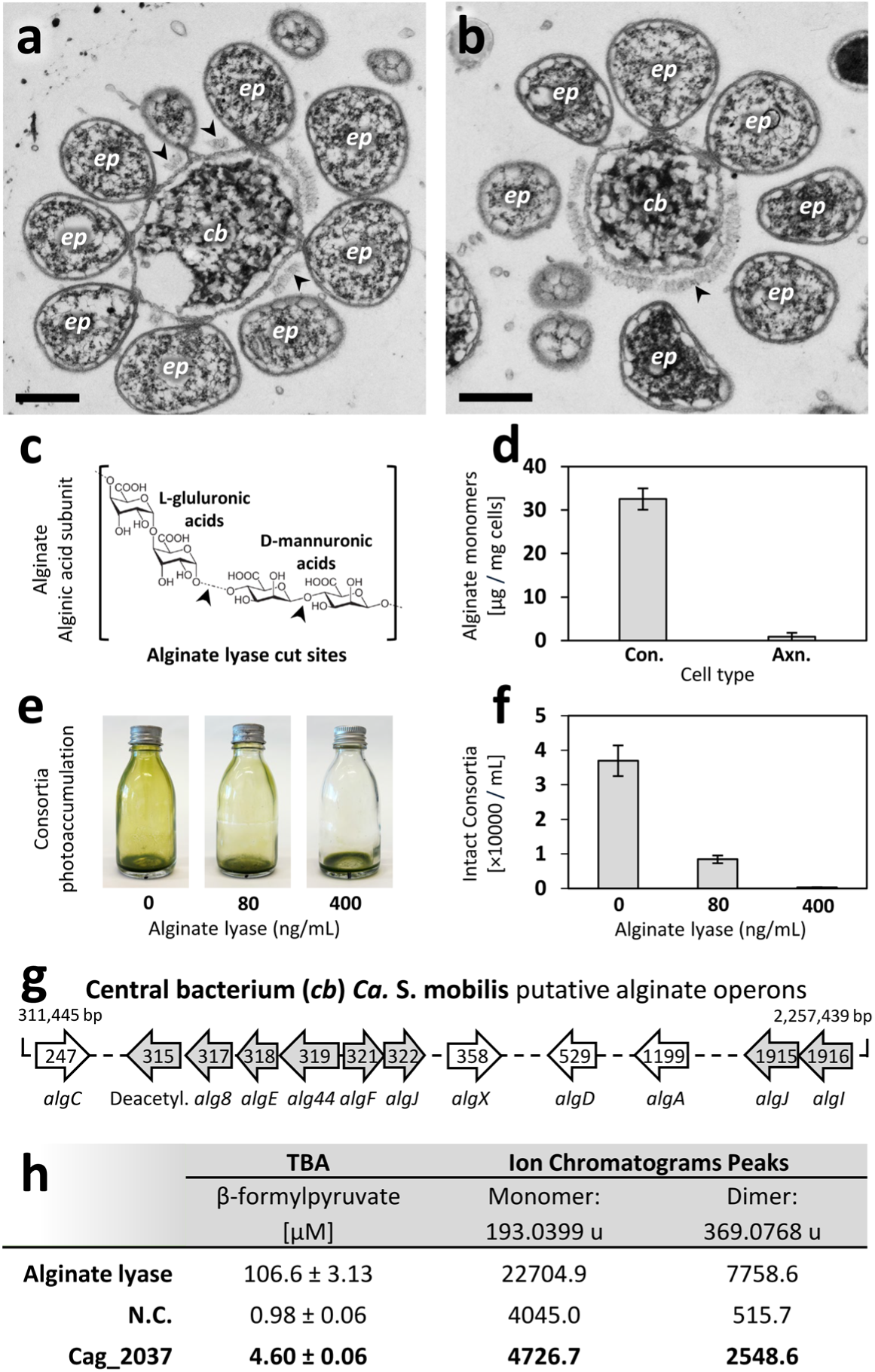
Alginate lyase degrades the ‘*Ca.* S. mobilis’ capsule. **(a-b)** TEM of consortia thin section fixed with LRR showing the presence of capsule-like material surrounding the central bacterium (*cb*) (arrows), absent at contact sites with epibionts (*ep*). Scale bar, 0.5 µm. **(c)** Alginate structure has alginic acid subunits of L-guluronic and D-mannuronic acid monomers, with putative alginate lyase cut sites (arrows). **(d)** Alginate monomers (mannuronic and guluronic acid) detected via mass-spectrometry (MS) in consortia cultures (Con.) and axenic epibiont cultures (Axn.) treated with exogenous alginate lyase. Minimal concentrations of exogenous alginate lyase induce **(e)** macroscopically visible dissolution of ‘*C. aggregatum*’ biofilms and reduction of phototactic ability, resulting from **(f)** microscopically observed disintegration of intact consortia. **(g)** Presence and localization of genes for alginate formation in the ‘*Ca.* S. mobilis’ genome. Locus tags provided on gene arrows without “Smb_” prefix. **(h)** Comparison of enzyme activity and degradation products of heterologously expressed Cag_2037 with commercially available alginate lyase through thiobarbituric acid (TBA) assay and by MS detection of alginate monomers or dimers.

A complete set of corresponding genes for the alginate biosynthetic pathway was detected in the genome of the central bacterium ‘*Ca.* S. mobilis’ (Fig. 5g), and can be found in other *Comamonadaceae*, suggesting vertical inheritance (Table S9B). Unlike gammaproteobacterial *Pseudomonas* and *Azotobacter* alginate synthesis genes arranged in two primary operons [**60,61**], the betaproteobacterial genes are split into several operons (Fig. S2h). Only *alg8, alg44, algEFJ* and a putative polysaccharide deacetylation gene formed the primary cluster in the central bacterium genome while the putative *algD*, *algA*, *algX*, as well as two copies of *algIJ* were found at distant loci. Most of the 14 genes directly involved in alginate biosynthesis of *‘Ca.* S. mobilis’ (Table S9A) were transcribed constitutively (with the exception of Smb_318,_321, _1481; Table S10A), suggesting a continuous need of capsular-production enzymes for consortium integrity.

To confirm the functionality of the RTX-type Cag_2037 alginate lyase, enzymatic activity of refolded and concentrated protein was assessed after heterologous expression in *E. coli* XL-1 Blue, employing the thiobarbituric acid (TBA) assay and MS assessment of degradation products (Fig. 5h). Compared to a standard endo-type alginate lyase purified from *Flavobacterium,* heterologously expressed Cag_2037 degraded alginate primarily into dimeric saccharides, but nevertheless acted as a weak, but significant, alginate lyase. In many bacteria, extracellular polysaccharides act as a defense against bacterial predators by inhibiting diffusion of metabolites or other compounds [**59,62**]. In the phototrophic consortium ‘*C. aggregatum*’, local degradation of the alginate capsule likely is a precondition for the direct cell-cell-contact between the mutualistic bacterial partners to enable nutrient exchange [**19**] and the observed transfer of Cag_663 and Cag_665 proteins into the central bacterium.

### A model of the novel bacterial mutualistic interaction and its general implications

Interspecies interactions occur frequently among complex bacterial communities, whereby negative associations like competition, amensalism or parasitism are thought to dominate [**63**]. Stronger negative interactions with a narrow target spectrum involve specialized cellular mechanisms which are considered a key selective force in the evolution of bacterial multicellularity, the origin of eukaryotes, and of human pathogens [**62**]. For instance, diffusible bacteriocins or secretion systems that deliver toxins upon cell contact are employed in interference competition [**64**] and a multitude of excreted proteases and specific adhesins are used by the > 100 known predatory bacteria from 12 different phyla [**36,65**].

The genomes of various bacterial predators often also contain large putative genes. In ‘*Ca.* Vampirococcus lugosií proteins up to 4,163 aa long were predicted to form fibrous cell surface structures that enable host-adhesion [**66**] and a large (5,898 aa) ‘grappling hook’ is utilized by *Aureispira* for binding its bacterial prey [**67**]. Bioinformatic analyses revealed the presence of even bigger (‘giant’) genes exceeding 20 kbp (>6,500 aa proteins) in genomes from 12 other bacterial phyla and 7 archaeal phyla [**41,68,69**]. Accordingly, giant genes/proteins were hypothesized to represent a general mechanism for bacterial predation. To date, the actual molecular size, localization, and functionalities of these supposedly giant proteins has remained enigmatic.

We here propose the first molecular model for the mutualistic interaction of ‘*C. aggregatum*’ (Fig. 6). The Cag_2037 protein enters the epibiont periplasm through constitutively expressed ABC transporter Cag_940. Under sulfide limitation, Cag_2037 and genes encoding the periplasmic and outer membrane components of T1SS Cag_939-942 are upregulated, resulting in extracellular excretion of Cag_2037 protein with its unfolded C- terminus first. In the extracellular milieu calcium triggers correct folding of its BIg domains [**70**], and the RTX motif into compact and stable structures [**71**]. Cag_2037 likely remains bound to the *Chl. chlorochromatii* cell surface through its N-terminal cohesin moiety adhering with associated dockerin domain of the Cag_2038 gene product. The numerous BIg domains may extend the Cag_2037 alginate lyase from the epibiont to reach and degrade the alginate capsule synthesized by the central bacterium. As RTX-domains are known for cell surface adherence [**52,72**], the RTX-domain of Cag_2037 may bind to the central bacterium. Bridging by Cag_2037 and Cag_2038 would generate the close and stable cell-cell contact regions observed between both partners [**15**].

**Fig. 6.**
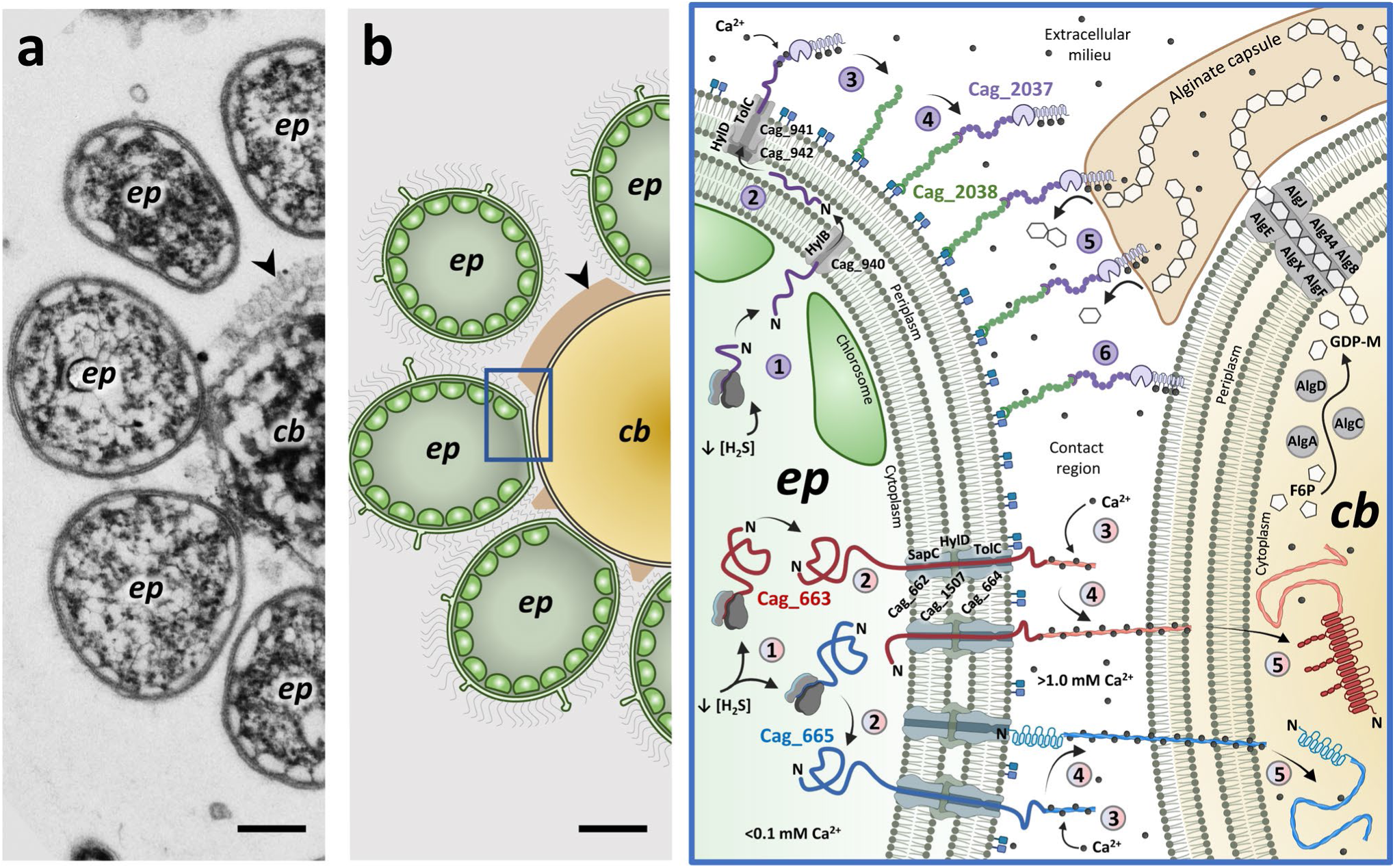
Putative model of symbiosis protein exchange and function. **(a)** Axial cross-section of consortia with LRR fixation depicting region with some epibiotic (*ep*) *Chl. chlorochromatii* contact regions, while others are near the central bacterium (*cb*) ‘Ca. S. mobilis’ but not at their respective contact site. Central bacterium alginate capsule develops in all regions without contact sites (arrow). Scale bar, 0.25 µm. **(b)** Schematic depiction of axial cross-section with contact region highlighted and enlarged. Central bacterium generates alginate capsule by converting fructose-6-phosphate (F6P) to guanosine diphosphate mannuronate (GDP-M) via AlgACD (Smb_1199, _247, _529, respectively), followed by extension-based export with Alg8-E-44FJ (Smb_317-322) and AlgX (Smb_358). In purple, (1) constitutively expressed HylB (Cag_940) excretes Cag_2037 to the periplasm, while sulfide limitation (<[H_2_S]) upregulates Cag_2037 and (2) the remaining T1SS: TolC-HlyBD (Cag_941-942) to transport it across the cell envelope. (3) In the extracellular milieu it will encounter and bind Ca^2+^ permitting correct folding and activation of RTX, alginate lyase, and BIg fold extender domains. (4) Localizing at the subcellular *Chl. chlorochromatii* envelope, Cag_2037 is predicted to bind an elongated cellulosome protein Cag_2038, (5) where it will weakly degrade alginate in proximity. (6) Cag_2037 further localizes among the central bacterium outer envelope via its interacting RTX- region [**43**]. As an outstretched protein it will be one of the first to encounter the central bacteria, degrading the alginate capsule at contact sites with the epibiont to permit free diffusion of metabolites and associated proteins between partners. Proteins Cag_663 (red) and Cag_665 (blue) are (1) both upregulated during consortia growth and sulfide limitation, (2) where they likely become excreted in consortia-partnership with a unique TolC (Cag_664) stabilized by SapC (Cag_662) and HlyD (Cag_1507). (3) External contact with Ca^2+^ folds each C-terminal autotransporter into a rigid structure (4) that can successfully reach, penetrate, and enter the central bacterium. (5) Both complete proteins are thus found among the central rods sub-cellular envelop and cytoplasm, delivering the unique C- terminal virulence-factor-associated twisted β-sheet with hydrophobic cleft or laminated β-sheet of Cag_663 and Cag_665, respectively, to the partnering bacterium. Schematic depiction partially built utilizing BioRender.

During sulfide limitation, the giant Cag_663 and Cag_665 proteins are also highly expressed in the epibiont. Via their extended C-terminal repetitive parallel β-helix spike domain reminiscent of T6SS, they may be exported unfolded and C-terminus first through the *sap*-like T1SS of Cag_662, Cag_664 and Cag_1507. Binding of multiple Ca^2+^ in the intercellular space leads to folding into rigid filamentous spikes spanning 200 nm. Because of their homology to autotransporting proteins [**25,26**], these domains are subsequently expected to penetrate the central bacterium, resulting in localization of Cag_663 and Cag_665 proteins either in the central bacterium cell envelope or cytosol. Having high structural homology to proteins predicted for MMB, the active transport of these large proteins may in itself be instrumental for maintaining prokaryotic multicellularity, as similarly gigantic proteins are anticipated to be associated to bacterial adhesion [**52**]. Moreover, the complete localization of both unique N-terminal virulence-factor-domains entirely within the central bacterium cytosol may cause the aggregated subcellular arrays observed by TEM in ‘*Ca.* S. mobilis’ (Fig. S7, [**15,73**]).

These bacterial consortia represent the most highly differentiated bacterial multicellularity across all recorded interactions [**4**], but to date, the specific molecular mechanisms underlying such interactions have largely remained elusive. While RTX-type proteins of bacterial pathogens are considered evolutionarily old [**48,49,51**], our results indicate they are also involved in mutualistic bacteria-bacteria interactions, potentially underlying bacterial commensal relationships as well. In addition, canonical scaffoldin and dockerin modules so far known from *Clostridia* and *Bacteroidia* also evolved in the distant bacterial phylum *Chlorobiota*, enabling intercellular transfer and extracellular exposure of its alginate lyase. Similar to polymorphic toxin systems involved in negative interactions of bacteria with other bacteria or eukaryotic host cells [**27**], the giant multidomain proteins of *Chl. chlorochromatii* likely arose by recombination of different modules. With their giant size, novel combination of fused domains, and limited sequence similarity to described toxin systems, these symbiosis proteins represent a novel means of bacterial interaction, indicating the diversity of giant proteins to be even larger than previously assumed. Our results highlight the need for broader studies of the distribution and adaptation of symbiosis factors for better insights into the general evolutionary principles that shape bacterial interactions [**74**].

## Materials and Methods

### Bacterial strains and plasmids

To date, ‘*Chlorochromatium aggregatum*’ was successfully enriched and cultured [**10**], followed by the isolation and purification of *Chlorobium chlorochromatii* [**14**], which facilitated the comparative growth of this epibiont axenically, and in consortium. In comparison, the central rod of ‘C. aggregatum’, ‘*Ca.* S. mobilis’ could not be isolated in pure culture and in fact may be obligately symbiotic, potentially due to its reduced genome size [**18**]. In addition, dissociation-reassociation studies of consortia cells are not feasible since cells of ‘*Ca.* S. mobilis’ lyse rapidly upon loss of its phototrophic partner. Since standard genetic manipulation studies thus cannot be applied to both partnering bacteria, we followed the growth of epibiotic *Chlorobium* in and out of consortia, and used a combination of omics, protein localization, modeling approaches, and functional assays to study the role of the enigmatic symbiosis proteins in ‘*C. aggregatum’*.

Cultures of ‘*C. aggregatum*’ consortia and its epibiont *Chlorobium chlorochromatii* CaD were grown in triplicate sealed 5 L glass carboys using anoxic sulfide-reduced K3 medium (pH 7.2) [**75**], incubated at room temperature (RT) under continuous 25 µmol quanta m^-2^ s^-1^ surface illumination (determined with a Li Cor LI-250 quantum meter equipped with a LI-200 SA pyranometer sensor; Licoln, NE, USA) of Osram/Ledvance T5 Lumilux L 13W/827 2700 K fluorescent tubes. Neutralized sulfide solution was supplemented as electron donor every second day over a period of one week to enable exponential growth [**13**]. After an increase in OD_600_ of > 0.5, sulfide addition was halted and cultures rapidly entered stationary phase. Under the light limitation induced by large-volume incubation conditions, motile consortia exhibited a pronounced scotophobic response to form an almost pure biofilm on the inner walls of the glass vessels (Fig. E6a; [**75**]) that could be efficiently and rapidly harvested for the subsequent analyses. Growth was monitored via measurements of OD_600_, cellular protein, dry weight, and BChl *c* concentrations (Fig. E6c,d). Briefly, at each timepoint the cultures were mixed via magnetic stir bar, supplied neutralized sulfide if warranted, with a planktonic sample rapidly collected while the anoxic headspace of the carboy was maintained with filtered N_2_ gas prior to returning to illuminated incubator. Planktonic samples were utilized for OD_600_ measurements (Perkin-Elmer Lambda 365+, Rodgau, Germany), while cellular protein was determined for 1 mL of culture cells spun down and rinsed with 50 mM Tris-HCl buffer pH 7.5, prior to alkali hydrolysis using 0.5N NaOH for 1 h at 100°C and final measurement via BCA assay (QPBCA, Sigma-Aldrich, St.Louis, MO, USA) [**76**]. An additional 1 mL washed-cell fraction was utilized to extract total pigments in 0.5 mL 7:2 acetone:methanol overnight, with BChl *c* levels quantified via OD_663_ absorbance (Perkin-Elmer Lambda 365+, Rodgau, Germany; Fig. E6b; [**76**]). Entire 5 L cultures were also decanted at each timepoint, with biomass collected via centrifugation at 10,000 rpm for 10 min intervals, with the biofilm collected separately by rinsing flask with 50 mM Tris-HCl buffer pH 7.5 to easily dislodge consortia prior to similar collection. Biomass was individually placed in previously desiccated and weighted aluminium tins, dried in an oven at 60°C for 2 days, with routine measurements until dry weights stabilized. These four parameters were found to be tightly correlated in our growth experiments (Fig. E6c,d).

*Escherichia coli* M15 (Qiagen, Venlo, Netherlands) *Nal^S^, Str^S^, Rif^S^, Thi^-^, Lac^-^, Ara^+^, Gal^+^, Mtl^-^, F^-^, RecA^+^, Uvr^+^, Lon^+^* was used for plasmid cloning, and *E. coli* XL-1 Blue (Agilent Technologies, CA, USA) *recA1 endA1 gyrA96 thi-1 hsdR17, supE44 relA1 lac* [F’ *pro AB lacI*^q^, *ZΔM15* Tn*10* (Tet^r^)] employed for the expression of recombinant proteins.

Three plasmids, pQE2037, pQE0663 and pQE0665 were constructed for the heterologous expression of recombinant symbiosis proteins of the epibiont. *Nco*I-*Bam*HI restriction sites were introduced through amplification of the whole epibiont gene Cag_2037 (Acc. No. PZ062969) with primers encoding the restriction sites followed by ligation into pQE60 (Qiagen, Venlo, Netherlands), yielding pQE2037. The plasmid pRep4 (Qiagen, Venlo, Netherlands) constitutively expressed an *in trans lac* repressor, which was used to control the expression of the recombinant protein Cag_2037 in *E. coli XL-1Blue*. Among the gigantic proteins Cag_663 and Cag_665, suitable domains for heterologous expression were identified as putative surface regions using a Kyte and Doolittle plot in ProtoScale with a window size of 9 and a hydropathy score set above 2 [**77**]. *Bam*HI-*Pst*I fragments of the predicted protein surface regions of Cag_663 (Acc. No. PZ062967; nucleotide positions 19,798-20,773) and of Cag_665 (Acc. No. PZ062968; 742-1648) were subsequently cloned into the vector pQE30Xa (Qiagen, Venlo, Netherlands) yielding plasmids pQE0663 and pQE0665.

### Microscopic analysis of consortium morphology

Bright field, phase contrast, transmission and scanning electron microscopy (TEM/SEM) were performed as described previously [**15, 78**], with the following modifications: Consortia were prepared for electron microscopy with two different fixation protocols. Glutaraldehyde was added to the growth medium to a final concentration of 2% v/v (GA fixative) and was used to visualize chlorosomes of the epibiont and the intracellular paracrystalline structure of the central bacterium [**15**] whereas treatment with lysine and ruthenium red (LRR fixative) preserved capsules and surface structures [**79**]. For the latter, consortia were pelleted at RT (1 min, 4,000×g), resuspended in fixative solution A (0.2 M cacodylate, 0.15% w/v ruthenium red, 2% v/v formaldehyde, 2.5% v/v glutaraldehyde, and 75 mM L-lysine acetate, pH 6.9), incubated for 20 min on ice, washed twice (0.2 M cacodylate, with 0.15% w/v ruthenium red, pH 6.9), and resuspended in solution B (composition as solution A, but without L-lysine acetate) for 2 h on ice. After three washing steps, the pellet was treated with OsO_4_ (800 µl washing solution plus 200 µl 5% w/v OsO_4_ aq.), and washed twice in 0.1 M EM HEPES buffer (HEPES 0.1 M, 0.09 M sucrose, 10 mM CaCl_2_, 10 mM MgCl_2_, pH 6.9). For SEM, samples fixed in GA or LRR were spotted on poly-L-lysine pre-treated cover-slips and dehydrated in a gradient series of acetone on ice (10%, 30%, 50%, 70%, 90% v/v), each step lasting for 10 min. Two final washes in 100% acetone at RT completed the series. Samples were critical point dried (CPD300, Leica, Wetzlar, Germany) and sputter coated with gold palladium (SCD 500, Bal-Tec, Wetzlar, Germany). Images were acquired with a field emission scanning electron microscope (Merlin, Zeiss, Jena, Germany) at an acceleration voltage of 5 kV using an Everhart Thornley HESE2 / inlens SE detector ratio of 25:75. For TEM, GA fixed samples were washed twice in 0.1 M EM HEPES buffer, treated with 1% w/v OsO_4_ in 0.08 M EM-HEPES for 1 h at RT, and then twice with 0.1 M EM HEPES buffer. In the following, both GA and LRR fixed samples were immobilized in 2% noble agar and dehydrated in a gradient series of ethanol on ice (10%, 30%, 50%, 70%, 90%, 2 × 100%), each step for 30 min with the exception of the 70% step. For the latter, GA samples were incubated overnight with 2% uranyl acetate (UAc) in 70% EtOH at 4°C. Samples were subsequently infiltrated with LR White resin (Sigma, Los Angeles, CA, USA) with increasing concentrations of LR White in EtOH (50, 66%, 100% w/v) and finally polymerized at 55 °C for two days. Ultrathin sections were cut with an ultramicrotome (Ultracut 7, Leica, Wetzlar, Germany) and counterstained with 4% aqueous UAc (3 min) and 4% lead citrate (10 s). Images were acquired with a transmission electron microscope (Libra 120, Zeiss, Jena, Germany) at an acceleration voltage of 120 kV and at calibrated magnifications. For Cryo-SEM high pressure frozen cells (Leica HPM 100, Wetzlar, Germany) were fractured with a Leica MED020 sublimated for 1-2 min at -95°C, coated with 5nm of platinum and transferred with a cryo shuttle to the cryo stage of the SEM and examined at 0.8-1 kV. Focused Ion Beam (FIB) FIB/SEM tomography utilized high pressure freeze-substituted cells embedded in Spurr’s low viscosity epoxy resin [**80**], and seried sections of ‘*C. aggregatum*’ enabled 3D reconstruction of entire consortia as described following Protocol A [**81**].

### ‘*C. aggregatum’* genome sequencing and analysis

At the outset, circular genomes of *Chl. chlorochromatii* and ‘*Ca.* S. mobilis’ that had been produced from short-reads via Sanger sequencing of small inserts and fosmid clones [**7**], or by Roche 454 GS FLX Titanium sequencing more than a decade ago [**18**] were reevaluated using PacBio Sequel*IIe* long-reads obtained for DNA extracted directly from cultivated consortia. Reads were assembled using Flye (2.9), and freshly annotated with Prokka (1.14) [**82**]. Complete genome comparisons using Mauve [**83**] revealed the presence of additional repeat regions, gene duplications, and transposons compared to the existing genome sequences (Fig. S1). An updated list of pertinent genes and locus tags is provided in Table S1 and S2. The locus tags Cag_663, Cag_665 and Cag_2037 used here for the three symbiosis proteins refer to the original numbers Cag_0614, Cag_0616 and Cag_1919 in [**20**].

Phylogenomic relationships to type strains of either phylum *Chlorobiota* (for the epibiont) or family *Comamonadaceae* (for the central rod) were inferred via multi-loci-sequence analysis (MLSA) phylogenetic trees using RAxML [**84**] based on 222-411 single-copy core genes, dependent on the maximum number detected per genome-group. Differential gene content among members of each group were then further identified using ProteinOrtho groups visualized as upset plots [**85**].

### Predictions of protein domains and functionalities

ORFs of interest [**7,20**] were first screened using BlastP (2.14.0) to determine the similarity to known proteins containing domains involved in molecular interactions or with specific functions. Domains of interest were localized using HHPred [**86**], InterProScan (using InterPro102.0) [**87**], and MotifScan (1.3.0). HHpred is a method for sequence database searching and structure prediction that is more sensitive in finding remote homologs to known proteins than homology searches (BlastP), as it compares aa sequences and 3D structure to various profile Hidden Markov Models (HMMs). InterPro recognizes known domains separately, by classifying them into families and predicting domains and important sites. To classify proteins, InterPro uses predictive models, known as signatures, provided by several member databases that make up the InterPro consortium. HHpred and InterPro thus are complementary and are currently both used in novel protein domain-identification. AlphaFold3 [**88**] was used to predict the three-dimensional structure of each symbiosis protein, using 1500 aa sections with 300 aa overlap of the translated ORF in Geneious Prime (2023.0.2). The sliding window enabled models for the entire ORF to be built in PyMOL (3.0.3) (Fig. 3, S5, S14, S15). The 1500 aa fragments were longer than sizes used in previous modeling of putative large proteins [**41**] and determined to be optimal as they covered entire domains of interest and avoided unnecessary artifacts of self-folding (Fig. S1c). Resulting structures were subsequently analyzed for hydrophobic or electrostaticity in ChimeraX (1.8), cation binding via PeSTo [**89**], and structural homology to documented proteins via Foldseek (9-427) [**90**]. Different metrics were employed to assess the confidence of the predictions. The predicted TM score (pTM) gives an overall structural accuracy of the predicted protein structure compared to a template structure, with a higher pTM score indicating a more accurate prediction in terms of overall topology. The predicted local Distance Difference Test (plDDT) is a residue-level metric that indicates the confidence of the prediction at each individual aa position. A high plDDT value suggests that the model is highly confident in the predicted position of that specific residue. Regions with lower plDDT scores are less well-defined in the prediction. The interface predicted TM-score (ipTM) focuses specifically on the accuracy of the predicted protein-protein interactions, particularly at the interface between different protein chains. A high ipTM score indicates that the predicted interaction interface is accurate, while a low ipTM suggests that the model may not show interaction confidently. The B-factor refers to the atomic displacement parameter, typically representing the extent of atomic motion or disorder. This helps identify areas of flexibility or potentially disordered regions in a protein. Electrostaticity refers to the electrical interactions between charged aa residues. It is useful for understanding the distribution of charge, to identify areas of positive or negative charge, and to assess the potential for interactions with other molecules.

### Analysis of protein secretion systems

The origin of 32 *Chl. chlorochromatii* ORFs encoding components of T1SS (Table S4) were analyzed via amino acid homology searches in ConSurf [**91**]. In addition, four T1SS components detected in ‘*Ca.* S. mobilis’ were analyzed. In ConSurf, aa sequences were compared via a HMMER search algorithm to determine 150 diverse homologs from UniRef90 (<u>Uni</u>Prot <u>Ref</u>erence Clusters, with representative sequences filtered to <u>90</u>% identity to reduce redundancy), with identities set between minimal and maximal values of 30-98%. The top 5 BlastP RefSeq-Select hits which excluded uncultured/ environmental sample sequences were added to each list and aligned utilizing MAFFT (v.7) [**92**]. A Bayesian best-fit model calculated scores of conserved sites, and the resulting alignments were subsequently utilized for phylogenetic Neighbor Joining [**93**], comparing conserved aa and gap-free sites by the JTT substitution model [**94**] and using 1000 bootstrap replicates. Phylogenetic trees were visualized in iTOL (7.2.1) [**95**] rooted to midpoint. *Chl. chlorochromatii* secretion system genes found to have close homologs among other type strains of *Chlorobiota* via BlastP RefSeq-Select hits and/or ConSurf were declared as vertically inherited since they could be found in diverse members of the phyla (Table S4). Genes found only in *Chl. chlorochromatii* and potentially uncultured MAGs, with nearest homologs identified among taxa outside of the phylum were considered recently acquired via HGT and not considered ancestral vertically inherited genes. Here, uncultured MAGs could represent strains of this species, not free-living examples, and thus may also be consortia-forming.

### Transcriptomic analysis

‘*C. aggregatum’* consortia or axenic epibiont *Chl. chlorochromatii* cultures in the mid-exponential or stationary phases of growth were harvested from the illuminated inner surface of each glass bottle. 15 ml of cell suspension were mixed with an equal volume of RNAprotect® (Qiagen, Germany), rapidly concentrated by centrifugation (1 min 20,000×g), the supernatant discarded, cells rapidly frozen in liquid nitrogen, and stored at -80°C. Bulk RNA was extracted via bead beating with sodium dodecyl sulphate and phenol/chloroform purification [**96**], an RNA cleanup using RNeasy® Mini kit (Qiagen, Venlo, the Netherlands) which included on-column DNAse digestion, followed by an additional DNAse I treatment (Thermo Fisher, MA, USA), prior to excess rRNA removal via an Ribo-off Depletion kit V2-Bacteria (Vazyme Biotech, Jiangsu Province, China). RNA concentrations were monitored by Bioanalyzer with RNA 6000 Pico Reagents (Agilent, CA, USA), before cDNA libraries were created by a TruSeq® Standard mRNA Library Prep Kit for Illumina® (San Diego, CA, USA), agarose gel electrophoresis confirmation, and quantified by Qubit analysis (Thermo Fisher, MA, USA). The Illumina® NextSeq 2000 platform using P3 Reagents (San Diego, CA, USA) was employed for sequencing, running 50 cycles of single reads. Output BCL files were demultiplexed, the barcodes trimmed and sequences converted into FASTQ files via bclfastq2 (2.15.0). The resulting datasets comprised an average of 6.5 Mio sequences of 2.5 Bbp summed length. FastQC (0.12.1) analysis revealed that sequences had Phred scores >32 and thus sufficient quality. Reads were mapped to the genomes of either *Chl. chlorochromatii* or ‘*Ca.* S. mobilis’ via HiSat2 (2.2.1), converted to .bam files with SamTools [**97**], and a count matrix produced using SubRead (2.0.6).

Relative gene expression was then analyzed by DeSeq2 (1.40.2) in RStudio (2023.06.2) and visualized through heatmaps or volcano plots, employing pheatmap and ggplot, respectively. The p-values determined for difference between samples were assessed via the Wald test applying the Benjamini and Hochberg method [**98**]. Normalized transcript counts were individually Z-score transformed to a scale between 0-100, so as to track patterns of expression in primary figures (Fig. 3b-d). Genes which contained significant differences between exponential and stationary growth phases, or between symbiotic states, were graphed as groups. To determine transcript coverage over large ORFs and to delineate polycistronic features, the epibiont-specific .bam files generated by HiSat2 for each growth phase were mapped against the complete genome of *Chl. chlorochromatii* using the Burrows-Wheeler Aligner [**99**]. The integrative genome viewer [**100**] then allowed a comparison of all transcriptomes to one another. A network analysis of epibiont transcripts was also constructed with BioLayout (3.4) [**45,101**], utilizing a 2.2× inflation for the Markov clustering (MCL) algorithm. Calculations were based on the correlations between standardized transcript counts of the different genes under different growth and symbiotic conditions. Each individual gene could be represented as a node (sphere), where edges (lines) indicated correlations above a 0.85 Pearson correlation threshold, with lengths of edges inversely proportional to the strength of correlation. Clusters below 3 nodes were filtered out, with 4 primary clusters listed in supplements.

### Expression of recombinant proteins for antibody production

Recombinant proteins were expressed in *E. coli* XL-1 Blue grown at 28°C in Terrific Broth Medium [**102**] with the addition of ampicillin (100 µg·ml^-1^) and tetracycline (75 µg·ml^-1^) and, in the case of recombinant Cag_2037, with the addition of kanamycin (50 µg·ml^-1^). Overproduction of recombinant proteins was induced by isopropyl-ß-ᴅ-1-thiogalactopyranoside (IPTG) at an optical density of OD_595_ = 0.7. After 50 to 72 h of induction, recombinant proteins were purified under denaturing conditions with the BugBuster kit (Novagen, Darmstadt, Germany) using inclusion body protocols, followed by 1 h shaking at 4°C in 8 M urea buffer (100 mM NaH_2_PO_4_, 10 mM Tris-HCl, pH 7.2). After centrifugation (30 min at 20,000×g, 8°C), the supernatant containing the protein fragments of Cag_663 or Cag_665 were purified via Ni-nitrilotriacetic acid HisTrap FF 1 mL columns for purification, whereas recombinant protein Cag_2037 required a HiPrep 16/60 Sephacryl S-300 HR column (GE Healthcare, Buckinghamshire, Great Britain). Antisera were manufactured by Eurogentec (Seraing, Belgium), purified and employed as primary antisera in the immunoblots, immunofluorescence and immunogold analysis.

### Cell fractionation

To obtain different cell fractions from the negative control *E. coli* XL-1 Blue, 2 g of cells were pelleted and washed with 50 mM Tris-HCl buffer pH 7.5, prior to resuspension in 50 mM Tris-HCl buffer with 40% (v/v) sucrose. Lysozyme (0.6 g/ml) and EDTA (5mM) were added to digest cell walls for 30 min at 37°C. The spheroplasts generated were spun down (20 min at 1700×g, 4°C) and the periplasmic fraction collected as supernatant. After resuspending the pellet in 1 ml of Tris-HCl buffer, spheroplasts were lysed by osmotic shock. The lysate was homogenised, sonicated (Bandelin Sonopuls HD 2070) and centrifuged (20 min at 20,000×g, 4°C) to remove debris. Ultracentrifugation (30 min at 160,000×g, 4°C) generated the membrane fraction as a pellet, which was solubilised in 50 mM Tris-HCl pH 7.5, 5 mM MgCl_2_ and 10 % (v/v) glycerol, and the supernatant collected as the cytoplasmic fraction.

A modified protocol was developed for the fractionation of phototrophic consortia and epibiont cells. One g of cell pellet was washed with phosphate buffer saline (PBS) (137 mM NaCl, 2.7 mM KCl, 10 mM Na_2_HPO_4_, 1.8 mM, KH_2_PO_4_, pH 7.4) and resuspended in 0.3 M Tris-HCl buffer containing 40 % (v/v) sucrose. Ethylenediaminetetraacetic acid (EDTA) (1 mM) and lysozyme (1 mg·ml^-1^) were added to the buffer and the periplasmic fraction was collected as described above. After the spheroplasts were lysed by osmotic shock the cell pellet was resuspended in 10 mM Tris-HCl pH 7, 7.5 mM EDTA with 0.2 mM dithiothreitol (DTT), prior to sonication and centrifugation (as above). Cell lysis was confirmed by microscopy of the treated cells. Cell membranes were collected through extended ultracentrifugation (3 h at 160,000×g, 4°C) and the cytoplasmic fraction was collected as the supernatant. Cell membranes were solubilised in 50 mM Tris-HCl (pH 8), containing 2% (w/v) Triton X-100, 10 mM MgCl_2_, 100 mM NaCl and 10% (v/v) glycerol.

### Determination of symbiosis protein sizes

The size of native Cag_663 and Cag_665 proteins could not be determined by conventional sodium dodecyl sulphate polyacrylamide gel electrophoresis (SDS-PAGE), either 9% w/v, or gradient gels with 4 - 15% w/v polyacrylamide, since theoretical peptide sizes would not enter such matrices, and attempts revealed unresolved protein bands at leading edge of loading wells. Moreover, at 3741.4 kDa, Cag_663 surpasses all largest available standard ladder markers by size (∼500 kDa), and would need alternative comparative markers. Therefore, an agarose stabilized polyacrylamide gel was employed [**103**] with modified running buffer (50 mM Tris-NaOH, 0.384 M glycine, 0.10% w/v SDS) [**104**], was run at 200 V for 200 min in a water-cooled system, and afterwards stained and destained as described [**103**]. *In lieu* of a standard ladder, titin protein of 3,000-3,200 kDa (N2B and N2BA) and Myosin heavy chain of ∼233 kDa was extracted from CD1 *Mus musculus* cardiac tissue whole lysate in sample buffer [**105**] and heat treated 2 times for 5 min at 100°C for cell lysis and solubilization. This lysate was run against lysates of axenic *Chlorobium* and consortia treated similarly, as these large mammalian proteins would segregate from all other smaller biomass and act as reference material of similar weight in denaturing gel electrophoresis experiments.

Excised bands were destained using 50mM Triethylammoniumbicarbonat (TEAB) in 40% acetonitrile for 30 min at RT, then dried with 100% acetonitrile. For reduction, samples were overlayered with 20 mM dithiothreitol (DTT) in 50 mM TEAB and incubated at 56°C for 60 min. Alkylation followed with addition of 50 mM iodoaceamide (IAA) for 30 min at RT. Excessive IAA was removed by successive washing for 30 min with 50 mM TEAB and a 50 mM TEAB in 40% acetonitrile afterwards. The gels were again dried with 100% acetonitrile before overnight incubation at 37°C with 2ng/µL Trypsin in 50 mM TEAB with 5% acetonitrile. The digested peptides were extracted from the gel using (a) 0.2% TFA in water, (b) 0.2% TFA in 40% acetonitrile, and (c) 0.2% TFA in 100% acetonitrile. During every extraction step, the samples were shaking at 500 rpm for 30 minutes. All collected extraction volumes were united and dried in a vacuum concentrator prior to LC-MS (below).

For proteomic analysis, 2 g of *Chlorobium* biomass (axenic epibiont or consortia) was pelleted (5 min at 10,000 *g*, 4°C), washed with 50 mM Tris-HCl buffer pH 7.5, pelleted again, and resuspended in lysis buffer of 1% SDS 4.3 mM Tris-HCl, pH 8.8, 4.3mM EDTA, and 8ug/mL leupeptin. Cells were lysed via French-Press cell, cellular debris was removed (20 min at 20,000 *g*, 4°C), and lysates were digested with different enzymes, including Trypsin/LysC, Chymotrypsin, or combined digestion with pepsin and trypsin. For Trypsin/LysC- and Chymotrypsin digest, the lysates were solved in 50 mM TRIS-HCl (pH 8) containing 4% SDS, 5 mM EDTA, and 1 mM EGTA before solubilization of the sample in the Bioruptor (10 cycles, 30s on/ 30s off). For reduction and alkylation, the samples incubated for 1 hour at 56°C with 5 mM DTT and 15 min at room temperature with 10 mM chloroacetamide (CAA), respectively. Trypsin/LysC was added with a protein:enzyme ratio of 25:1 and chymotrypsin with a ratio of 15:1. For the combined digest with pepsin and trypsin, the lysates were resuspended in 1% TFA and incubated 4 hours at 37°C in the presence of pepsin. Afterwards, the samples were diluted with the same volume of 1M TEAB with 1% SDS and 5 mM EDTA before reduction, alkylation and addition of trypsin (25:1). Each digest incubated overnight at 37°C. After acidification of the samples with 1% formic acid (FA), a peptide clean-up was performed using 50 µg SP3 beads (Preomics, Martinsried, Germany), and brought to 95% acetonitrile. Kept shaking overnight, supernatant was removed and the beads were washed two times with 100% acetonitrile. Samples were finally dried and eluted first in 2% DMSO, and then in 0.1% FA. The united elution fractions were dried in the vacuum concentrator prior to LC-MS (below).

LC-MS of all digested protein samples began with 3min sonication in 0.1% FA. The solved peptides were analysed with a TimsTOF Pro (Bruker, Bremen, Germany) coupled to a Evosep One (Evosep, Odense, Denmark). After Evotip loading, the samples were injected and trapped with an Endurance C18 column (8 cm x 100 µm, Evosep) heated to 30°C and separated with the pre-build 60 SPD-method (Evosep) using 0.1% FA in water for Buffer A and 0.1% FA in acetonitrile for Buffer B. Peptides were ionized using a Captive Spray 2 Emitter (inner diameter 20 μm, Bruker) and a Capillary Voltage of 1600V. Spectra were acquired in DDA-PASEF Mode. The Ion Mobility window (1/k^0^) was set to 0.7-1.4 Vs/cm^2^ with a ramp and accumulation time of 100 ms. The mass scan range was set to 100-1700 m/z. Precursor were fragmented with a collision energy of 10 eV and analysed in 4 PASEF ramps with a total cycle time of 530 ms. The intensity threshold was set to 2500 and the target intensity to 20000. LC-MS runs were analysed using Peaks v3.5 (Bioinformatics Solutions). For each sample, the digestion enzyme was defined (trypsin, chymotrypsin) for a specific (gel analyses) or semi-specific (multi-enzyme) digest mode and compared to a built database of ‘*Ca.* S. mobilis’ and *Chl. chlorochromatii* (from genome sequencing) or *Mus musculus* (Uniprot). Peptides with a minimum length of 6 and maximum 45 aa, a fixed carbamidomethylation on cysteines (+45.99) and variable oxidation on methionines were characterized. From the resulting lists, only top proteins of a protein group were kept and common contaminants were filtered out. Final coverages were compared to *in silico* digests determined via Expasy [**77**].

### Western and dot blot analyses

The expression and subcellar localisation of the Cag_2037 protein was analysed by Western blots. 2 ml of cell suspension (grown to an OD_595_ of 0.7) was boiled for 5 min in Laemmli sample buffer (Serva Electrophoresis, Heidelberg, Germany) [**106**], the debris centrifuged (20,000×*g* for 5 min), and 10 µg soluble protein separated by SDS PAGE (9% w/v). Proteins separated were blotted onto polyvinylidene difluoride (PVDF) membranes (Carl Roth, Karlsruhe, Germany) in a Bio-Rad system at 50 V for 3 h using transfer buffer (25 mM Tris-NaOH, 192 mM glycine and 20% v/v methanol). The membrane was blocked overnight, incubated for 1 h with the primary antibody against the Cag_2037 protein, and then 1 h with a horseradish peroxidase (HRP) conjugated secondary antibody. Due to the much larger molecular size of the Cag_663 or Cag_665 proteins, and since Western blots revealed no evidence of small-sized fragments for either protein, their expression had to be analysed via dot blot. Per sample, 100 µg protein in 100 µl 1 M Tris-HCl (pH 7.2) was loaded onto PVDR membranes using a Bico BRL device (Life Technologies, CA, USA). After three washes with PBS, membranes were dried for 1 h at RT, incubated for 1 h with the primary antibody against the Cag_663 and Cag_665 proteins, and then 1 h with HRP conjugated secondary antibody. For detection, the chemiluminescent HRP substrate system Lumi-Light (Roche Diagnostics, Basel, Switzerland) was used with an incubation time of 5 min after which images were captured (Fujifilm LAS-3000, Japan).

### Super-resolution immunofluorescence microscopy

Consortia or axenic *Chl. chlorochromatii* epibionts were collected on black polycarbonate filters (0.2 µm pore size; Merck Millipore, MA, USA) and washed three times with PBS prior to applying a modified immunofluorescence protocol [**107**].

While cells for Cag_2037 imaging were fixed in 4% (v/v) paraformaldehyde overnight at 4°C, the Cag_663 and Cag_665 products were detected in cells fixed with 3.5% (v/v) formaldehyde and 0.008% (v/v) glutaraldehyde in PBS at RT for 4 h. Subsequently, filters were washed three times with PBS, incubated in 0.2% (v/v) Triton X-100 for 1 h at RT, washed again, subjected to PBS containing lysozyme (100 µg·ml^-1^) and 5 mM EDTA for 1 h at RT, and then treated with 0.5% (w/v) blocking reagent (Boehringer, Ingelheim, Germany). The first antibody was applied in PBST (PBS supplemented with 0.1% v/v Tween 20) during an overnight incubation of filters at 4°C, followed by three washes with PBS and then 1 h of incubation with secondary antibodies. For total internal reflection fluorescence (TIRF) microscopy, Alexa Fluor 488 nm conjugated goat anti-rabbit antibody was used, whereas for direct stochastic optical reconstruction microscopy (dSTORM), an Alexa Fluor 647 nm coupled anti-rabbit antibody (Agilent Technologies, CA, USA) was employed, allowing a higher resolution (80 nm). Three washing steps with PBS preceded a counterstaining of all cells with 4’,6-diamidino-2-phenylindole (DAPI; 100 ng·ml^-1^, 10 min), followed by three final washes with PBS and sectioning of the filters with a sterile scalpel. Bacteriochlorophyll (BChl) *c* constitutes the main photosynthetic pigment of the green sulfur bacterial epibionts where it is densely packed in molecular aggregates housed in specialized antenna structures (chlorosomes) which are bound to the inner surface of the cytoplasmic membrane [**14,15**]. This particular arrangement results in a distinct infrared fluorescence, encompassing the entire epibiont cell, with a peak emission at 775 nm upon excitation with blue light [**108**]. This specific emission wavelength of BChl *c in vivo* allowed us to study the co-localization of epibiont cells, the DAPI-stained cells, and the immunofluorescence signals. In the absence of autofluorescence, the fluorescence membrane dye FM4-64FX (Thermo Fisher, Waltham MA, USA) was employed for staining of *E. coli* XL-1 Blue cells that served as positive controls.

For TIRF microscopy, filter sections were mounted on glass slides in TIRF buffer (20 mM Tris-HCl pH 8, 0.5% w/v N-propylgallate and 90 % v/v glycerol) and sealed under a cover slip (High Precision Glass 170 ± 5 µm, Carl Roth, Karlsruhe, Germany) with nail varnish. Employing a Nikon Eclipse Ti high resolution microscope (Nikon, Tokyo, Japan), images were captured with an Andor camera DU-897X-4669 using excitation filter 442/46 and emission filter 525/45 for Alexa Fluor 488 detection, excitation filter 430/24 and emission filter 760/50 for the detection of BChl *c* autofluorescence, as well as excitation filter 377/50 and emission filter 442/46 for the detection of DAPI. Sets of z-stack images of 0.1 µm distance were analysed and deconvoluted using NIS-Elements AR (4.13.01). For additional dSTORM analysis, filter sections were placed in a glass bottom dish (MatTek Corp., USA) with 4 µl dSTORM buffer: A 7 µl glox solution (containing 0.28 g·ml^-1^ glucose oxidase and 0.017 g·ml^-1^ catalase in distilled H_2_O), combined with 200 µl ‘buffer A’ (10 mM Tris-HCl, 50 mM NaCl, pH 8), 620 µl ‘buffer B’ (50 mM Tris-HCl, 10 % w/v glucose, 10 mM NaCl, pH 8), and with 70 µl 1 M cysteamine [**109**]. Each filter was covered by a circular agarose pad (0.8% w/v) and with 12 µl of dSTORM buffer, covered with a coverslip and sealed with vaseline. The cells mounted in the glass bottom dish were then illuminated and microphotographed using the Nikon microscope similar to TIRF. In dSTORM mode, visualization of protein-bound Alexa Fluor was achieved with either 100% 647 nm or 10% 405 nm fibre-laser light intensity to observe total internal reflection geometry. A frame rate of 30 ms, Em gain of 30, and 10 MHz at 14 bit-gain conversion allowed for image capture at 80 nm resolution using the NIS-Elements AR (4.13.01) software.

Cells of *E. coli* XL-1 Blue expressing each protein (fragment) served as positive controls, whereas *Chl. chlorochromatii* and ‘*C. aggregatum*’ were used as negative controls which were treated with pre-immunisation sera and the secondary antibody only. The chlorosomes lining the inner membrane surface were used to delineate cells of the green sulfur bacteria and were visualized by the autofluorescence of chlorosome BChl *c* as described above. *E. coli* cell membranes were selectively stained with FM4-64FX (Thermo Fisher in Waltham, USA) using excitation and emission wavelengths 558 nm and 634 nm, respectively.

### Immunogold electron microscopy

Intact ‘*C. aggregatum*’ consortia were fixed in acetone containing 0.3% w/v uranyl acetate (Leica EM HPM 100, Vienna, Austria). Each sample was then cooled to -90°C for 28 h, slowly heated to -60°C over 6 h, kept at -60°C for 4 h, and warmed to -50°C for 2.5 h. All subsequent manipulations were performed at -50°C. Samples were washed for 15, 30 and then 45 min with pure acetone, submerged in a graded series of HM20 (Electron Microscopy Sciences, Hatfield, USA)/acetone mixtures (25% HM20 w/v for 1.5 h, 50% HM20 for 2 h, 75% HM20 for 2.5 h and pure HM20 for 16 h, 3.5 h, and 4 h) and subsequently polymerised with ultraviolet light for 48 h. Finally, the temperature was raised to 20°C over 14 h and samples completely polymerised for 22 h. Samples were cryosectioned and washed twice with PBS containing 50 mM glycine. The cryosections were placed in blocking buffer (0.5% w/v BSA, 0.05% w/v gelatine, 0.01% v/v Tween 20 in PBS) for 1 h and incubated over night at 4°C with the primary antibody (5 µg·ml^-1^). Samples were then washed six times for 5 min with blocking buffer (PBS containing glycine; Ampuwa, Fresenius, Germany), incubated for 90 min at RT with the secondary anti-rabbit antibody conjugated with 6 nm gold particles (Dianova, Hamburg, Germany), and finally washed twice for 5 min with blocking buffer. A total of 38 immunogold labelling electron micrographs were analyzed, including negative controls. Gold particles were considered localized in the cell envelope if they were observed within a distance of ≤ 25 nm from the membrane (the approximate length of an antibody complex). The statistical significance of the labeling compared to the negative controls was analyzed by an unpaired t-test, whereas the average number of immunogold particles per µm^2^ area was calculated for each subcellular location and statistically evaluated by t-tests employing the R code command pairwise.t.test in Rstudio (2023.06.2).

### Detection of alginate capsules in phototrophic consortia

Alginate biosynthesis genes were screened and discovered in the ‘*Ca.* S. mobilis’ newly annotated genome sequence (Fig. S2a, Table S2, Fig. S13h), while absent in the *Chl. chlorochromatii* genome (Table S1). The presence of alginate as a predominant component of the consortia was confirmed by formic acid hydrolysis and identification of mannuronic and guluronic acid components via gas chromatography coupled mass spectrometry (GC-MS) [**110**]. Detection limits were established by dissolving different amounts of Na-alginate (Sigma, Los Angeles, CA, USA) in triplicates (0, 0.01, 0.05, 0.1, 0.5, 1.0, 3.0, 5.0, 7.5, 10, 15, or 20 mg) to 1.5 ml of 90% v/v formic acid, incubation of samples at 100°C for 6 h prior to an addition of 6.5 ml ddH_2_O and 2 h of incubation further hydrolysis. The samples were evaporated in a speed-vac, resuspended in 1 ml H_2_O, and 5 µl subsamples were supplemented with 100 µl 1 % w/v ^13^C ribitol, dried and derivatized with methoxyamine hydrochloride and N-methyl-N-(trimethylsilyl)-trifluoracetamide for GC-MS analysis as described [**111**]. Similarly, pelleted biomass of stationary phase consortia or axenic *Chl. chlorochromatii* epibiont cultures were hydrolyzed and analyzed. Standard curves for correlations between wet and dry weight biomass ratios were determined (Fig. E6d), and final measurements were calculated as combined hydrolyzed alginate monomers (mannuronic and guluronic acids measured as pentakis-O-(trimethylsilyl)-, o-methyloxyme derivatives; Fig. 5d) per mg of dry biomass. To determine the structural function of the alginate capsule in intact consortia, ‘*C. aggregatum’* cultures were supplemented with 400 ng·ml^-1^ alginate lyase (A1603, Sigma-Aldrich, St. Louis, USA) and grown for 96 h. Samples from different timepoints were fixed in 2% paraformaldehyde, washed twice with PBS, and intact consortia enumerated via counting chamber.

### Alginate lyase functional assays

The recombinant protein Cag_2037 was expressed for 72 h and purified under denaturing conditions following the BugBuster protocol (Novagen, Darmstadt, Germany). Two additional washing steps were followed by solubilisation of the purified protein in 8 M urea buffer and shaking at 4°C overnight. After ultracentrifugation (45 min at 160,000×g, 8°C), supernatant protein content was determined using Lowry Assay (Bio-Rad, California, USA) and purity screened by 9% SDS PAGE. Cag_2037 was refolded via multi-step dialysis at 4°C in a Spectrum Labs unit (Los Angeles, USA) using buffer A [10 mM 4-(2-hydroxyethyl)-1-piperazineethanesulfonic acid, pH 7.9, 300 mM KCl, 10 mM MgCl, 5 mM CaCl, 10% v/v glycerol and 100 µM dithiothreitol (DTT)] and then buffer B [100 mM 2-amino-2-(hydroxymethyl)propane-1,3-diol, pH 7.5, 50 mM KCl, 50 mM NaCl, 10 mM MgCl_2_, 5 mM CaCl, 1% v/v glycerol and 100 µM DTT] and then concentrated by placing the 100 kDa pore-size dialysis tubing in Spectra/Gel^®^ absorbent (polyacrylate-polyalcohol; Spectrum Labs).

The refolded protein was tested for alginate lysis via an OD_548_ thiobarbituric acid (TBA) assay that measures formation of the alginate degradation products mannuronic acid and guluronic acid [**112**]. A Na-alginate solution (0.66 g of Na-alginate; Sigma, Los Angeles, CA, USA; dissolved in 300 mM NaCl, 50 mM NaH_2_PO_4_) was mixed in equal portions with either 1.5 mg·ml^-1^ refolded Cag_2037 protein or known alginate lyase (A1603; Sigma-Aldrich, St. Louis, MO, USA) and incubated for 48 h. For subsequent chemical analysis, 0.1 ml of each enzyme assay was mixed with 0.125 ml of 0.025 N H_2_IO_6_ in 0.125 N H_2_SO_4_, incubated 20 min at RT, then 0.25 ml of 2% NaAsO₂ v/v in 0.5 N HCl was added, incubated for 2 min, followed by an addition of 1 ml of 0.3% w/v thiobarbituric acid, and by a final heating step at 98°C for 10 min. Before analysing the degradation products, proteins were removed by addition of an equal volume of chloroform, the two phases separated by centrifugation at 10,000×g for 10 min, and the upper phase collected. 50 µl of sample or standard were injected on a Dionex ultimate 3000 system (Thermo Scientific Inc., USA) coupled to a Bruker MicroTOF QII mass spectrometer (Bruker Daltonik GmbH, Germany) equipped with an electrospray ionisation interface (ESI). Separation was carried out on a Hi-Plex Na column (300×7.7 mm; Agilent, Germany) at a constant temperature of 85°C with a flow rate of 0.3 ml·min^-1^ of 0.5 mM Na-formate. The MS was operated in negative ESI mode with 3 Hz data acquisition of full scan mass spectra between 90 and 1178 m/z. Extracted ion chromatograms of the exact masses of M-H ion (monosaccharide 193.0399 m/z, disaccharide 369.0768 m/z) using commercially available standards (Carbosynth Limited, Compton, UK) showed a retention time of 12.69 min for the monomer and 11.46 min for the dimer. Mannuronic acid and guluronic acid had the same elution time.

## Database analysis

Sequence titles and isolation sources of small sub-unit (SSU) rRNA sequences in the SILVA database [**2**] were searched for matches with lists of predefined descriptive keywords to determine the total numbers of mutualistic or pathogenic sequence types. For retrieving mututalistic microorganisms, the list comprised the keywords “symbiont”, “symbio”, “mutualistic”, “mutual”, “commensal”, “cooperation”, “cooperate”, “syntrophy”, “syntroph”, “endophyte”, “beneficial”, “benefit”, “partner”. For retrieving pathogens and microorganisms causing other types of negative interactions, the keywords “pathogenesis”, “pathogen”, “parasitic”, “parasite”, “predator”, “illness”, “disease”, “infect”, “infectious”, “clinical”, and “virulence” were employed.

## Acknowledgements

Illumina and PacBio platforms were run by the DSMZ sequencing facility and proteomics analysis conducted by the HZI proteomics core facility for translational proteomics (CFTP) of Prof. Dr. Lothar Jänsch. We would like to thank Ina Brentrop and Sylvia Dobler for excellent technical support in several electron microscope sample preparations. For her help establishing immunofluorescence protocols for consortia, we are indebted to Sabrina Willems. We would also like to thank Isabel Schober for conducting the NCBI and Silva database search, generating current numbers regarding bacteria annotated associated with a symbiotic lifestyle. Philipp Halama is appreciated for his assistance with genome comparisons and upset plot generation. This research was funded by a grant of German Research Foundation (DFG OV 20/10-2).

## Data Availability

PacBio sequenced and circularized genomes of *Chlorobium chlorochromatii* CaD and ‘*Ca.* Symbiobacter mobilis’ M will be available with Genbank accession numbers JBRGNI000000000 and JBRGNJ000000000, respectively. Symbiosis genes Cag_663, _665, _2037, and _2038 have accession numbers PZ062967- PZ062970. The mass spectrometry proteomics data has been deposited to the ProteomeXchange Consortium via the PRIDE partner repository (dataset and identifier on request).

## Supplemental Table Legends

**Table S1. Analysis of genes of *Chlorobium chlorochromatii* and other *Chlorobiota*. (A)** 222 single-copy core genes used for a MLSA tree to differentiate *Chlorobiota*. **(B)** ORFs shared among all members of *Chlorobiaceae*, include most (22) anoxygenic photosynthesis genes. **(C)** List of 356 ORFs unique to epibiotic *Chl. chlorochromatii* but absent from all other known type strain genomes of the phylum *Chlorobiota*. RAST annotations of the newly sequenced genome are provided by ascending locus-tag number. Symbiosis genes targeted in the present work are highlighted in dark green.

**Table S2. List of annotated genes of epibiotic *Chlorobium chlorochromatii* with potential implications for symbiosis, and of housekeeping genes in the focus of our transcriptomic analysis.** Current locus tags and RAST annotations of the newly sequenced and circularized genome are provided along with original locus tags and grouped by physiological role. Unique genes to *Chl. chlorochromatii* highlighted in light green, with the three symbiosis genes studied marked in dark green.

**Table S3. Domains of symbiotic proteins of the epibiotic *Chlorobium chlorochromatii* identified by HHPred.** Individual fragments of Cag_2037, Cag_2038, Cag_663, Cag_664, Cag_665 were examined and the highest probability hits for each region are listed.

**Table S4. Similarity and deduced origin of secretion system components of *Chl. chlorochromatii* and ‘*Ca.* S. mobilis’.** Annotated ORFs were translated and analyzed in BlastP RefSeq-Select to exclude uncultured/ environmental sample sequences to look for the top 5 nearest orthologs via aa sequence similarity (given in %) to those in the database for each secretion system related gene. ORFs found in 6 distinct operons are framed in grey and their structure depicted in Fig. S2C. Genes unique to *Chl. chlorochromatii* are highlighted in light green, with nearest homology listed in pink.

**Table S5. Comparisons of AlphaFold3 predicted structures to those held in databases. (A)** Predicted structural regions were aligned in FoldSeq to structures available in databases (AlphaFold, Proteome, Swiss-Prot, UniProt50, BFMD, CATH50, GMGCL, MGnify-ESM30 and PDB100). As no experimentally verified structures similar to either giant protein were found, the top 10 ORFs with predicted structures are listed. In contrast, each domain of Cag_2037 could be matched to the 5 experimentally solved structures listed with decreasing probability, followed by top 5 hits to predicted models. **(B)** Specific homologous regions found in multicellular magnetotactic bacteria. **(C)** Putative cation binding sites annotated (PeSTo).

**Table S6. Network analysis of changes in *Chl. chlorochromatii* transcriptomes upon transition between exponential and stationary phases of growth and in either axenic or in consortia cultures. (A)** Complete list of 2152 annotated ORFs, including the 2096 protein-coding genes, **(B)** ORFs in network cluster 1, **(C)** ORFs in network cluster 2, **(D)** ORFs in network cluster 3, and (E) ORFs in network cluster 4 (compare Fig. S8b). Current locus tags highlighted as *Chl. chlorochromatii* specific (dark green) or *Chlorobiaceae* specific (light green) with RAST annotations listed.

**Table S7. Differentially expressed *Chl. chlorochromatii* genes.** Transcriptome sequenced and normalized from axenic and symbiotic cultures during exponential and stationary phases (4 conditions) are compared, utilizing means of three biological replicates ranked by p-value, sorted by fold change if p<0.05. Contrasting [A] Axenic *Chl. chlorochromatii* in stationary vs. exponential phase, [B] Symbiotic *Chl. chlorochromatii* in stationary vs. exponential phase, [C] Differences of symbiotic vs. axenic *Chl. chlorochromatii* both in the exponential phase, or [D] Differences of symbiotic vs. axenic *Chl. chlorochromatii* both in the stationary phase, provides 16 conditions in which genes can be differentially expressed. Those significantly higher/lower in only one of the 4 comparisons are listed independently as tables A, B, C, or D. Those found in two or more conditions, but not differentially regulated in others, are listed at intersections, where [A] ⌒ [B] is AB. Genes not differentially expressed listed as N. Thus, sub-tables **(A)**, **(B)**, **(C)**, **(D)**, **(AB)**, **(AC)**, **(AD)**, **(BC)**, **(BD)**, **(CD)**, **(ABC)**, **(ABD), (ACD), (BCD), (ABCD)**, and **(N)** are provided as seen in Fig. S9. Current locus tags highlighted as *C. chlorochromatii* specific (dark green) or *Chlorobiaceae* specific (light green) with RAST annotations listed.

**Table S8. Analysis of genes of** ‘***Ca.* Symbiobacter mobilis**’ **and other *Comamonadaceae*. (A)** 411 single-copy core genes used for a MLSA tree to differentiate *Comamonadaceae*. **(B)** 777 ORFs shared among all members of *Comamonadaceae*. **(C)** 254 genes missing in the reduced genome of ‘*Ca.* S. mobilis’, present in all other *Comamonadaceae*. **(D)** List of 657 ORFs unique to central rod ‘*Ca.* S. mobilis’ but absent from all other known type strain genomes of the family. RAST annotations of the newly sequenced genome are provided by ascending locus-tag number.

**Table S9. List of annotated genes of ‘*Candidatus* Symbiobacter mobilis’ with potential implications for interaction covered by the present study. (A)** Genes grouped by physiological role provided with current locus tags and RAST annotations from newly sequenced and circularized genome. **(B)** Gene groups linked to symbiosis visualized with presence/absence of protein orthologs from neighboring species. Unique genes to ‘*Ca.* S. mobilis’ highlighted in light pink.

**Table S10. Differentially expressed ‘*Ca.* S. mobilis genes’.** Means of three biological replicates ranked by p-value, sorted by fold change if p<0.05, with current locus tags and RAST annotations listed. **(A)** Exponential vs. stationary phases compared for ‘*Ca.* S. mobilis’ in consortia. **(B)** Clustered differentially expressed genes compared between all samples as in heatmap (Fig. S2c). ORFs unique to ‘*Ca.* S. mobilis’ in pink. Genes of interest include ORFs upregulated when the *Chlorobium* symbiosis proteins are up-regulated (in sulfide-starved stationary phase; Table S7; Fig. S10b), labelled in purple.

**Table S11. Giant Symbiosis Protein LCMS data from SDS-PAGE and whole cell lysate Proteomics runs digested with trypsin. (A)** The top 10 identified proteins from each SDS-PAGE excised band are listed, confirming titin and mysosin-6 of mouse cardiac tissue (highlighted green), and low peptide hits for the giant symbiosis proteins (highlighted orange). **(B)** All identified enzymatically lysed peptide fragments of Cag_663 and Cag_665, from all runs and analysis.

## Legends of Extended Data

**Fig. E1. Comparative genomics of consortia genomes. (a)** Alignment of improved chromosome sequences generated in this study vs. existing sequences using a Progressive Mauve algorithm (similarity scored using match-seed-weight of 15 and 30,000 minimum compute Locally Collinear Blocks; as a plugin to *Geneious* 2023.0.2). The circularized genomes revealed several new motifs and improved locus tags. The *Chl. chlorochromatii* genome is 2,575,364 bp long (labeling is in Mbp) and encodes 2,096 CDS, including 3 *rrn* operons and 46 tRNAs. The ‘*Ca.* Symbiobacter mobilis’ genome is 3,045,420 bp long and has 2,469 CDS, including 6 *rrn* operons and 45 tRNAs. **(b)** Phylogenomic relationships of available genome sequences of the type strains of members of the phylum *Chlorobiota* (*left*) shown as MLSA phylogenetic tree using RAxML based on 222 single-copy core genes (Table S1A). The upset plot (*right*) depicts the differential gene content of *Chlorobiota*, including the 354 genes unique to *Chl. chlorochromatii* (green dot; Table S1C), and the 217 genes shared by all members of the family *Chlorobiaceae* (labeled in pink; Table S1B).

**Fig. E2. Predicted domains and operon structure of the three symbiosis genes, and operon structure of secretion system genes. (a)** ORFs of symbiosis genes contain numerous specific domains as identified via InterPro (102.0), as well as HK97-folds and C-terminus alginate lyase/ MtxA aerotaxis domains annotated with HHPred (PDB_mmCIF70_30_Mar). The specific protein regions used for generating primary antibodies for localization studies in this study are also indicated for Cag_663 and Cag_665. **(b)** Structure of the entire Cag_662/663/664/665 and Cag_2037/2038 operons. **(c)** Structure of the six T1SS operons present in the *Chl. chlorochromatii* genome which encode all three essential components of T1SSs (compare also Table S8). Genes unique to this genome are highlighted in green. Genes encoding inner membrane ATP-binding cassette *abc*, membrane fusion protein *mfp*, outer membrane factor *omf*, transcriptional regulator *xre*, unknown protein ORFs “*?”*, and zeta toxin “*ζ toxin”* are indicated.

**Fig. E3. Predicted domains, 3D-structures and potential interaction of Cag_2037 and Cag_2038 gene products. (a)** Specific domains as identified via InterPro (102.0). DUF, N-terminal domain of unknown function. **(b)** Predicted structures of Cag_2037 and Cag_2038 gene products and model of the interaction between the Cag_2037 cohesin and the Cag_2038 dockerin domains. Model building and predictive strength provided later in supplemental files.

**Fig. E4. T1SSs of *Chl. chlorochromatii*. (a)** *Chlorobiota* phylum phylogenetic relationship (*left*) utilizing available genomes of type strains for a MLSA phylogenetic tree with RAxML [**84**] based on 222 single-copy core genes, permitted the comparison of tallied number of annotated T1SS genes per strain (*tolC*, black squares; *hlyD*, grey squares; *hlyB*, white squares). **(b)** Phylogenetic analysis of T1SS components (*tolC*, *hlyB*, *hlyD*) found in an operon, are likely recently horizontally transferred, due to low sequence homology with all other *Chlorobiota* type strains. Nearest neighbors and distant relatives compiled via BlastN and ConSurf [Yariv *et al.,* 2023], aligned aa in MAFFT (v.7) [**91**] with conserved aa used for phylogenetic trees built using Neighbor-Joining method [**92**], JTT substitution model [**93**], and 1000 bootstrap replicates. Trees rooted at midpoint, displaying bootstrap values >10 at nodes. *Chlorobiota* highlighted in green. Outer membrane, OM; Inner membrane, IM; amino acid, aa.

**Fig. E5. Specific aspects of the structural domains predicted for giant symbiosis proteins.** The entire modelled structure of **(a)** Cag_663 and **(b)** Cag_665 both contain ∼12,000 to 14,000 aa-long repetitive parallel β-helix C- terminal regions that are hydrophilic, have low flexibility, multiple cation-binding domains, and high similarity to proteins of uncultured multicellular magnetotactic bacteria. While C-termini are similar, both proteins have unique N-terminal hydrophobic β-sheet structures (see panels c and d). **(c)** The Cag_663 N-terminus between aa positions 1 and 13,350 contains a long filamentous β-sheet with regular intervals of protruding BIg-like domains with HK97-folds known from phage capsid assembly. **(d)** The N-terminus of Cag_665 between aa positions 1 and 5,100 is predicted to contain a laminated β-sheet. The hydrophobic clefts are indicated by black arrows, cation-binding domains with red arrows. Numbers in colored circles indicate ∼1,500 aa fragments of Cag_663 or Cag_665 that were independently structurally predicted in AlphaFold3 prior to alignment in PyMOL to generate complete structural predictions of either protein (Fig. S1).

**Fig. E6. Culture conditions and growth kinetics of ‘*Chlorochromatium aggregatum’*. (a)** Batch cultures were grown in sealed 5L Duran flasks (Schott Mainz, Germany) to obtain the cell yields required for transcriptomic and proteomic analysis. Cultures were supplemented with neutralized sulfide solution (final concentration 0.5 mM) every second day over a period of 10 days. Afterwards, sulfide addition was stopped to induce sulfide starvation and transition cell metabolism into stationary phase. Decanting dense planktonic biomass exposed the scotophobically formed biofilm of pure consortia positioned nearest the light-source. **(b)** Spectra of whole cells of a consortia culture (bold) and of the pigment extract (thin line) indicating optimal location for optical density (OD) measurement of cultures at 600 nm and of BChl *c* extracts at 663nm. **(c)** Parallel biomass measurements included OD 600, protein, total dry weight, and quantification of BChl *c* (Absorbance A at 663 nm) of planktonic cells (blue), scotophobic biofilm (green), and calculated total biomass per 5L flask (grey). **(d)** Correlation of the different types of biomass measurements of phototrophic consortia.

**Fig. E7. Expression of *Chl. chlorochromatii* symbiosis factors associated within operons or connected among a transcriptome network. (a)** Delineation of Cag_663, Cag_665, Cag_2037, and neighboring gene transcripts with ORF coverage for exponential and stationary growth of axenic/symbiotic cultures. No. of transcripts were determined in triplicate based on transcriptome short reads mapped to genome, visualized in log10 scale up to 10,000 reads. Inserts depict the transcript frequencies of intergenic regions in an expanded view, with transcript ends denoted (arrows). **(b)** Network analysis of changes in *Chl. chlorochromatii* transcriptomes upon transition between exponential and stationary phases of growth and in either axenic or in consortia cultures constructed with BioLayout (3.4) utilizing a 2.2× inflation for the Markov clustering (MCL) algorithm. Calculations are based on the correlations between standardized transcript counts of the different genes under varied growth and symbiotic conditions. Each individual gene is represented by a node (sphere); edges (lines) indicate correlations above a 0.85 Pearson correlation threshold, with lengths of edges inversely proportional to the strength of correlation. Clusters below 3 nodes were filtered out. Individual genes associated with clusters 1 to 4 are listed in Tables S6B-S6E. **(c)** The three symbiosis genes Cag_663, Cag_665 and Cag_2037 are found in cluster 1, along with associated secretion system genes Cag_662, Cag_665, Cag_941, Cag_1676, and Cag_2038. Network graph in 3D animation can be viewed as a supplemental file.

**Fig. E8. Comparative genomics and transcriptomics of consortia central rod ‘*Ca.* S. mobilis’. (a)** Phylogenomic relationship of available genome sequences of symbiont to the type strains of related *Comomonadaceae* (*left*) shown as MLSA phylogenetic tree using RAxML based on 411 single-copy core genes (Table S8A). The upset plot (*middle*) depicts the differential gene content, including the 777 ORFS shared among all members (blue; Table S8B), 254 genes absent in ‘*Ca.* S. mobilis’ (green; Table S8C), and the 657 unique to ‘*Ca.* S. mobilis’ (maroon; Table S8D). Total genome size (right) shows the diminished length of symbiotic ‘*Ca.* S. mobilis’ (black arrow) as compared to all other free living *Comamonadaceae*. **(b)** Heatmap plot of significantly differentially expressed normalized values of ‘*Ca.* S. mobilis’genes between exponential (Expo.) and stationary (Stat.) phases of growth induced by H_2_S limitation (Fig. S7). Total differential genes (Table S10A) and clusters (Table S10B) provided. **(c)** Volcano plot of differentially expressed genes, highlighting those linked to symbiosis; motility, pili, chemotaxis and alginate synthesis related. Genes unique to ‘*Ca.* S. mobilis’, highlighted pink.

**Fig. E9. Controls of the super-resolution protein localization experiments of the three symbiosis proteins. (a)** Cag_2027, **(b)** Cag_663, and **(c)** Cag_665 protein specific fluorescence-based tests. *E. coli* XL-1 Blue cells expressing recombinant proteins Cag_2038, or fragments of Cag_663 or Cag_665 as antisera targets served as positive controls. Fluorescence of FM4-64FX membrane stain, 4’,6-diamidino-2-phenylindole (DAPI), and Alexa 488 nm immunofluorescence (IMF) signals of the expressed proteins are shown as well as the overlay of the three different fluorescence signals. In negative controls, pre-immune sera were applied in TIRF or dSTORM analyses of the phototrophic consortium ‘*C. aggregatum*’ or its isolated epibiont *Chl. chlorochromatii* CaD. In this case, bacteriochlorophyll *c* (BChl) autofluorescence was used instead of FM4-64FX to specifically localize epibionts. Arrows indicate the position of each central bacterium ‘*Ca.* S. mobilis’. Scale bar, 5 µm. **(d)** Immuno-gold labelling controls, shown as an average number of Cag_2037, Cag_663, or Cag_665 protein specific immunogold-particles (IMG) per cell compartments cytoplasm (Cyt.) or envelope (Env.) of ‘*C. aggregatum*’ with or without pre-immune sera. Vertical bars depict standard error. Significant differences were determined by unpaired t-tests (*, p<0.05; **, p<0.01; ***, p<0.001).

## Legends of Supplementary Figures

**Fig. S1. Modeling the predicted structure of giant symbiosis proteins.** Full protein structural predictions of **(a)** Cag_663 and **(b)** Cag_665 using AlphaFold3 to determine 1500 aa fragments with 300 aa overlap which could then be assembled in PyMOL to construct the final assembly. **(c)** Inputs of 5000 aa segments, such as Cag_663:27001-32000 yielded tertiary bundling or hairball structures with low confidence values, indicating artifacts. For final assembly amino acids 14712, 19443, and 19960 of Cag_663, as well as amino acids 5990 and 10614 of Cag_665 had to be manually rotated to prevent erroneous folding back onto the predicted structure. Fragments used to construct each protein are displayed on following pages, as Cag_663, Cag_664, and Cag_665, followed by Cag_2037 with Cag_2038.

**Fig. S2. Alginate lyase supporting information. (a)** Cag_2037 structural predictions were strengthened by addition of five calcium cations, **(b)** which imbed into the RTX domain. **(c)** Partial predicted structures elongated BIg domains, with **(d)** concatenated complete Cag_2037 structure revealing a stable, hydrophilic structure that has a flexible N- terminal cohesion domain and an electrostatic C- terminal RTX domain. **(e)** Precise amounts of alginate were hydrolyzed and analyzed via MS to achieve a standard curve of detectable levels. **(F)** Exogenous alginate lyase diminishes ‘*C. aggregatum*’ biofilm production, **(G)** as seen from the side the reduction of phototactic ability is a result of disintegrated consortia. **(H)** Alginate synthesis genes clustered within the genome of *Gammaproteobacteria* representative *Azotobacter vinlandii,* while a new gene cluster arrangement was detected for *Betaproteobacteria* ‘*Ca.* S. mobilis’, and close relatives within *Rhodoferax*.

**Fig. S3. Modeling of protein-protein interaction for the β-plate region of Cag_663. (a)** The largest section possible (2,200 aa) of the twisted β-sheet with hydrophobic cleft and one extended filamentous hemagglutinin arm encoded by Cag_663 was modelled as a monomer (upper part) or dimer (lower part). Two β-sheets can dimerize at their hydrophobic clefts, where each co-occurring capsid-like/HK97-fold motifs remain exposed to the outside. **(b)** A dimeric region which excludes the extended capsid-like/HK97-fold motifs also hides the hydrophobic cleft, exposing only hydrophilic regions to the outside.

**Fig. S4. Differentially expressed *Chl. chlorochromatii* genes.** Transcriptome sequenced and normalized from axenic and symbiotic cultures during exponential and stationary phases (4 conditions) are compared, utilizing means of three biological replicates ranked by p-value, sorted by fold change if p<0.05. Contrasting [A] Axenic *Chl. chlorochromatii* in stationary vs. exponential phase, [B] Symbiotic *Chl. chlorochromatii* in stationary vs. exponential phase, [C] Differences of symbiotic vs. axenic *Chl. chlorochromatii* both in the exponential phase, or [D] Differences of symbiotic vs. axenic *Chl. chlorochromatii* both in the stationary phase, provided 16 conditions in which genes can be differentially expressed. **(a)** Total genes differentially expressed in all conditions for each intersection are listed (without differentiating if up or down regulated), where specific genes of each group are provided in the corresponding supplemental Table S7. **(b)** *Chl. chlorochromatii* genes that are specific to the *Chlorobiaceae* family in brackets, or unique to this species that are differentially expressed in all conditions for each intersection are listed (without differentiating if up or down regulated). Cag_663, Cag_665 and Cag_2037 are in Groups ABCD, ABCD, and AB, respectively. *While *Chl. chlorochromatii* has 2,096 protein coding genes, all 2,152 ORF’s are included in this analysis. **(c)** Significant differences of select normalized counts of symbiosis genes. Axenic, Axn.; Consortia, Con.; Exponential, Expo.; Stationary, Stat.

**Fig. S5. Detailed analysis of differential expression patterns in *Chl. chlorochromatii* across different growth conditions and symbiotic states.** Volcano plots of pairwise comparisons between exponential and stationary phase for the same **(a)** axenic or **(b)** symbiotic state (upper panels) are contrasted to expression patterns found between axenic or symbiotic state in the same **(c)** exponential or **(d)** stationary growth phase (lower panels). The three symbiosis genes, associated genes, and secretion systems are colored as seen in Figure 2,3,S8 and elsewhere. Genes highlighted green are unique to *Chl. chlorochromatii* among *Chlorobiota*.

**Fig S6. Trypsin enzymatic digestion of titin as compared to giant virulence-like symbiosis factors.** *In silico* peptides degraded by Trypsin are generated in Expasy [**77**], allowing 0 missing cleavages, and filtering peptides >500 Da, showing those below certain thresholds. Masses of peptides calculated as monoisotopic [M+H]+. Detectable peptides then mapped back to entire protein sequence N- to C-termini in ggplot (R). https://web.expasy.org/peptide_mass/. Below each *in silico* digest is pooled *in vitro* digest results of *Mus* or *Chlorobium* biomass, revealing similar density of cut sites for all 3. While titin is degradable by trypsin, this suggests both symbiosis proteins are indeed present, but are resistant to complete trypsin digest. Amino acid position, aa.

**Fig. S7. Chemotaxis array hypothesis of Cag_663 active domain. (a)** GA fixed ‘*C. aggregatum*’ consortia reveal ‘*Ca.* S. mobilis’ <u>c</u>entral <u>b</u>acterium to contain para<u>c</u>rystalline (CBC) structures (arrow) as previously documented [**15**], which are similar in appearance to (Aero)chemotaxis arrays [**73**]. Scale bar, 0.4µm. **(b)** As in Fig. S2, regions of Cag_663 hydrophobicity will form dimers, where **(c)** flexible BIg domain arms have the possibility to become extended. **(d)** Looking at density or paracrystalline structures, auto-correlation of CBC reveals oblique orientation of the subunits, in which (e) extended and stacked Cag_663 active domains potentially fit.

