## Extended and Supplemental Figures for "Interspecies transfer of giant virulence-factor-like proteins in a bacterial symbiosis"

Research Article
Biological Sciences – *Microbiology*

### Extended data for:

### Interspecies transfer of giant virulence-factor-like proteins in a bacterial symbiosis

Steven B. Kuzyk[a†], Petra Henke[a†], Anika Methner[a], Tina Rietschel[b], Franziska Burkart[a], Mathias Müsken[c], Meina Neumann-Schaal[d], Gerhard Wanner[e], and Jörg Overmann[a,f,g*]

[a] Department of Microbial Ecology and Diversity Research, Leibniz-Institute DSMZ - German Collection of Microorganisms and Cell Cultures, Braunschweig, Germany

[b] Cellular Proteome Research (CPRO), HZI - Helmholtz Centre for Infection Research, Braunschweig, Germany

[c] Central Facility for Microscopy, HZI - Helmholtz Centre for Infection Research, Braunschweig, Germany

[d] Department of Metabolomics & Services, Leibniz-Institute DSMZ - German Collection of Microorganisms and Cell Cultures, Braunschweig, Germany

[e] Plant Sciences, Faculty of Biology, Ludwig-Maximilians-University Munich, Germany

[f] Bavarian State Collections of Natural History, Munich, Germany

[g] Chair for Molecular Diversity Research, Faculty of Biology Ludwig-Maximilians-University Munich, Munich, Germany

† These authors contributed equally to the work

**Running title:** Interspecies transfer of giant symbiosis proteins

**Keywords:** Microbial symbiosis, RTX-toxin, RTX adhesion, Virulence factors, Microbial consortia

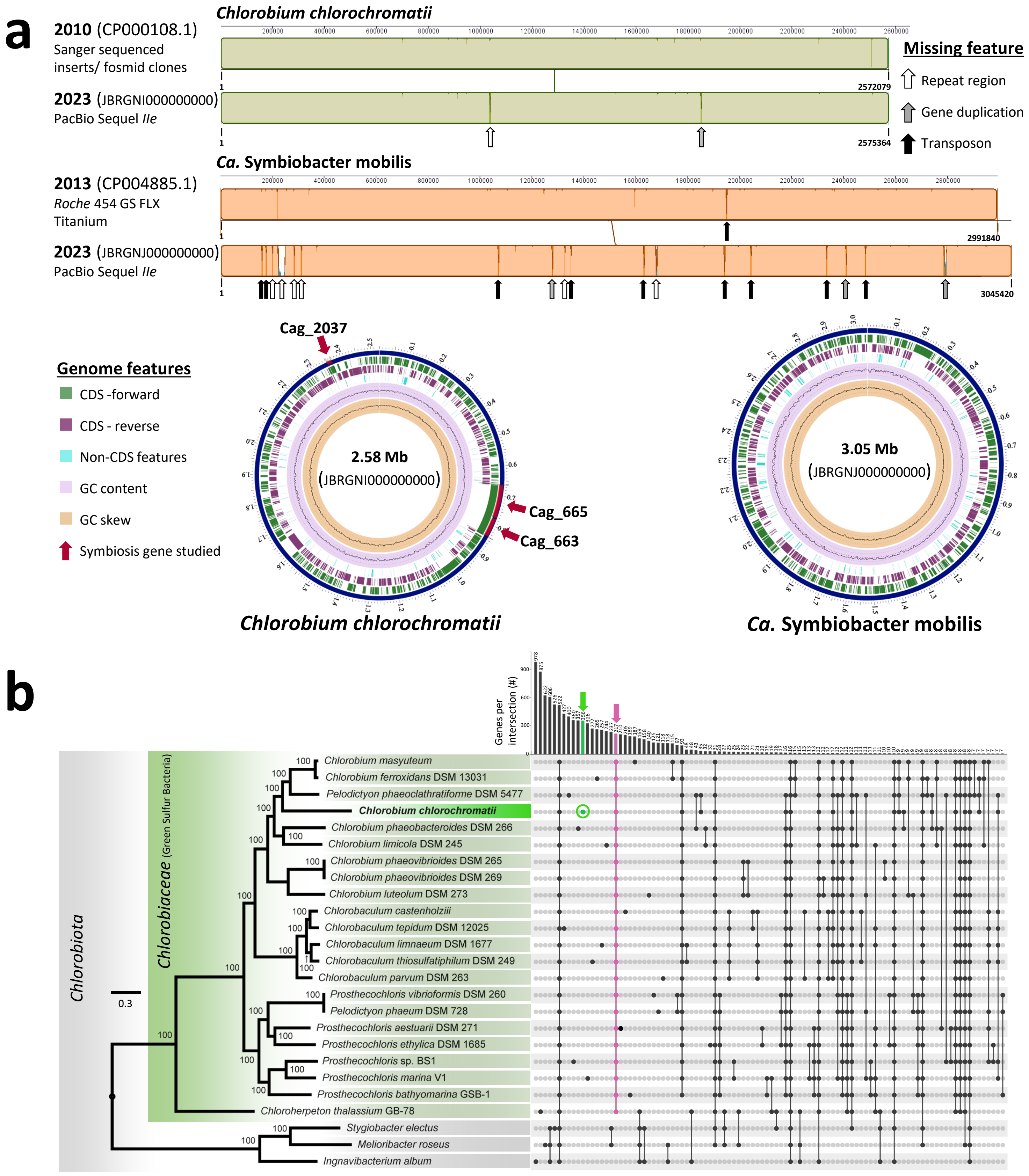


**Fig. E1. Comparative genomics of consortia genomes. (a)** Alignment of improved chromosome sequences generated in this study vs. existing sequences using a Progressive Mauve algorithm (similarity scored using match-seed-weight of 15 and 30,000 minimum compute Locally Collinear Blocks; as a plugin to *Geneious* 2023.0.2). The circularized genomes revealed several new motifs and improved locus tags. The *Chl. chlorochromatii* genome is 2,575,364 bp long (labeling is in Mbp) and encodes 2,096 CDS, including 3 *rrn* operons and 46 tRNAs. The `*Ca.* Symbiobacter mobilis´ genome is 3,045,420 bp long and has 2,469 CDS, including 6 *rrn* operons and 45 tRNAs. **(b)** Phylogenomic relationships of available genome sequences of the type strains of members of the phylum *Chlorobiota* (*left*) shown as MLSA phylogenetic tree using RAxML based on 222 single-copy core genes (Table S1A). The upset plot (*right*) depicts the differential gene content of *Chlorobiota*, including the 354 genes unique to *Chl. chlorochromatii* (green dot; Table S1C), and the 217 genes shared by all members of the family *Chlorobiaceae* (labeled in pink; Table S1B).

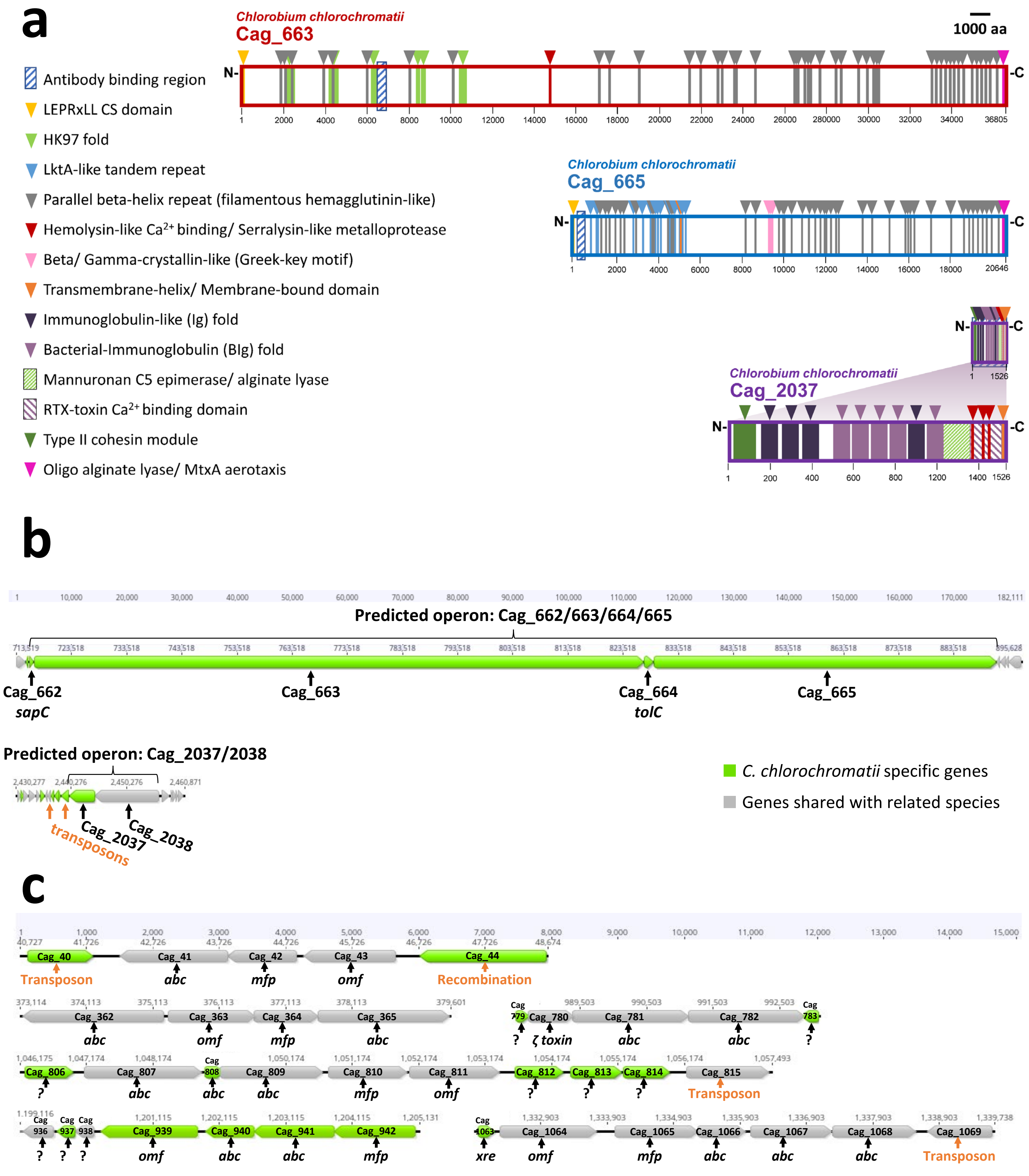


**Fig. E2. Predicted domains and operon structure of the three symbiosis genes, and operon structure of secretion system genes. (a)** ORFs of symbiosis genes contain numerous specific domains as identified via InterPro (102.0), as well as HK97-folds and C-terminus alginate lyase/ MtxA aerotaxis domains annotated with HHPred (PDB_mmCIF70_30_Mar). The specific protein regions used for generating primary antibodies for localization studies in this study are also indicated for Cag_663 and Cag_665. **(b)** Structure of the entire Cag_662/663/664/665 and Cag_2037/2038 operons. **(c)** Structure of the six T1SS operons present in the *Chl. chlorochromatii* genome which encode all three essential components of T1SSs (compare also Table S8). Genes unique to this genome are highlighted in green. Genes encoding inner membrane ATP-binding cassette *abc*, membrane fusion protein *mfp*, outer membrane factor *omf*, transcriptional regulator *xre*, unknown protein ORFs "*?*", and zeta toxin "*ζ toxin*" are indicated.

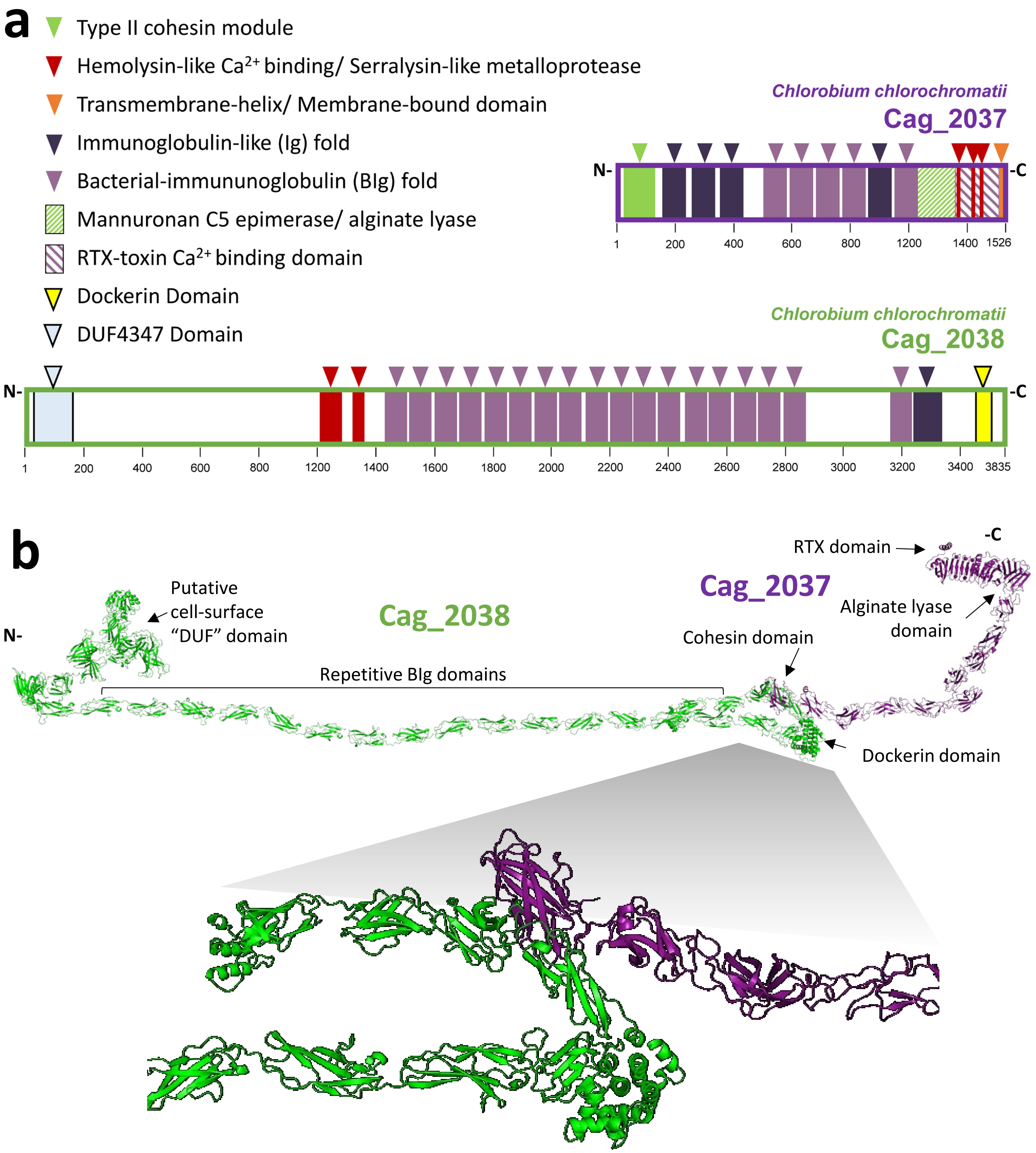


**Fig. E3. Predicted domains, 3D-structures and potential interaction of cellulosome-associated Cag_2037 and Cag_2038 gene products. (a)** Specific domains as identified via InterPro (102.0). DUF, N-terminal domain of unknown function. **(b)** Predicted structures of Cag_2037 and Cag_2038 gene products and model of the interaction between the Cag_2037 cohesin and the Cag_2038 dockerin domains. Model building and predictive strength provided later in supplemental files.

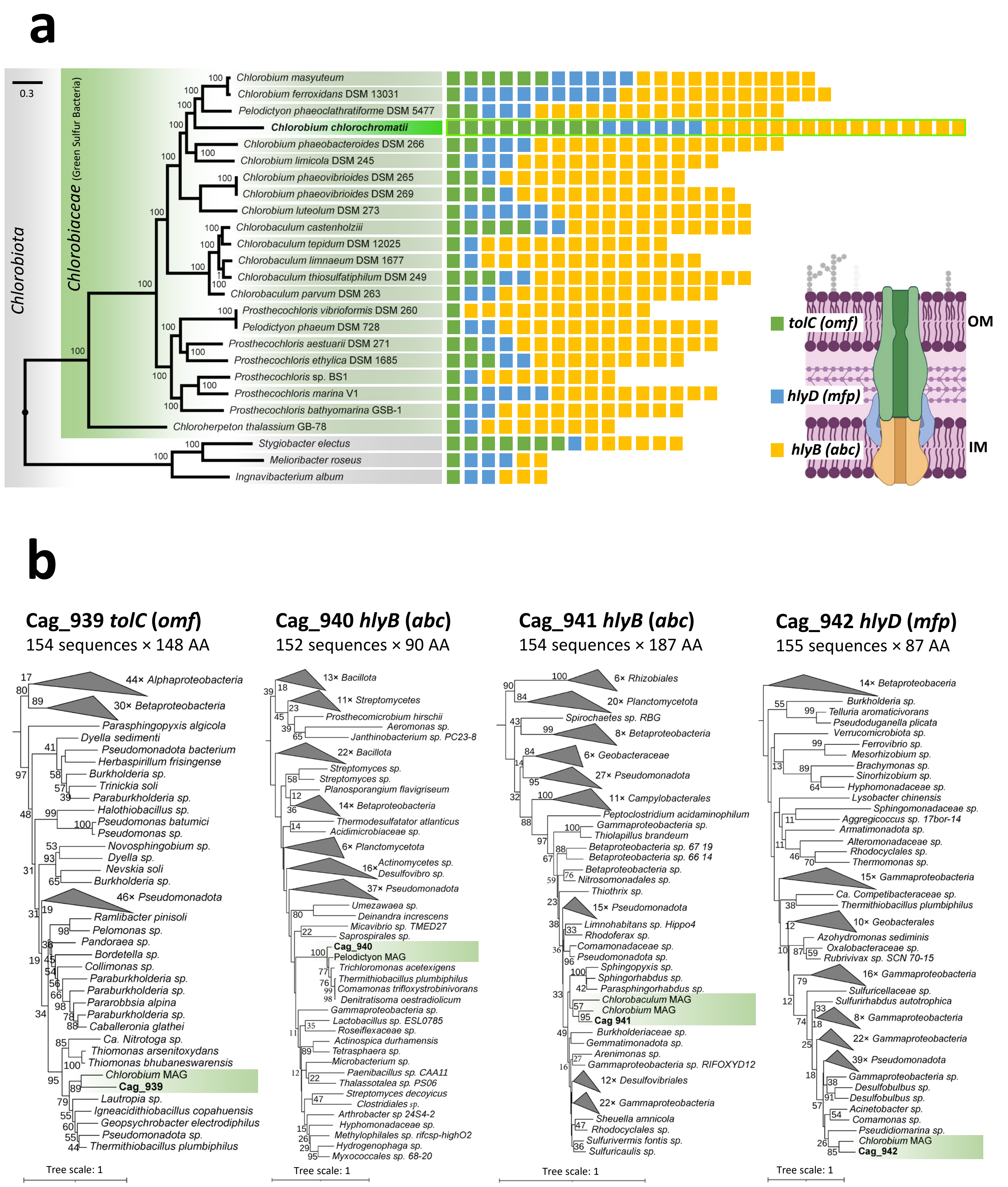


**Fig. E4. T1SSs of *Chl. chlorochromatii*. (a)** *Chlorobiota* phylum phylogenetic relationship (*left*) utilizing available genomes of type strains for a MLSA phylogenetic tree with RAxML [84] based on 222 single-copy core genes, permitted the comparison of tallied number of annotated T1SS genes per strain (*tolC*, black squares; *hlyD*, grey squares; *hlyB*, white squares). **(b)** Phylogenetic analysis of T1SS components (*tolC*, *hlyB*, *hlyD*) found in an operon, are likely recently horizontally transferred, due to low sequence homology with all other *Chlorobiota* type strains. Nearest neighbors and distant relatives compiled via BlastN and ConSurf [Yariv *et al.*, 2023], aligned aa in MAFFT (v.7) [93] with conserved aa used for phylogenetic trees built using Neighbor-Joining method [94], JTT substitution model [Jones *et al.*, 1992], and 1000 bootstrap replicates. Trees rooted at midpoint, displaying bootstrap values >10 at nodes. *Chlorobiota* highlighted in green. Outer membrane, OM; Inner membrane, IM; amino acid, aa.

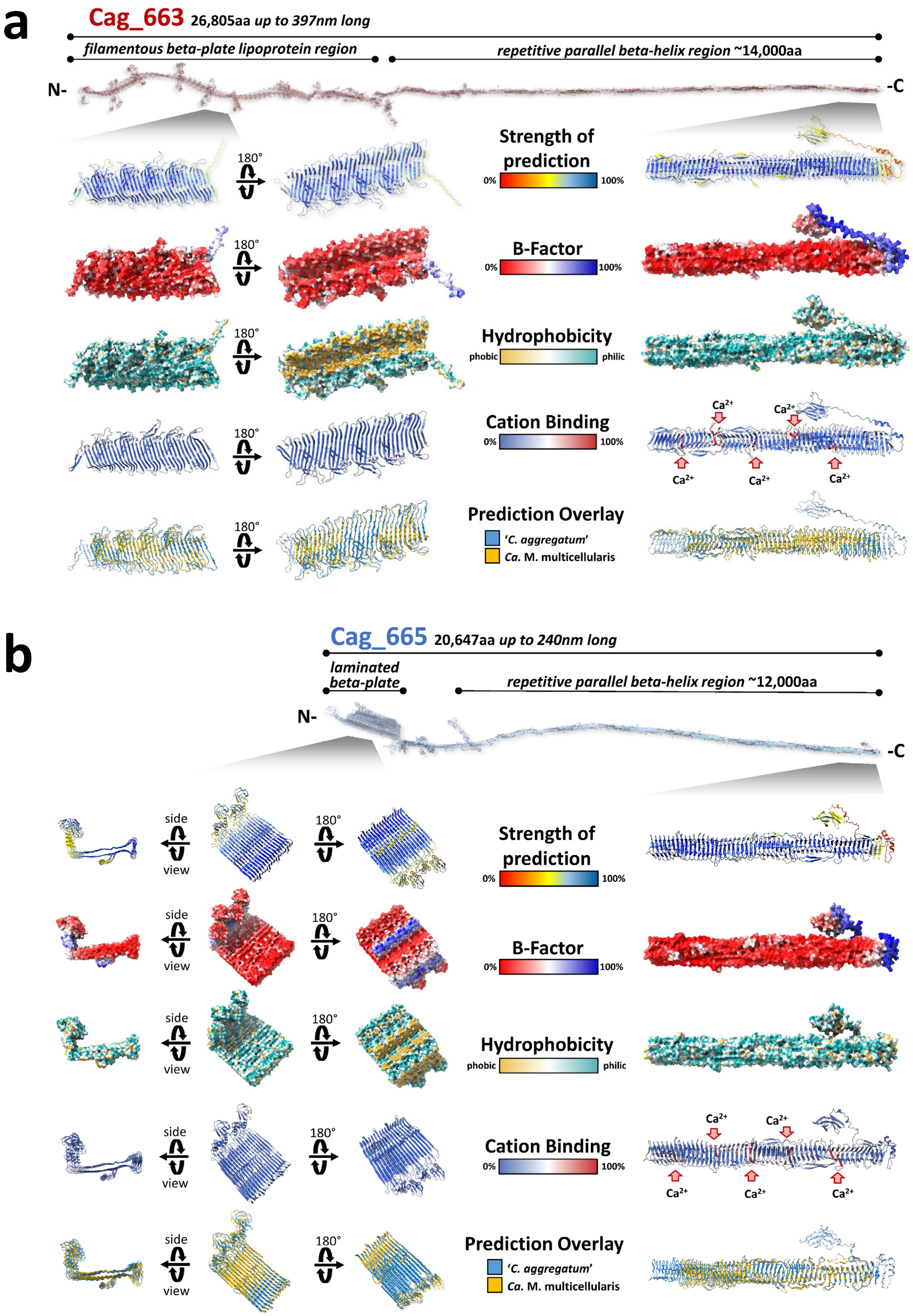


**Fig. E5.** (Continued on next page)

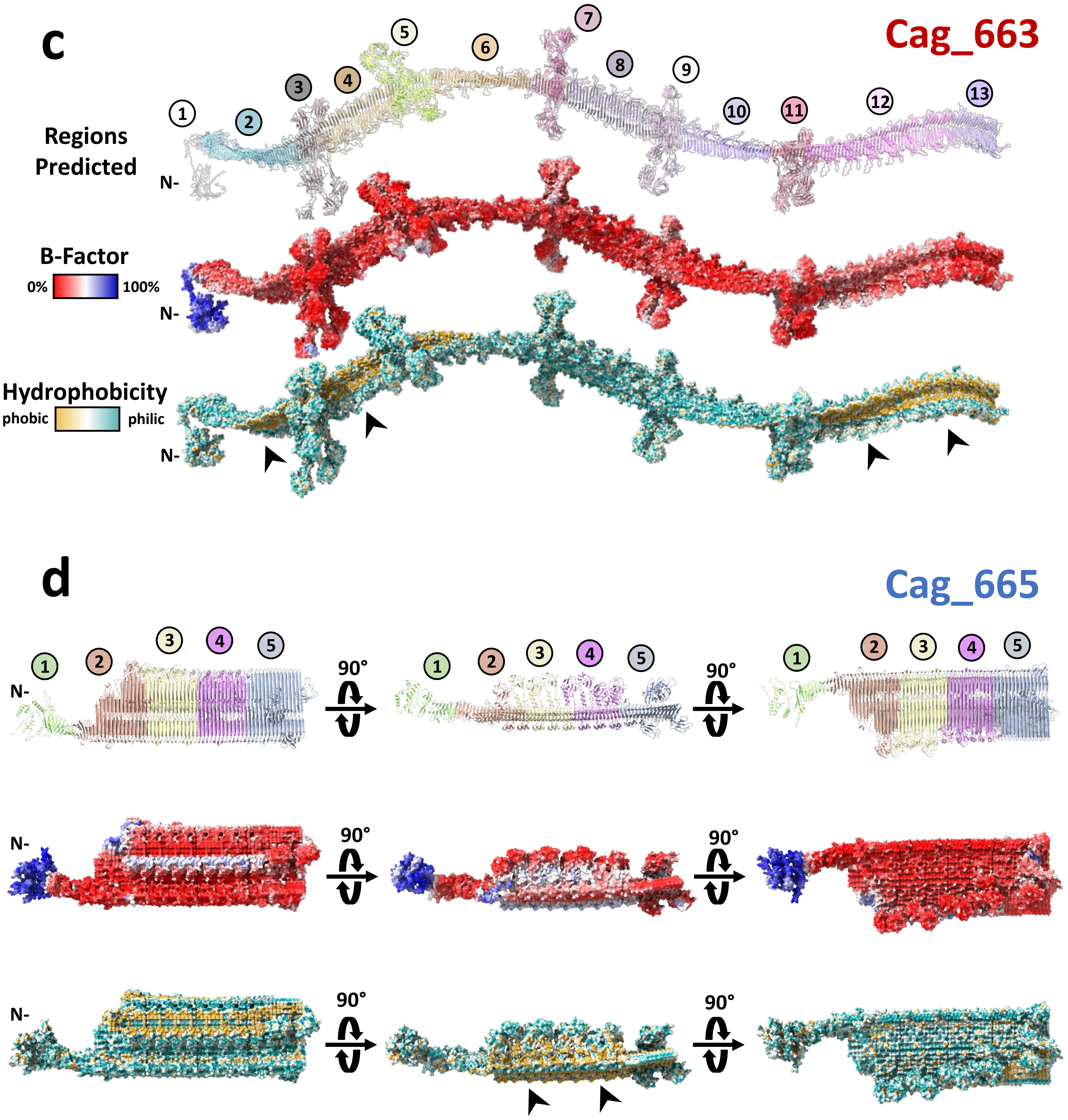


**Fig. E5. Specific aspects of the structural domains predicted for giant symbiosis proteins.** The entire modelled structure of **(a)** Cag_663 and **(b)** Cag_665 both contain ~12,000 to 14,000 aa-long repetitive parallel β-helix C- terminal regions that are hydrophilic, have low flexibility, multiple cation-binding domains, and high similarity to proteins of uncultured multicellular magnetotactic bacteria. While C- termini are similar, both proteins have unique N-terminal hydrophobic β-sheet structures (see panels c and d). **(c)** The Cag_663 N-terminus between aa positions 1 and 13,350 contains a long filamentous β-sheet with regular intervals of protruding BIg-like domains with HK97-folds known from phage capsid assembly. **(d)** The N-terminus of Cag_665 between aa positions 1 and 5,100 is predicted to contain a laminated β-sheet. The hydrophobic clefts are indicated by black arrows, cation-binding domains with red arrows. Numbers in colored circles indicate ~1,500 aa fragments of Cag_663 or Cag_665 that were independently structurally predicted in AlphaFold3 prior to alignment in PyMOL to generate complete structural predictions of either protein (Fig. S1).

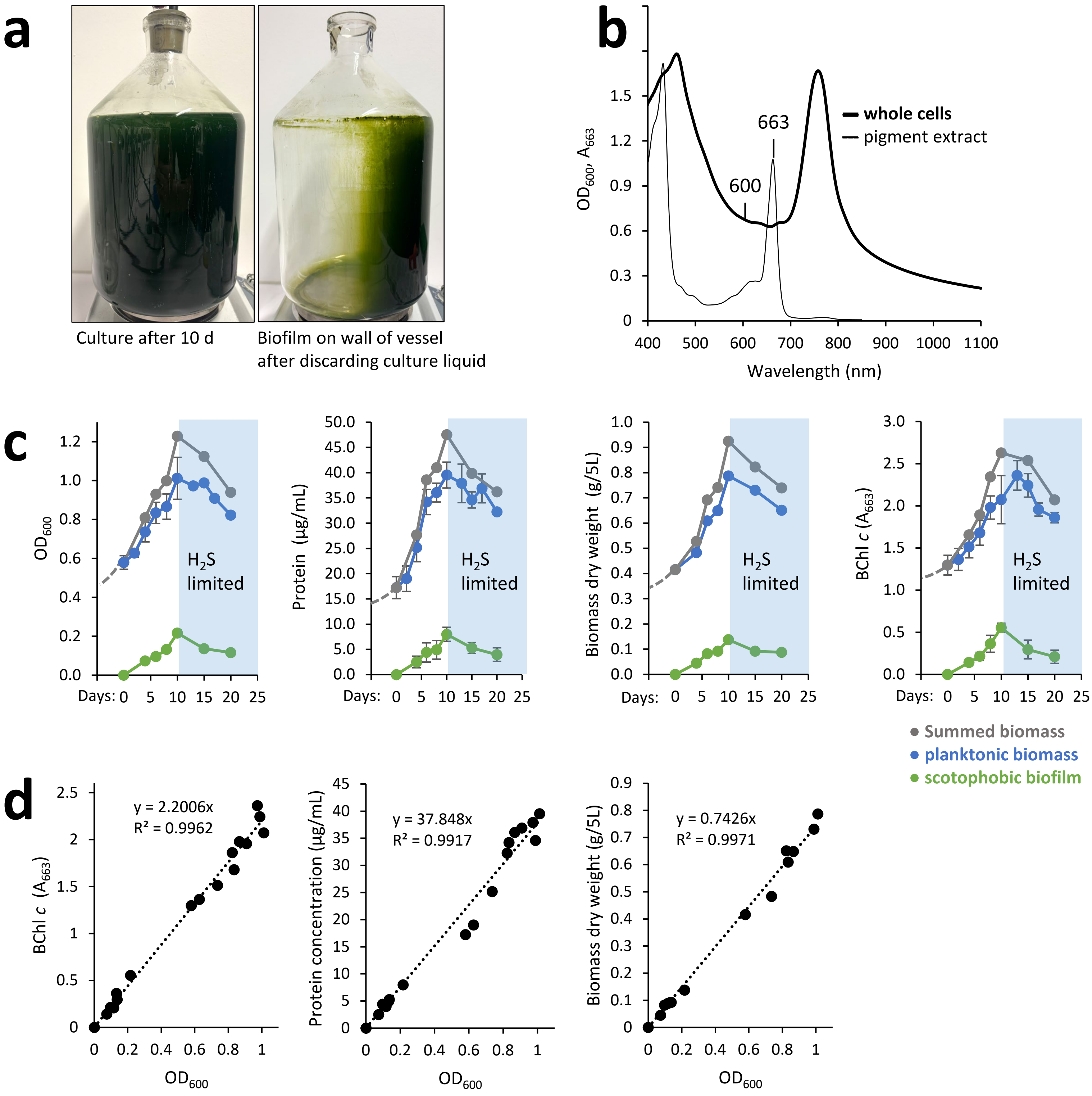


**Fig. E6. Culture conditions and growth kinetics of '*Chlorochromatium aggregatum*'. (a)** Cultures were grown in sealed static 5L Duran flasks (Schott Mainz, Germany) to obtain the necessary conditions required for transcriptomic and proteomic analysis. Cultures were supplemented with neutralized sulfide solution (final concentration 0.5 mM) every second day over a period of 10 days. Afterwards, sulfide addition was stopped to induce sulfide starvation and transition cell metabolism into stationary phase. Decanting dense planktonic biomass exposed the scotophobically formed biofilm of pure consortia positioned nearest the light-source. **(b)** Spectra of whole cells of a consortia culture (bold) and of the pigment extract (thin line) indicating optimal location for optical density (OD) measurement of cultures at 600 nm and of BChl *c* extracts at 663nm. **(c)** Parallel biomass measurements included OD 600, protein, total dry weight, and quantification of BChl *c* (Absorbance A at 663 nm) of planktonic cells (blue), scotophobic biofilm (green), and calculated total biomass per 5L flask (grey). **(d)** Correlation of the different types of biomass measurements of phototrophic consortia.

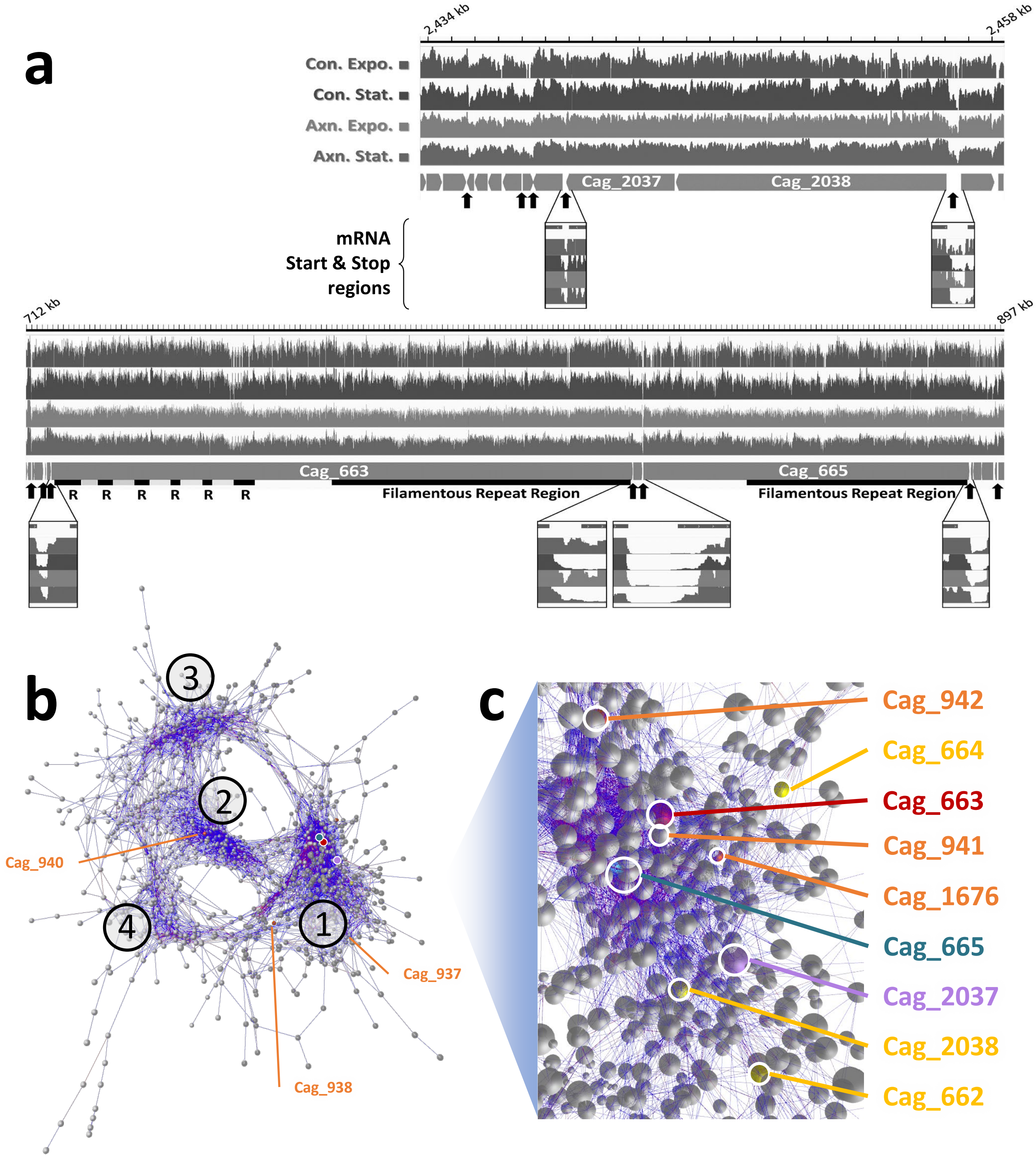


**Fig. E7. Expression of *Chl. chlorochromatii* symbiosis factors associated within operons or connected among a transcriptome network. (a)** Delineation of Cag_663, Cag_665, Cag_2037, and neighboring gene transcripts with ORF coverage for exponential and stationary growth of axenic/symbiotic cultures. No. of transcripts were determined in triplicate based on transcriptome short reads mapped to genome, visualized in log10 scale up to 10,000 reads. Inserts depict the transcript frequencies of intergenic regions in an expanded view, with transcript ends denoted (arrows). **(b)** Network analysis of changes in *Chl. chlorochromatii* transcriptomes upon transition between exponential and stationary phases of growth and in either axenic or in consortia cultures constructed with BioLayout (3.4) utilizing a 2.2× inflation for the Markov clustering (MCL) algorithm. Calculations are based on the correlations between standardized transcript counts of the different genes under varied growth and symbiotic conditions. Each individual gene is represented by a node (sphere); edges (lines) indicate correlations above a 0.85 Pearson correlation threshold, with lengths of edges inversely proportional to the strength of correlation. Clusters below 3 nodes were filtered out. Individual genes associated with clusters 1 to 4 are listed in Tables S6B-S6E. **(c)** The three symbiosis genes Cag_663, Cag_665 and Cag_2037 are found in cluster 1, along with associated secretion system genes Cag_662, Cag_665, Cag_941, Cag_1676, and Cag_2038. Network graph in 3D animation can be viewed as a supplemental file.

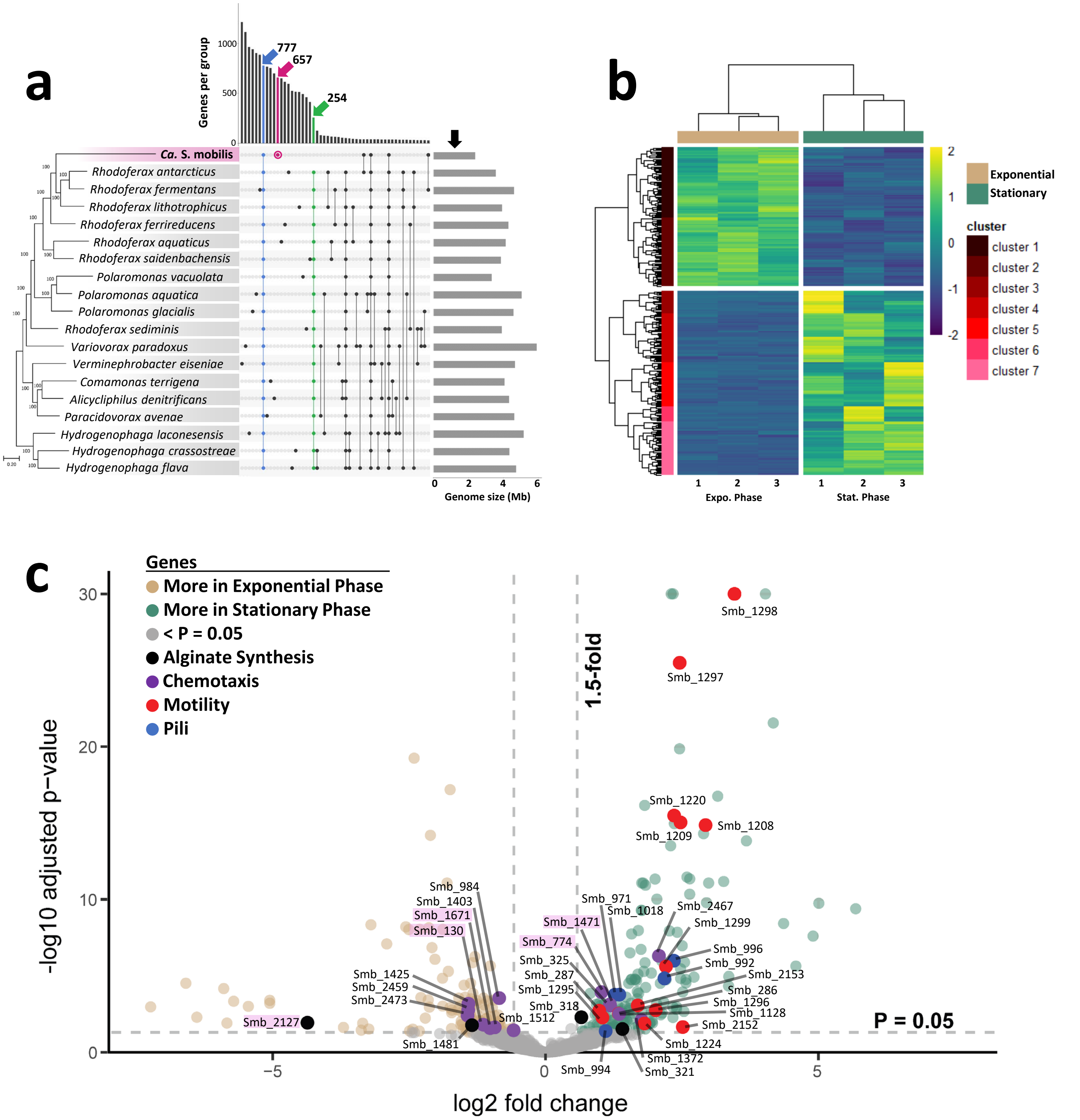


**Fig. E8. Comparative genomics and transcriptomics of consortia central rod \`*Ca.* S. mobilis´. (a)** Phylogenomic relationship of available genome sequences of symbiont to the type strains of related *Comomonadaceae* (*left*) shown as MLSA phylogenetic tree using RAxML based on 411 single-copy core genes (Table S8A). The upset plot (*middle*) depicts the differential gene content, including the 777 ORFS shared among all members (blue; Table S8B), 254 genes absent in \`*Ca.* S. mobilis´ (green; Table S8C), and the 657 unique to \`*Ca.* S. mobilis´ (maroon; Table S8D). Total genome size (right) shows the diminished length of symbiotic \`*Ca.* S. mobilis´ (black arrow) as compared to all other free living *Comamonadaceae*. **(b)** Heatmap plot of significantly differentially expressed normalized values of \`*Ca.* S. mobilis´genes between exponential (Expo.) and stationary (Stat.) phases of growth induced by $H_2S$ limitation (Fig. S7). Total differential genes (Table S10A) and clusters (Table S10B) provided. **(c)** Volcano plot of differentially expressed genes, highlighting those linked to symbiosis; motility, pili, chemotaxis and alginate synthesis related. Genes unique to \`*Ca.* S. mobilis´, highlighted pink.

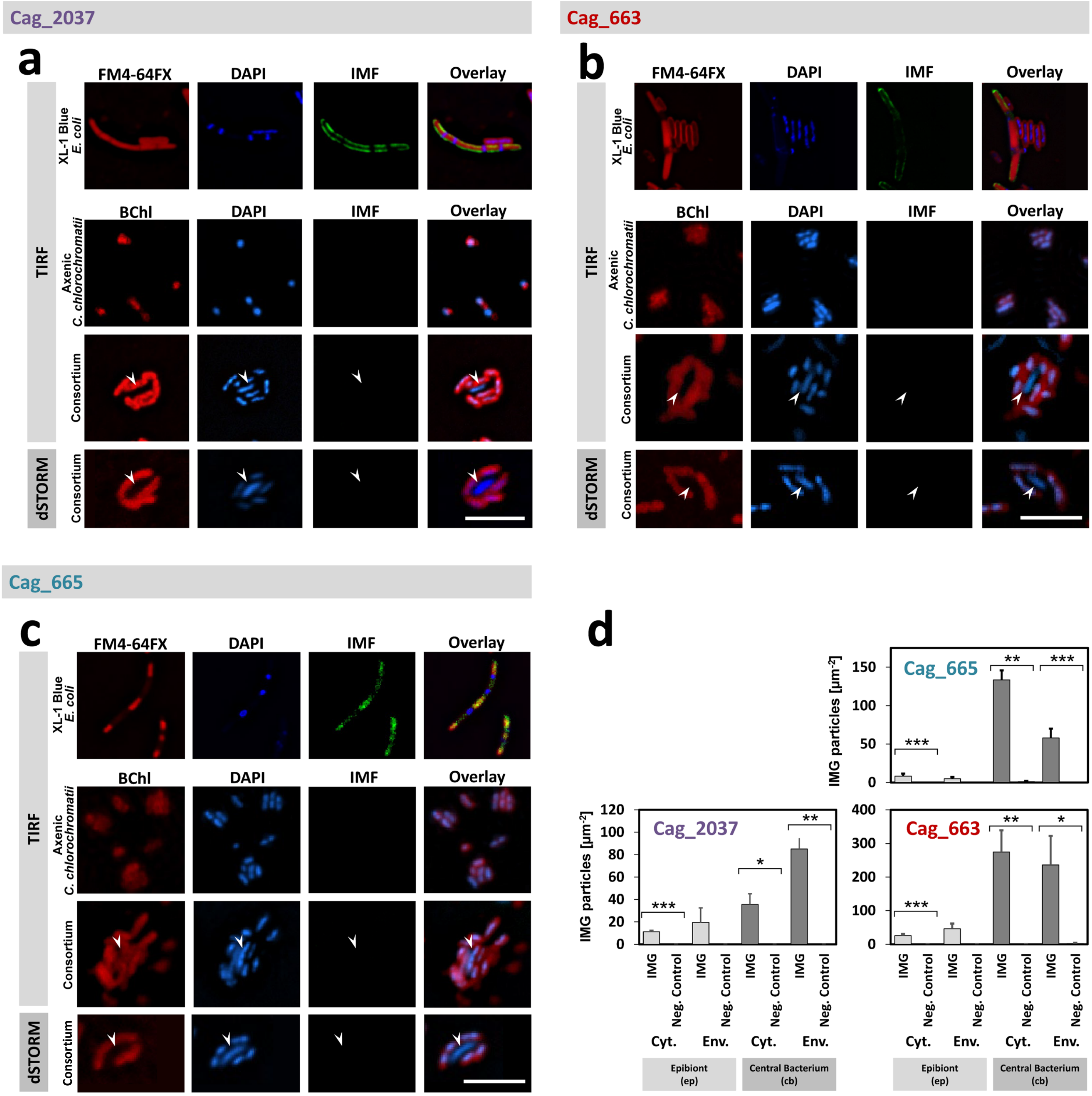


**Fig. E9. Controls of the super-resolution protein localization experiments of the three symbiosis proteins. (a)** Cag_2027, **(b)** Cag_663, and **(c)** Cag_665 protein specific fluorescence-based tests. *E. coli* XL-1 Blue cells expressing recombinant proteins Cag_2037, or fragments of Cag_663 or Cag_665 as antisera targets served as positive controls. Fluorescence of FM4-64FX membrane stain, 4',6-diamidino-2-phenylindole (DAPI), and Alexa 488 nm immunofluorescence (IMF) signals of the expressed proteins are shown as well as the overlay of the three different fluorescence signals. In negative controls, pre-immune sera were applied in TIRF or dSTORM analyses of the phototrophic consortium `*C. aggregatum*´ or its isolated epibiont *Chl. chlorochromatii* CaD. In this case, bacteriochlorophyll *c* (BChl) autofluorescence was used instead of FM4-64FX to specifically localize epibionts. Arrows indicate the position of each central bacterium `*Ca.* S. mobilis´. Scale bar, 5 µm. **(d)** Immuno-gold labelling controls, shown as an average number of Cag_2037, Cag_663, or Cag_665 protein specific immunogold-particles (IMG) per cell compartments cytoplasm (Cyt.) or envelope (Env.) of `*C. aggregatum*´ with or without pre-immune sera. Vertical bars depict standard error. Significant differences were determined by unpaired t-tests (*, p<0.05; **, p<0.01; ***, p<0.001).

Research Article
Biological Sciences – *Microbiology*

### Supplemental Information for:

### Interspecies transfer of giant virulence-factor-like proteins in a bacterial symbiosis

Steven B. Kuzyk[a†], Petra Henke[a†], Anika Methner[a], Tina Rietschel[b], Franziska Burkart[a], Mathias Müsken[c], Meina Neumann-Schaal[d], Gerhard Wanner[e], and Jörg Overmann[a,f,g*]

[a] Department of Microbial Ecology and Diversity Research, Leibniz-Institute DSMZ - German Collection of Microorganisms and Cell Cultures, Braunschweig, Germany

[b] Cellular Proteome Research (CPRO), HZI - Helmholtz Centre for Infection Research, Braunschweig, Germany

[c] Central Facility for Microscopy, HZI - Helmholtz Centre for Infection Research, Braunschweig, Germany

[d] Department of Metabolomics & Services, Leibniz-Institute DSMZ - German Collection of Microorganisms and Cell Cultures, Braunschweig, Germany

[e] Plant Sciences, Faculty of Biology, Ludwig-Maximilians-University Munich, Germany

[f] Bavarian State Collections of Natural History, Munich, Germany

[g] Chair for Molecular Diversity Research, Faculty of Biology Ludwig-Maximilians-University Munich, Munich, Germany

† These authors contributed equally to the work

**Running title:** Interspecies transfer of giant symbiosis proteins

**Keywords:** Microbial symbiosis, RTX-toxin, RTX adhesion, Virulence factors, Microbial consortia

**a**

| | Fragments start | Fragments end | length (aa) | overlap (aa) | predicted template modeling score pTM |
|---|---|---|---|---|---|
| 1 | 1 | 1500 | 1500 | - | 0.57 |
| 2 | 300 | 1800 | 1501 | 1200 | 0.64 |
| 3 | 1500 | 3000 | 1501 | 300 | 0.44 |
| 4 | 2700 | 3700 | 1001 | 300 | 0.78 |
| 5 | 3500 | 5000 | 1501 | 200 | 0.41 |
| 6 | 4900 | 6000 | 1101 | 100 | 0.80 |
| 7 | 5800 | 7300 | 1501 | 200 | 0.42 |
| 8 | 7100 | 8000 | 901 | 200 | 0.84 |
| 9 | 7900 | 9400 | 1501 | 100 | 0.43 |
| 10 | 9200 | 10100 | 901 | 200 | 0.84 |
| 11 | 9990 | 11490 | 1501 | 110 | 0.42 |
| 12 | 11290 | 12800 | 1511 | 200 | 0.61 |
| 13 | 12600 | 13350 | 751 | 200 | 0.87 |
| 14 | 13000 | 14800 | 1801 | 350 | 0.63 |
| 15 | 14500 | 16800 | 2301 | 300 | 0.60 |
| 16 | 16500 | 17900 | 1401 | 300 | 0.89 |
| 17 | 17307 | 18807 | 1501 | 593 | 0.55 |
| 18 | 18707 | 19957 | 1251 | 100 | 0.81 |
| 19 | 19680 | 21600 | 1921 | 277 | 0.47 |
| 20 | 21430 | 22430 | 1001 | 170 | 0.79 |
| 21 | 22107 | 23607 | 1501 | 323 | 0.85 |
| 22 | 23307 | 24807 | 1501 | 300 | 0.80 |
| 23 | 24507 | 26007 | 1501 | 300 | 0.78 |
| 24 | 25707 | 27207 | 1501 | 300 | 0.86 |
| 25 | 26907 | 28407 | 1501 | 300 | 0.87 |
| 26 | 28107 | 29607 | 1501 | 300 | 0.87 |
| 27 | 29307 | 30807 | 1501 | 300 | 0.85 |
| 28 | 30507 | 32007 | 1501 | 300 | 0.87 |
| 29 | 31707 | 33207 | 1501 | 300 | 0.85 |
| 30 | 32907 | 34407 | 1501 | 300 | 0.86 |
| 31 | 34107 | 35607 | 1501 | 300 | 0.85 |
| 32 | 35307 | 36806 | 1500 | 300 | 0.79 |

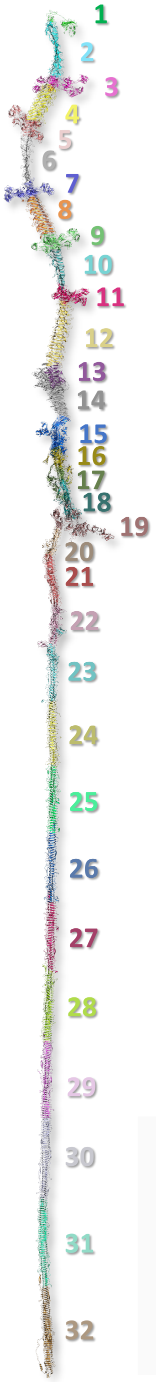


**b**

| | Fragments start | Fragments end | length (aa) | overlap (aa) | predicted template modeling score pTM |
|---|---|---|---|---|---|
| 1 | 1 | 500 | 500 | - | 0.57 |
| 2 | 250 | 1500 | 1250 | 250 | 0.83 |
| 3 | 1200 | 2700 | 1500 | 300 | 0.84 |
| 4 | 2400 | 3900 | 1500 | 300 | 0.89 |
| 5 | 3600 | 5100 | 1500 | 300 | 0.81 |
| 6 | 5000 | 7500 | 2500 | 100 | 0.78 |
| 7 | 7400 | 8900 | 1500 | 100 | 0.87 |
| 8 | 8700 | 10200 | 1500 | 200 | 0.57 |
| 9 | 9600 | 11100 | 1500 | 600 | 0.79 |
| 10 | 10800 | 12300 | 1500 | 300 | 0.81 |
| 11 | 12000 | 13500 | 1500 | 300 | 0.85 |
| 12 | 13200 | 14700 | 1500 | 300 | 0.88 |
| 13 | 14400 | 15900 | 1500 | 300 | 0.86 |
| 14 | 15600 | 17100 | 1500 | 300 | 0.87 |
| 15 | 16800 | 18300 | 1500 | 300 | 0.87 |
| 16 | 18000 | 19500 | 1500 | 300 | 0.85 |
| 17 | 19148 | 20647 | 1499 | 352 | 0.81 |

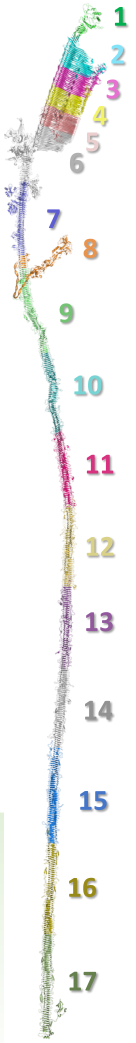


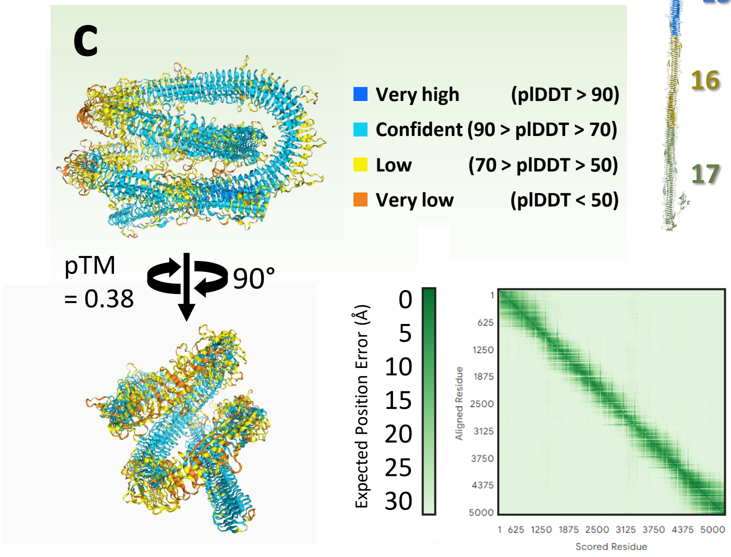


**Fig. S1. Modeling the predicted structure of giant symbiosis proteins**. Full protein structural predictions of **(a)** Cag_663 and **(b)** Cag_665 using AlphaFold3 to determine 1500 aa fragments with 300 aa overlap which could then be assembled in PyMOL to construct the final assembly. **(c)** Inputs of 5000 aa segments, such as Cag_663:27001-32000 yielded tertiary bundling or hairball structures with low confidence values, indicating artifacts. For final assembly amino acids 14712, 19443, and 19960 of Cag_663, as well as amino acids 5990 and 10614 of Cag_665 had to be manually rotated to prevent erroneous folding back onto the predicted structure. Fragments used to construct each protein are displayed on following pages, as Cag_663, Cag_664, and Cag_665, followed by Cag_2037 with Cag_2038.

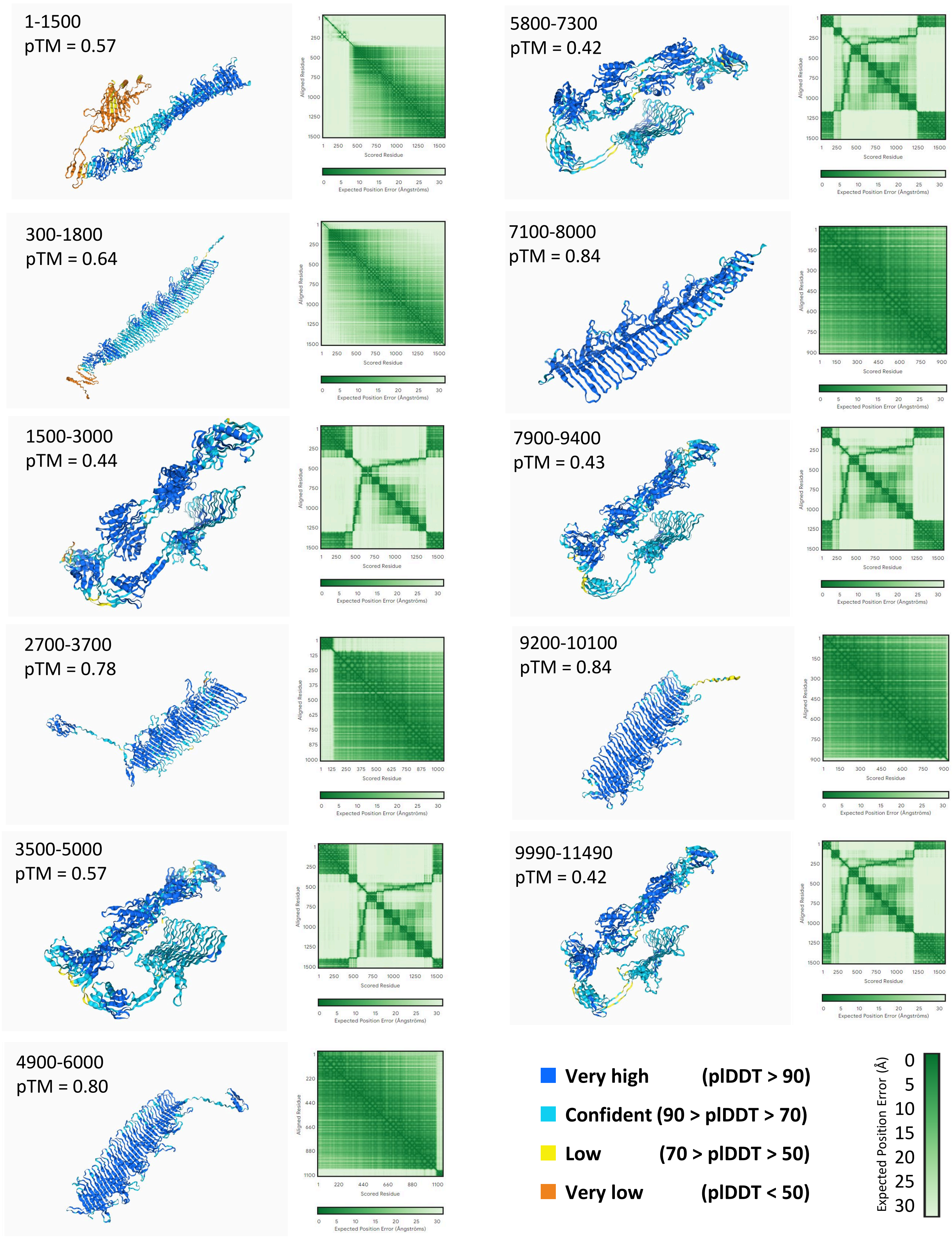


**Fig. S1. Continued. Fragments used to assemble Cag_663.**

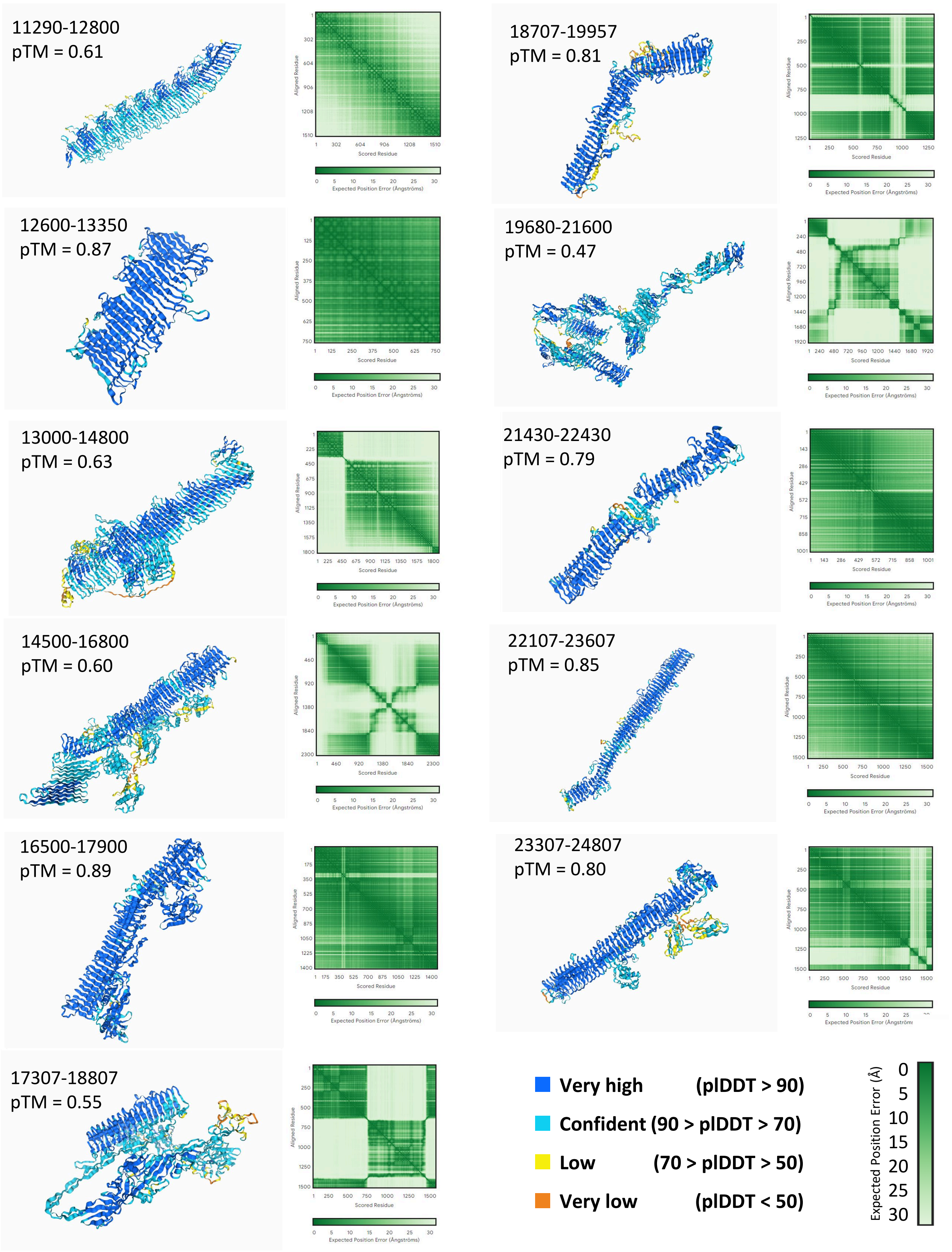


**Fig. S1. Continued. Fragments used to assemble Cag_663.**

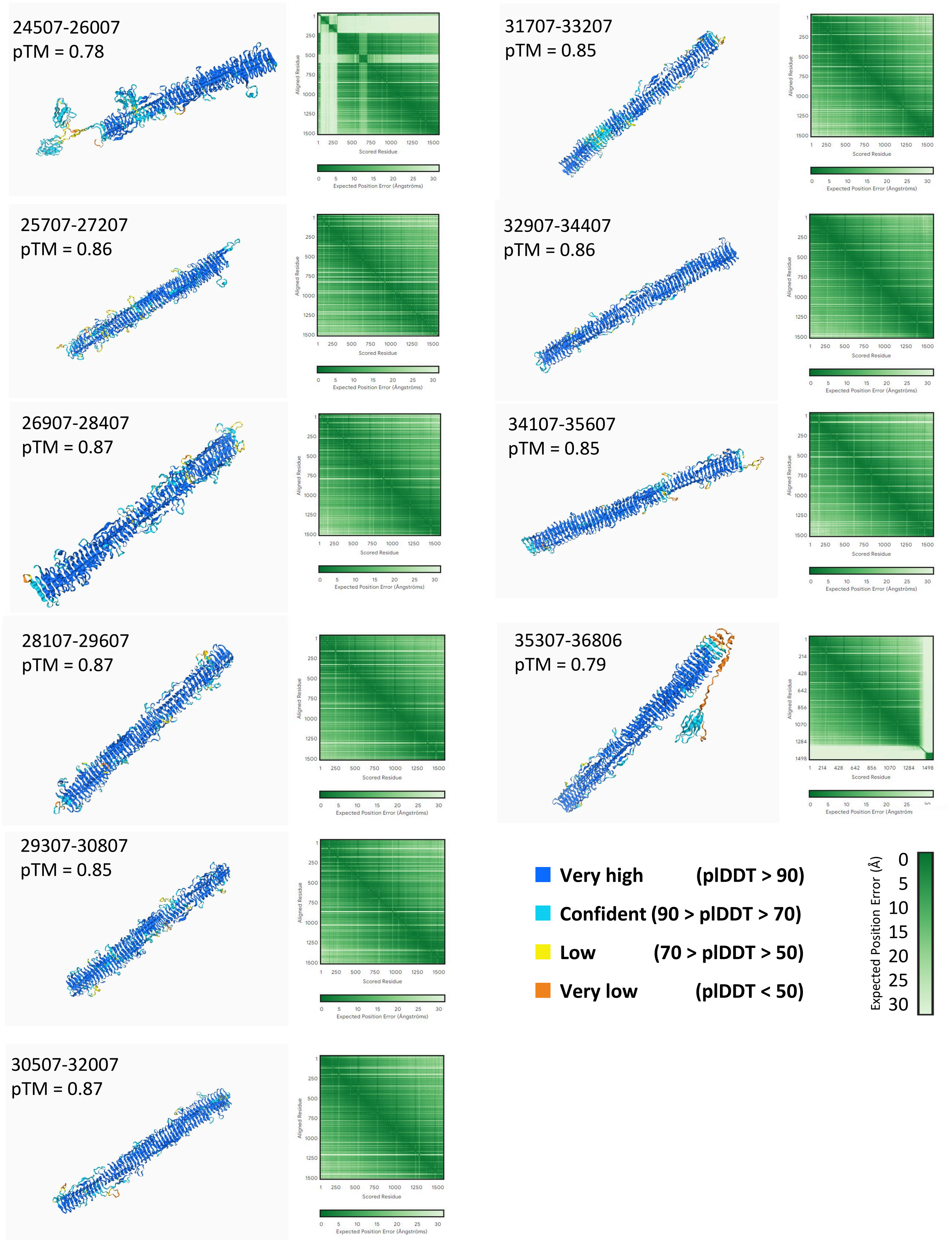


**Fig. S1. Continued. Fragments used to assemble Cag_663.**

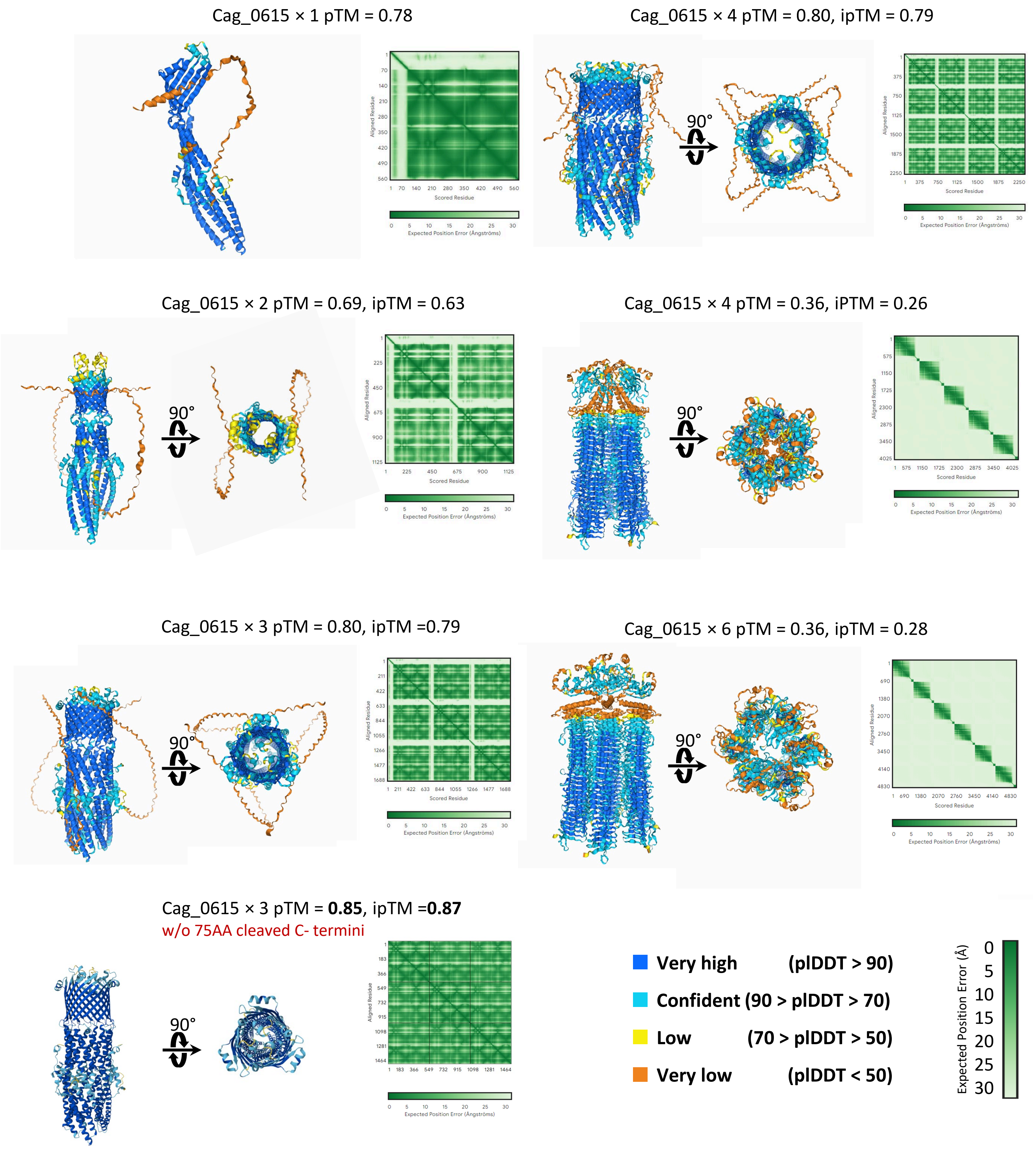


**Fig. S1. Continued. tolC Cag_664 oligomer predictions, suggesting 3× or 4× as highest likelihood, where 3 × with cleaved N- termini is typical, and strongest supported.**

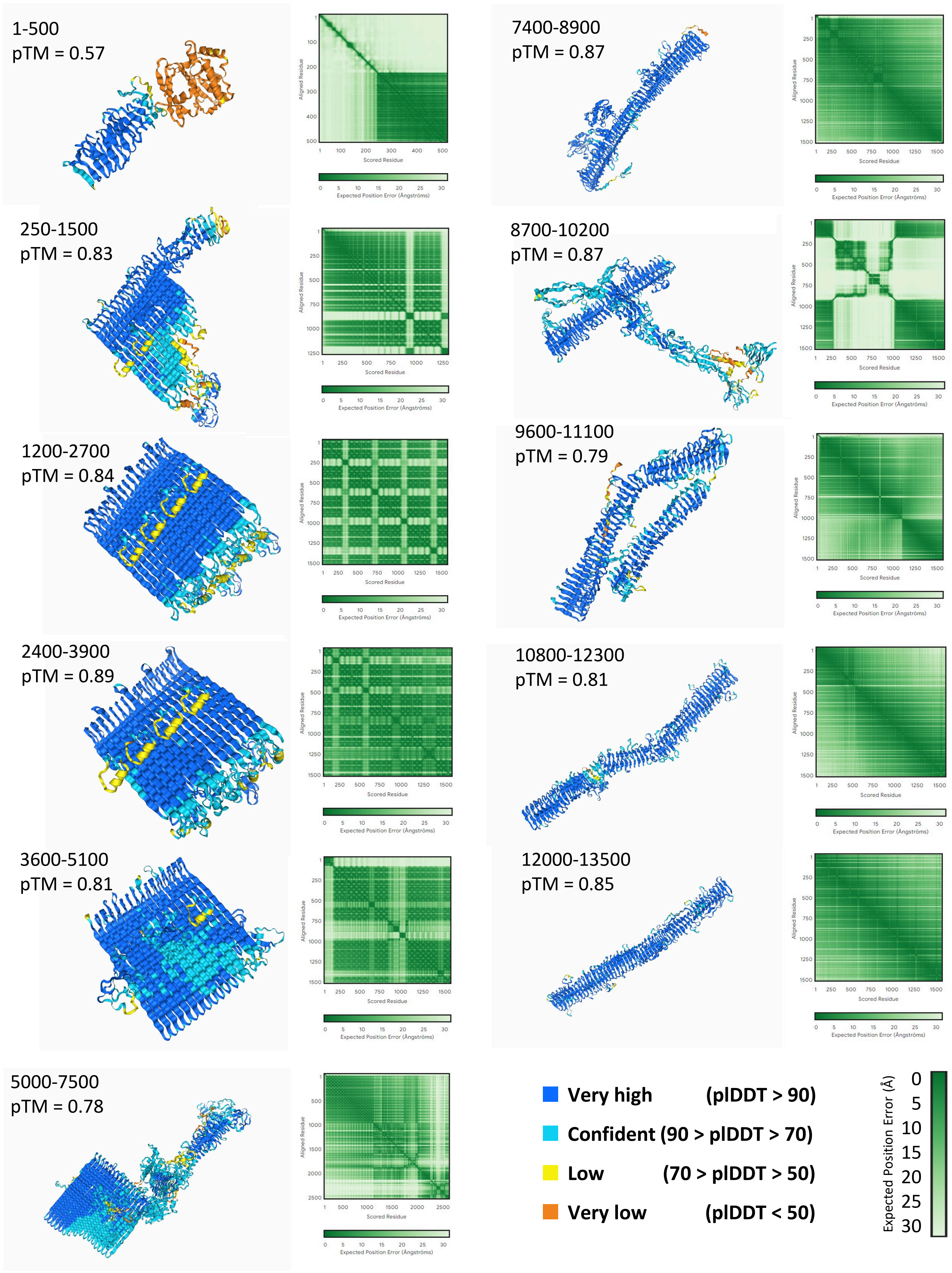


**Fig. S1. Continued. Fragments used to assemble Cag_665.**

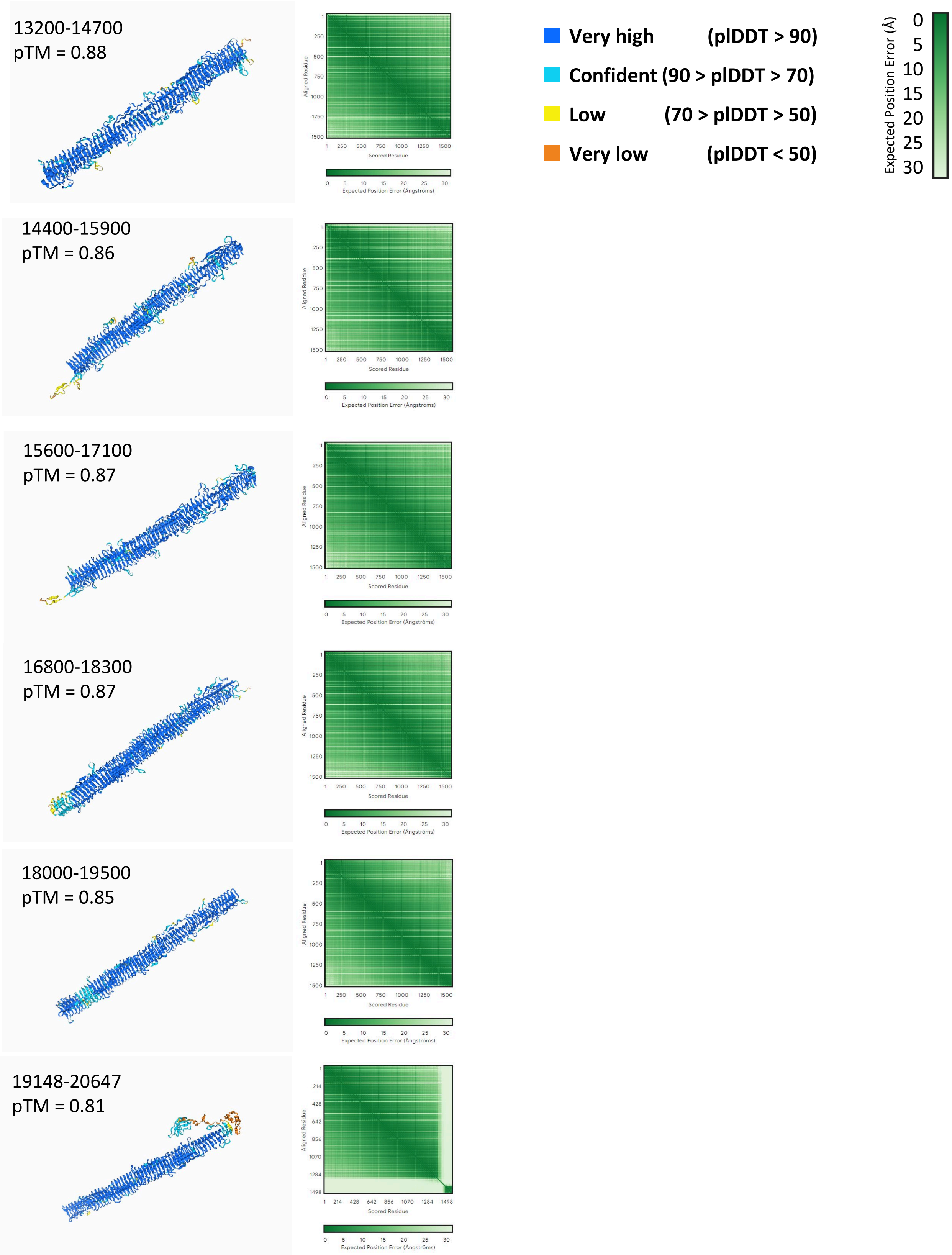


**Fig. S1. Continued. Fragments used to assemble Cag_665.**

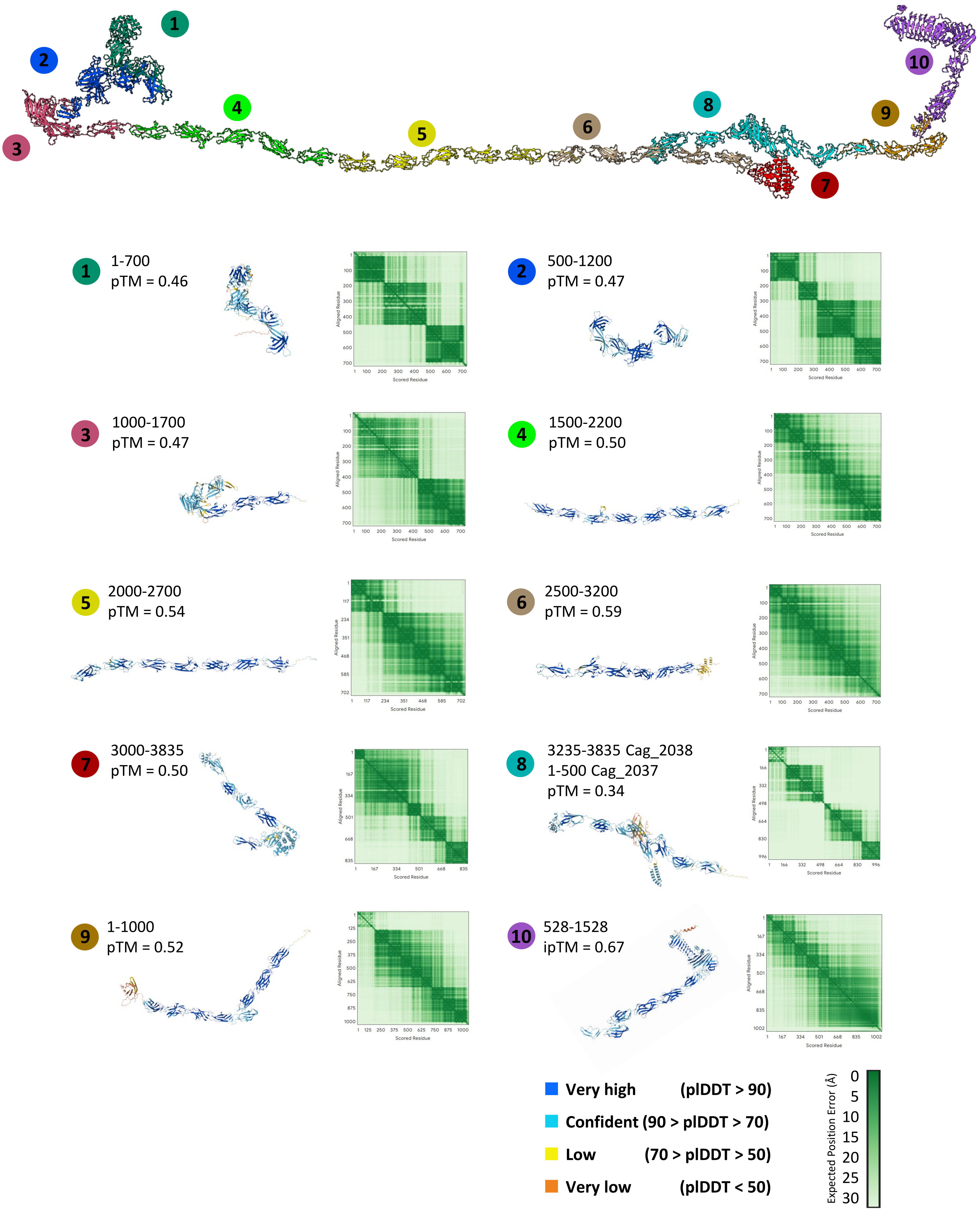


**Fig. S1. Continued.** Individual regions of Cag_2037 and Cag-2038 cellulosome proteins predicted in AlphaFold3 presented along with alignment (PyMOL).

| Ca²⁺ added | ipTM | pTM |
|---|---|---|
| $Ca^{2+}$ × 0 | - | 0.38 |
| $Ca^{2+}$ × 1 | 0.53 | 0.40 |
| $Ca^{2+}$ × 2 | 0.49 | 0.39 |
| $Ca^{2+}$ × 3 | 0.57 | 0.40 |
| $Ca^{2+}$ × 4 | 0.61 | 0.41 |
| **$Ca^{2+}$ × 5** | **0.64** | **0.41** |
| $Ca^{2+}$ × 6 | 0.50 | 0.34 |
| $Ca^{2+}$ × 12 | 0.45 | 0.36 |

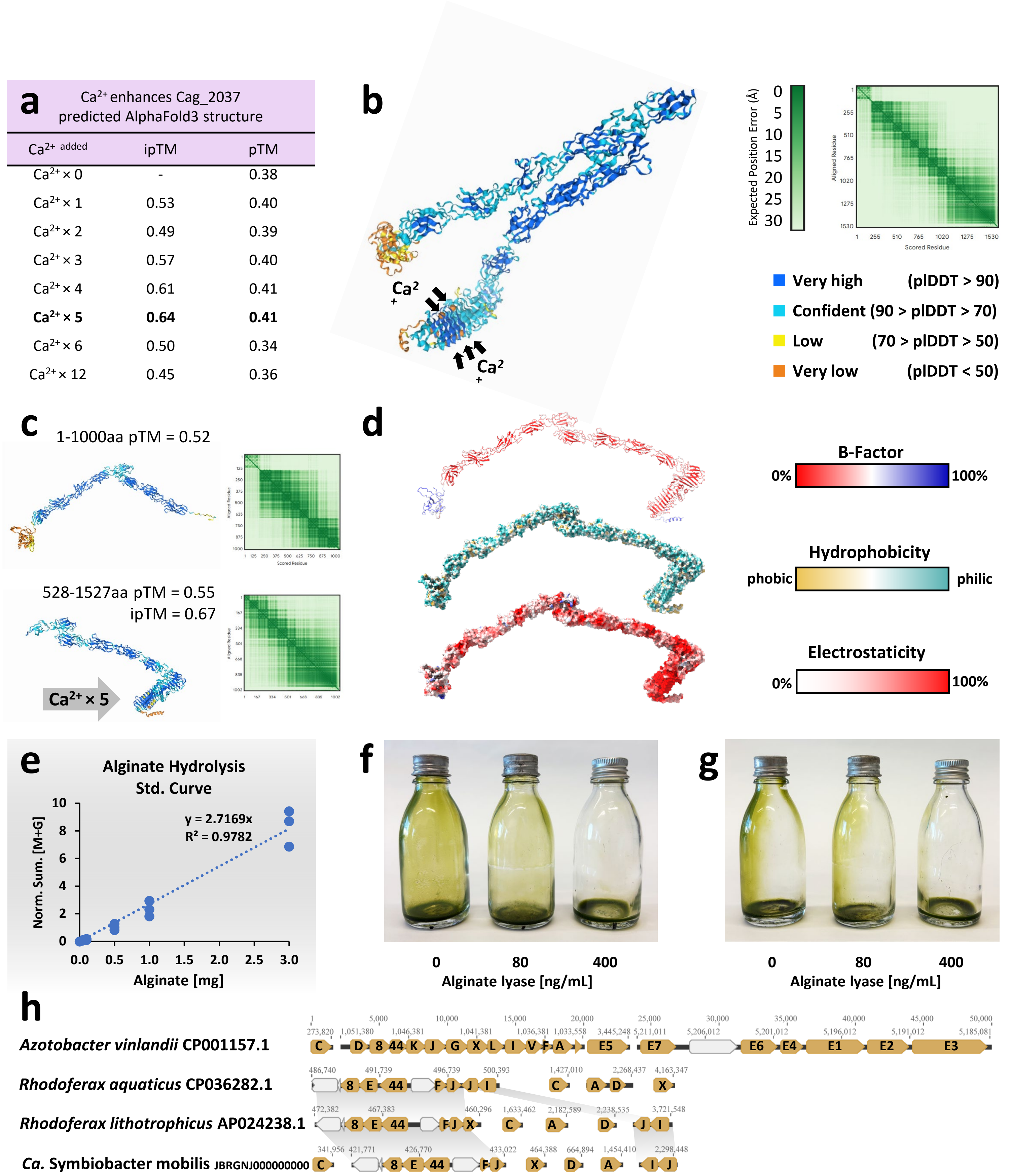


**Fig. S2. Alginate lyase supporting information. (a)** Cag_2037 structural predictions were strengthened by addition of five Calcium cations, **(b)** which imbed into the RTX domain. **(c)** Partial predicted structures elongated Big domains, with **(d)** concatenated complete Cag_2037 structure revealing a stable, hydrophilic structure that has a flexible N- terminal cohesion domain and an electrostatic C- terminal RTX domain. **(e)** Precise amounts of alginate was hydrolyzed and analyzed via MS to achieve a standard curve of detectable levels, for comparisons to measurements within consortia and axenic *Chl. chlorochromatii* biomass. **(f)** The macroscopic observation of diminished '*C. aggregatum*' biofilm production from exogenous alginate lyase additions, **(g)** as seen from the side the reduction of phototactic ability is a result of disintegrated consortia. **(h)** Alginate synthesis genes clustered within the genome of *Gammaproteobacteria* representative *Azotobacter vinlandii,* while a new gene cluster arrangement was detected for *Betaproteobacteria Ca.* Symbiobacter mobilis, and close relatives within *Rhodoferax*.

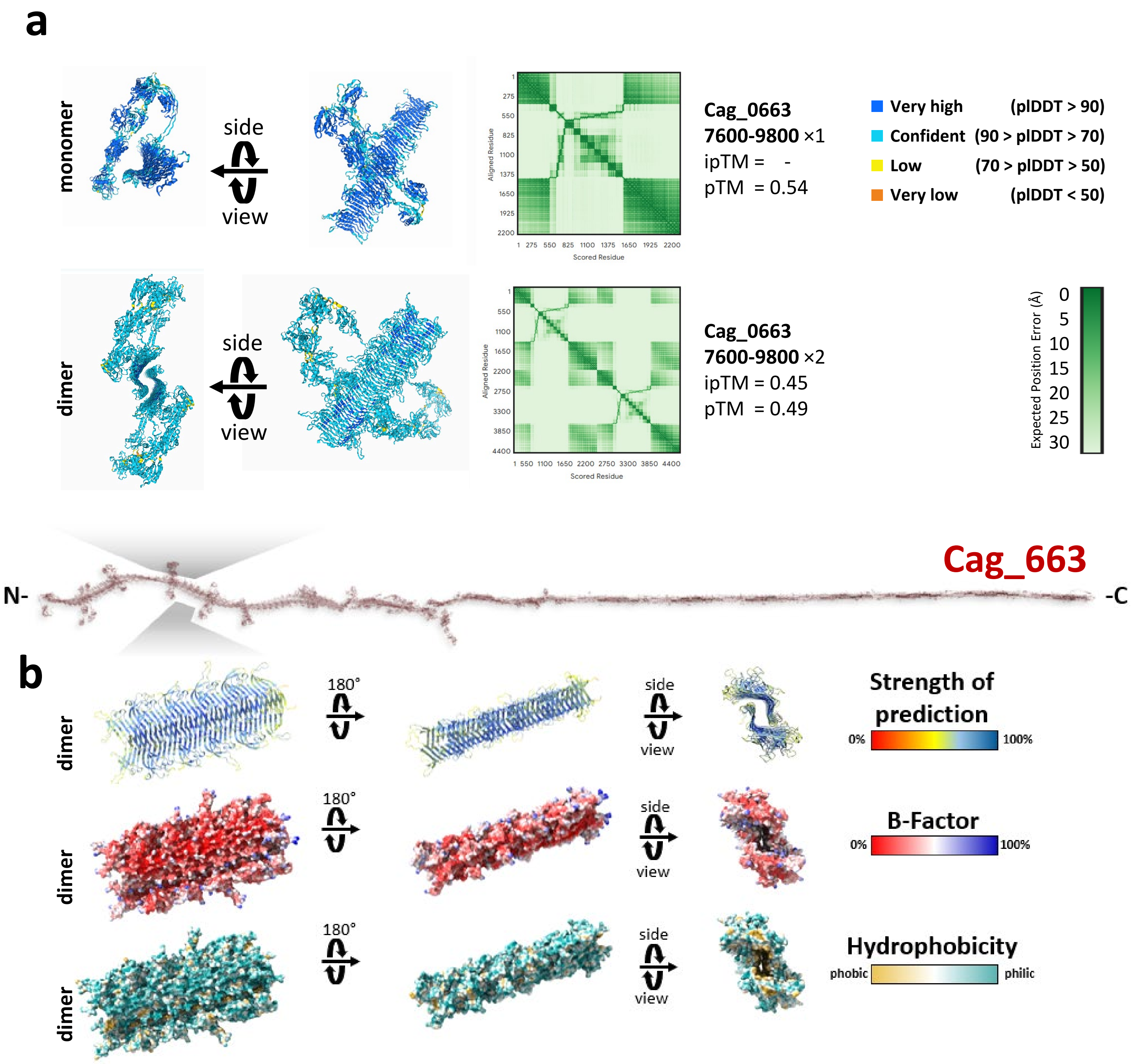


**Fig. S3. Modeling of protein-protein interaction for the β-plate region of Cag_663. (a)** The largest section possible (2,200 aa) of the twisted β-sheet with hydrophobic cleft and one extended filamentous hemagglutinin arm encoded by Cag_663 was modelled as a monomer (upper part) or dimer (lower part). Two β-sheets can dimerize at their hydrophobic clefts, where each co-occurring capsid-like/HK97-fold motifs remain exposed to the outside. **(b)** A dimeric region which excludes the extended capsid-like/HK97-fold motifs also hides the hydrophobic cleft, exposing only hydrophilic regions to the outside.

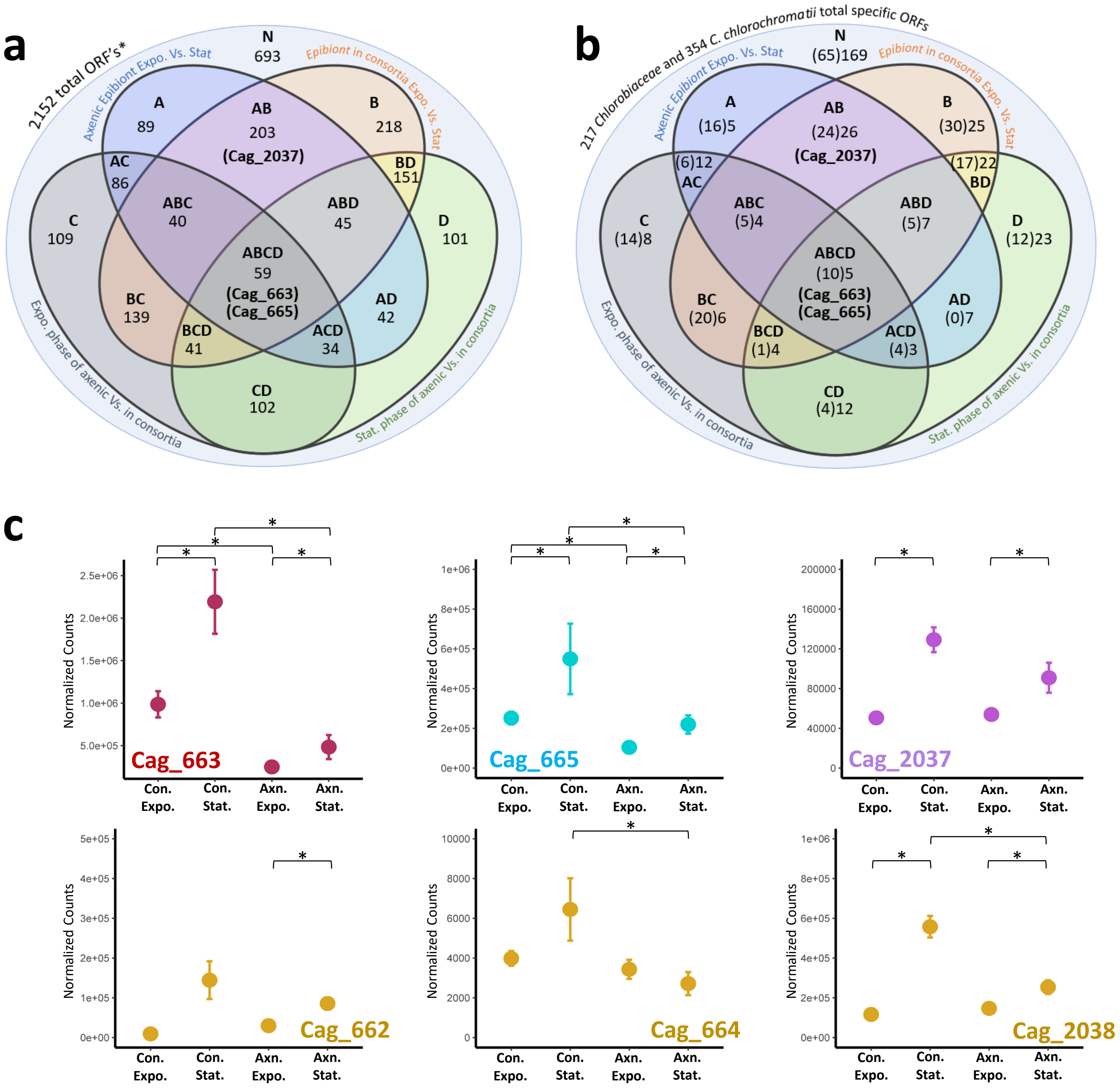


**Fig. S4. Differentially expressed *Chl. chlorochromatii* genes.** Transcriptome sequenced and normalized from axenic and symbiotic cultures during exponential and stationary phases (4 conditions) are compared, utilizing means of three biological replicates ranked by p-value, sorted by fold change if p<0.05. Contrasting [A] Axenic *Chl. chlorochromatii* in stationary vs. exponential phase, [B] Symbiotic *Chl. chlorochromatii* in stationary vs. exponential phase, [C] Differences of symbiotic vs. axenic *Chl. chlorochromatii* both in the exponential phase, or [D] Differences of symbiotic vs. axenic *Chl. chlorochromatii* both in the stationary phase, provided 16 conditions in which genes can be differentially expressed. **(a)** Total genes differentially expressed in all conditions for each intersection are listed (without differentiating if up or down regulated), where specific genes of each group are provided in the corresponding supplemental Table S7. **(b)** *Chl. chlorochromatii* genes that are specific to the *Chlorobiaceae* family in brackets, or unique to this species that are differentially expressed in all conditions for each intersection are listed (without differentiating if up or down regulated). Cag_663, Cag_665 and Cag_2037 are in Groups ABCD, ABCD, and AB, respectively. *While *Chl. chlorochromatii* has 2,096 protein coding genes, all 2,152 ORF's are included in this analysis. **(c)** Significant differences of select normalized counts of symbiosis genes. Axenic, Axn.; Consortia, Con.; Exponential, Expo.; Stationary, Stat.

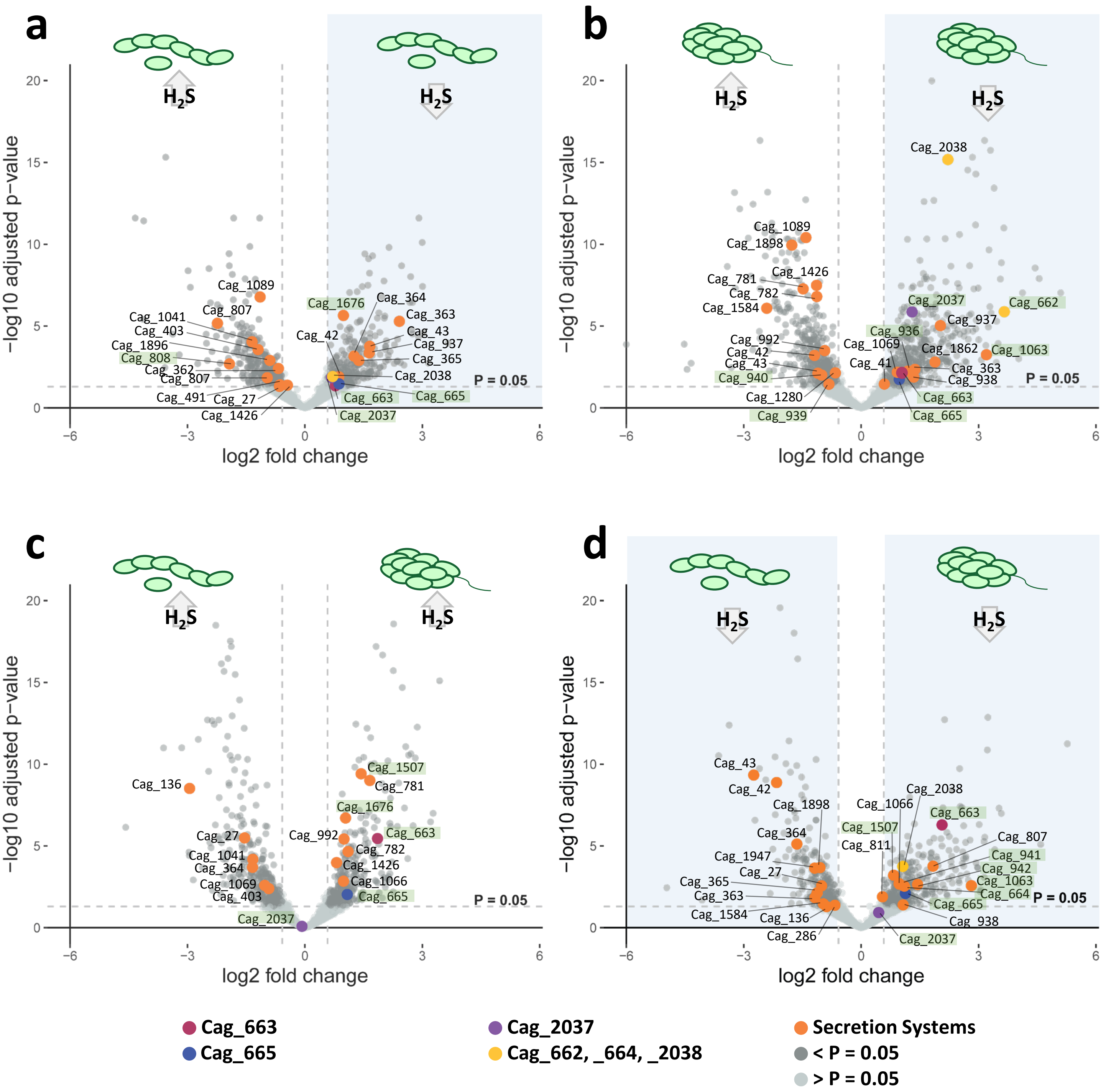


**Fig. S5. Detailed analysis of differential expression patterns in *Chl. chlorochromatii* across different growth conditions and symbiotic states.** Volcano plots of pairwise comparisons between exponential and stationary phase for the same **(a)** axenic or **(b)** symbiotic state (upper panels) are contrasted to expression patterns found between axenic or symbiotic state in the same **(c)** exponential or **(d)** stationary growth phase (lower panels). The three symbiosis genes, associated genes, and secretion systems are colored as seen in Figure 2,3,S8 and elsewhere. Genes highlighted green are unique to *Chl. chlorochromatii* among *Chlorobiota*.

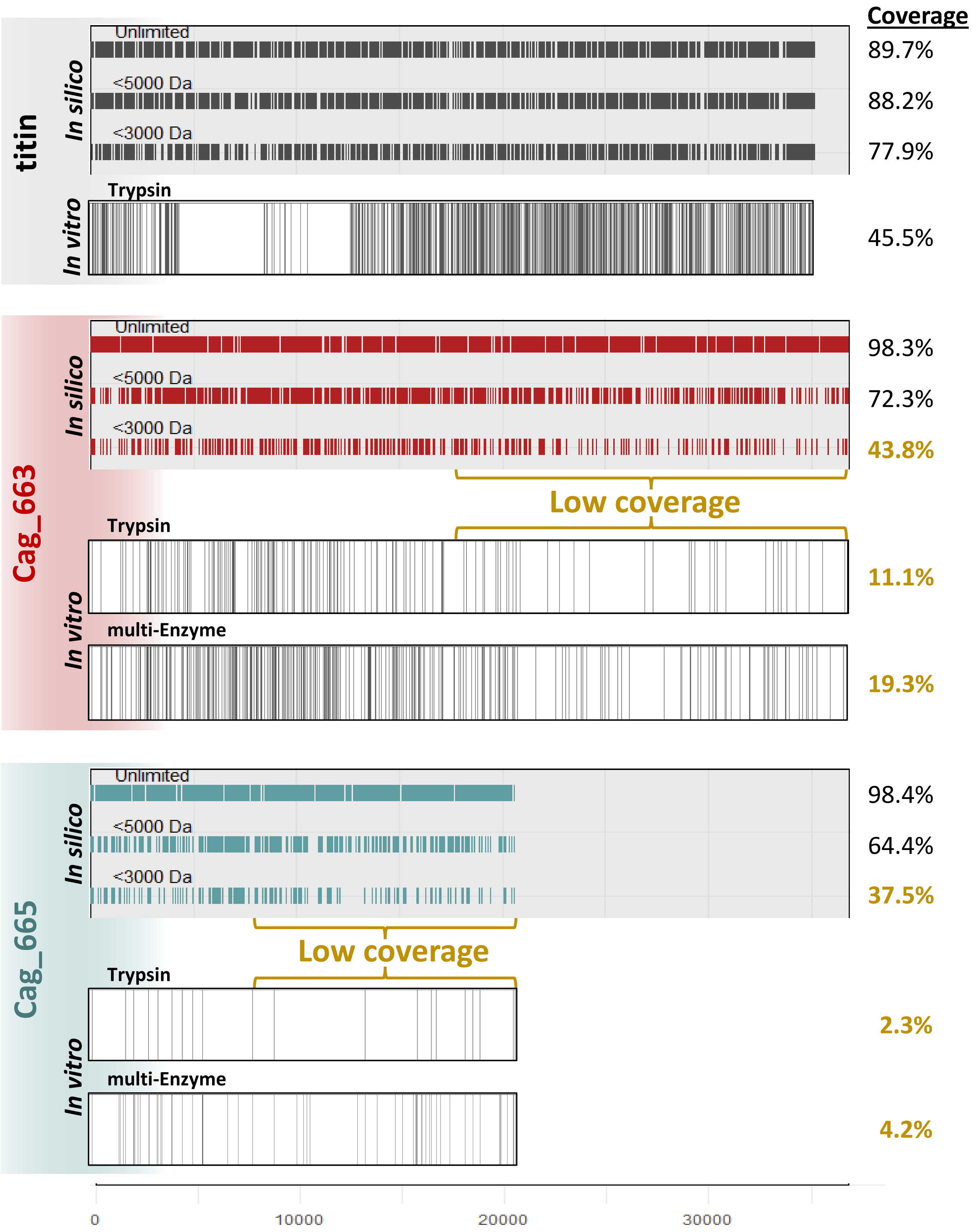


**Fig S6. Trypsin enzymatic digestion of titin as compared to giant virulence-like symbiosis factors.** *In silico* peptides degraded by Trypsin are generated in Expasy [77], allowing 0 missing cleavages, and filtering peptides >500 Da, showing those below certain thresholds. Masses of peptides calculated as monoisotopic [M+H]+. Detectable peptides then mapped back to entire protein sequence N- to C- termini in ggplot (R). https://web.expasy.org/peptide_mass/ . Below each *in silico* trypsin digest is pooled *in vitro* trypsin-only or multi-enzyme digests of *Mus* or *Chlorobium* biomass, revealing similarities between *in silico* and *in vitro* peptides for all 3. While titin is degradable by trypsin, these results further suggest both symbiosis proteins are indeed present, but are resistant to complete trypsin digest. Amino acid position, aa.

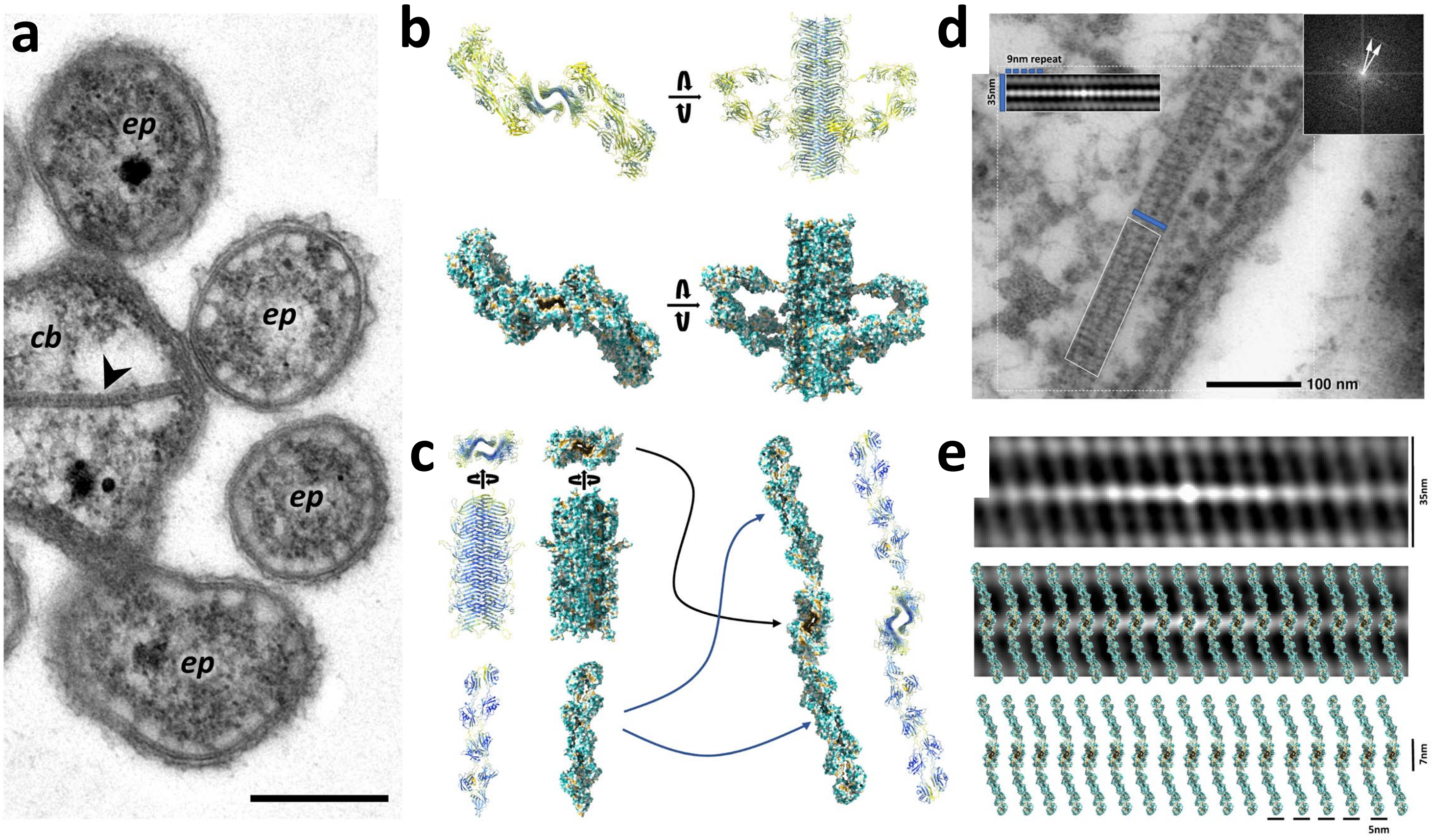


**Fig. S7. Chemotaxis array hypothesis of Cag_663 active domain. (a)** GA fixed '*C. aggregatum*' consortia reveal '*Ca.* S. mobilis' central bacterium to contain paracrystalline (CBC) structures (arrow) as previously documented [15], which are similar in appearance to (Aero)chemotaxis arrays [73]. Scale bar, 0.4µm. **(b)** As in Fig. S2, regions of Cag_663 hydrophobicity will form dimers, where **(c)** flexible BIg domain arms have the possibility to become extended. **(d)** Looking at density or paracrystalline structures, auto-correlation of CBC reveals oblique orientation of the subunits, in which **(e)** extended and stacked Cag_663 active domains potentially fit.
